# Multi-Sample and Multi-Group Spatial Colocalization Analysis Using PANORAMIC

**DOI:** 10.1101/2025.09.18.677135

**Authors:** Jacob Chang, Almudena Espín Pérez, Perla Molina, Rohit Khurana, Weiruo Zhang, Lu Tian, Sylvia Plevritis

## Abstract

**Motivation:** Spatial omics studies compare cell-cell organization across samples, but most methods model between-sample variability while treating sample-level spatial estimates as error-free. Overlooking within-sample uncertainty can distort inference in heterogeneous cohorts, motivating methods that explicitly quantify and propagate this uncertainty into cohort-level analyses.

**Results:** We present PANORAMIC, a hierarchical framework for spatial colocalization analysis that uses edge-corrected neighborhood enrichment to estimate local cell-type colocalization, spatial bootstrapping to quantify within-sample uncertainty, and multilevel random-effects meta-analysis to propagate this uncertainty across samples, patients, and conditions. In simulations, PANORAMIC improved recovery of within-sample uncertainty and between-sample heterogeneity relative to naive estimators across diverse spatial settings and progressive data degradation. Applied to a colorectal cancer tissue microarray profiled by multiplexed immunofluorescence imaging, PANORAMIC identified stronger B- and T-cell colocalization in tumors with Crohn’s-like reaction than in tumors with diffuse inflammatory infiltration, together with tighter higher-order immune organization consistent with immune aggregates. These results show that propagating within-sample spatial uncertainty can improve cohort-level inference in spatial omics studies.

**Availability and Implementation:** PANORAMIC is released as an open-source R package at https://github.com/plevritis-lab/panoramic.

## Introduction

Spatially resolved single-cell technologies are transforming the study of tissue architecture by providing molecular, phenotypic, and positional measurements for thousands of cells within intact tissues [5, 3, 26, 4, 18, 7, 12, 9, 11]. These technologies make it possible to quantify not only which cell types are present in a tissue, but also how they are spatially organized. A central analysis task in spatial omics has been to measure spatial colocalization between cell types, which can serve as a proxy for cellular interaction, a distinct microenvironment, and coordinated tissue organization [20, 11]. For example, the spatial organization of cytotoxic T cells relative to tumor cells is a clinically relevant feature of the tumor immune microenvironment [6], while organized lymphoid aggregates involving B cells, T cells, and antigen-presenting cells are characteristic of tertiary lymphoid structures that have been linked to favorable prognosis and immunotherapy response [19, 10]. These and related findings demonstrate that cell-type spatial relationships encode biologically and clinically meaningful aspects of tissue organization.

Despite the value of colocalization analysis, rigorous statistical comparison of colocalization across samples remains challenging. Each tissue sample can be regarded as a distinct realization of a high-dimensional stochastic spatial process, with thousands of spatially correlated cells nested within samples, samples nested within patients, and patients grouped by clinical condition or phenotype. Treating cells as independent experimental units can lead to pseudoreplication and inflated evidence in single-cell studies [29, 23]. In spatial omics, this challenge is compounded by local spatial heterogeneity: different regions of the same tissue section may exhibit distinct cellular neighborhoods, densities, and spatial association patterns. Thus, differential colocalization analysis must account for multiple layers of variability, including within-sample spatial uncertainty, between-sample biological and technical heterogeneity, and repeated sampling from the same patient.

Several recent methods address important components of this problem. SpicyR summarizes pairwise cell-type localization at the image level, and uses regression or mixed-effects models to test for differential spatial relationships, including settings with multiple images per subject and heterogeneous cell counts [2]. SpaceANOVA represents cell-type co-occurrence as a distance-indexed spatial interaction function and applies functional ANOVA to compare groups over spatial scales [22]. Other spatial association statistics, such as the colocation quotient and related point-pattern summaries, have also been used to quantify pairwise cell-type spatial relationships in tissue [14, 28, 27]. However, to our knowledge, existing approaches do not explicitly combine spatial bootstrap estimates of within-sample uncertainty with multilevel random-effects pooling across samples and patients. This distinction is consequential for clinical spatial omics studies, which often involve small cohorts, substantial biological heterogeneity, and multiple tissue regions or cores per patient.

We present PANORAMIC (Pooled ANalysis Of VaRiance-Aware Modeling and Inference of Colocalization), a statistical framework for pooled inference of cell-type colocalization across spatial omics samples. PANORAMIC estimates sample-specific colocalization statistics and their uncertainty using spatial bootstrapping, which preserves local spatial dependence while quantifying within-sample variability [15, 16]. These sample-level estimates and variances are then propagated through a multilevel random-effects meta-analysis fit by restricted maximum likelihood (REML), allowing PANORAMIC to pool colocalization estimates across samples while modeling heterogeneity across patients and repeated samples and estimating differences between biological conditions [25, 13, 17, 1]. This yields a hierarchical approach that respects both spatial dependence within tissue samples and heterogeneity across samples, patients, and clinical groups. Although we use edge-corrected neighborhood enrichment as the default colocalization statistic, PANORAMIC is statistic-agnostic: any sample-level spatial statistic for which bootstrap replicate estimates can be computed, such as Ripley’s *K*- and *L*-functions, may be propagated through the same hierarchical framework.

To evaluate PANORAMIC, we apply it to simulated spatial point-process data with known generative structure and estimands, benchmark its uncertainty calibration, heterogeneity recovery, and robustness to sample degradation. We then demonstrate its utility on multiplexed immunofluorescence (mIF) imaging datasets from colorectal cancer (CRC) and head and neck squamous cell carcinoma (HNSCC) tissue microarrays (TMAs). In CRC, Schürch et al. used mIF imaging to identify conserved cellular neighborhoods at the invasive tumor front and showed that Crohn’s-like reaction (CLR) tumors, compared with diffuse inflammatory infiltration (DII) tumors, exhibit organized immune hubs associated with favorable prognosis [21]. When applied to this dataset, PANORAMIC systematically tests directional cell-type pair colocalizations while propagating spatial uncertainty. We further apply PANORAMIC to a HNSCC TMA from Haist et al., who used spatial profiling of primary tumors and paired tumor-draining lymph nodes to study lymph node colonization and metastatic immune remodeling [8]. Across simulations and real datasets, PANORAMIC provides a practical framework for pooling spatial information within a biological condition, comparing spatial organization across conditions, and ultimately summarizing higher-order tissue organization through networks of differential colocalization.

## Materials and Methods

### Overview of the PANORAMIC workflow

PANORAMIC performs pooled inference for cell-type colocalization across spatial omics samples. For each sample, cell-type pair, and specified spatial radius, PANORAMIC first computes an edge-corrected neighborhood enrichment statistic that measures whether one cell type is enriched around another cell type relative to its sample-wide cell density. PANORAMIC then estimates within-sample uncertainty using a block-based spatial bootstrap that resamples local tissue regions rather than individual cells. The resulting sample-level estimates and bootstrap variances are used as inputs to multilevel random-effects meta-analysis models, which pool colocalization effects and test for differences between groups while modeling heterogeneity across patients and repeated tissue samples (**Figure 1**).

**Figure 1.**
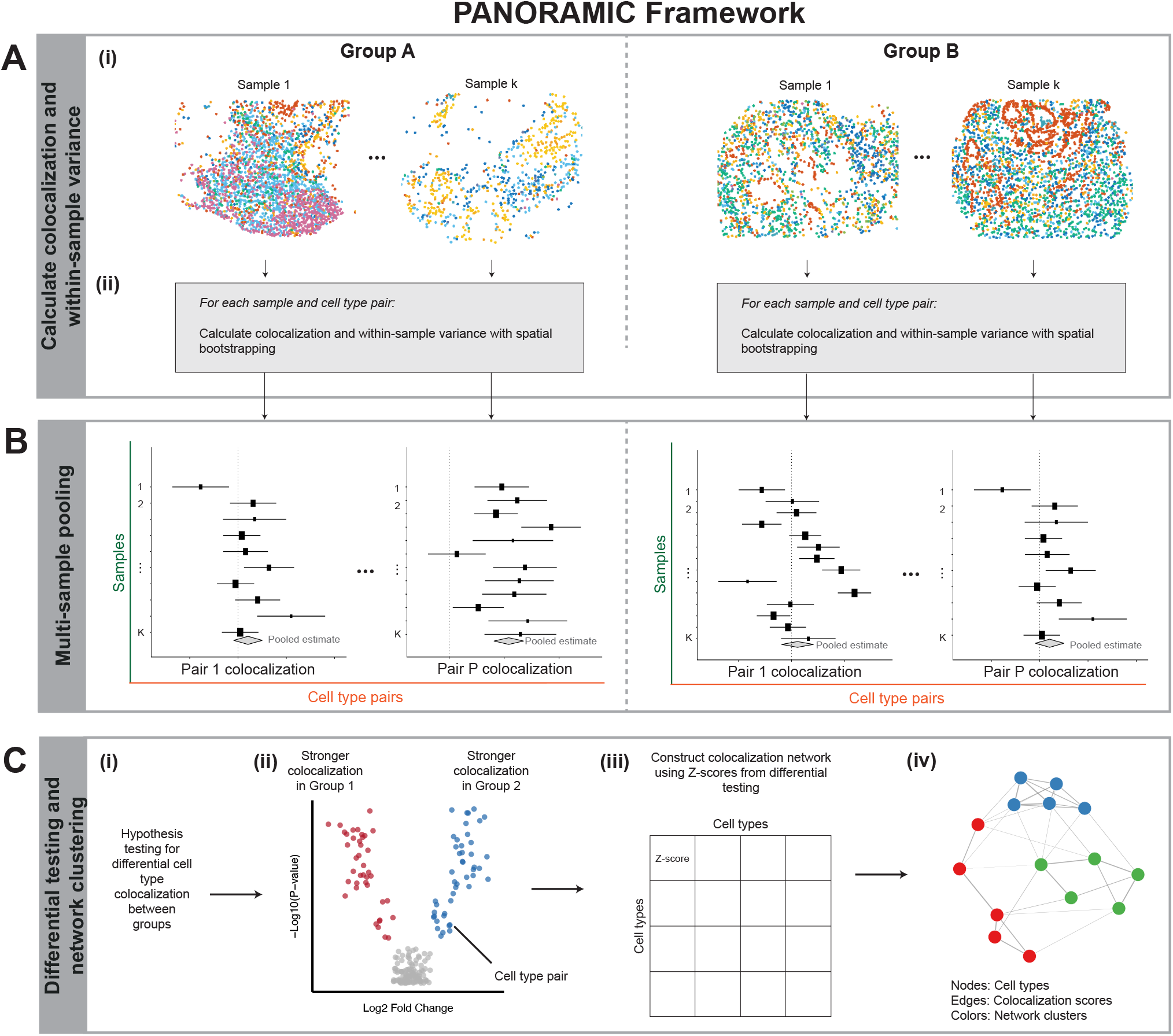
Overview of the PANORAMIC framework. **(A)** Calculation of colocalization and within-sample variance for each sample. (i) Spatial omics input data (cell type annotations and coordinates) formatted as a list of SpatialExperiment objects. (ii) For each sample *k* and cell type pair *p*, compute colocalization and estimate within-sample variance using spatial bootstrapping. **(B)** Multi-sample pooling. For each cell type pair *p*, pool samples within a group via multilevel random-effects meta-analysis. Forest plots show individual sample estimates with within-sample variance and pooled group means (gray diamonds). **(C)** Differential testing and network analysis. (i) Pooled means and standard errors are used to perform hypothesis testing for colocalization differences between groups A and B. (ii) Statistically significant cell type pairs visualized via volcano plot. Each point represents a cell type pair *p*. (iii) Colocalization matrix is constructed, populated by *Z*-scores between cell type pairs from the differential testing step. (iv) Network clustering is performed to identify higher-order colocalization subnetworks.

### Colocalization using edge-corrected neighborhood enrichment

For each sample, PANORAMIC quantifies pairwise cell-type colocalization using an edge-corrected neighborhood enrichment statistic. For an ordered pair *a* → *b*, PANORAMIC asks whether cells of type *a* are enriched in the local neighborhoods surrounding cells of type *b*, relative to the sample-wide density of type *a*.

For each anchor cell *i* of type *b*, we define a circular neighborhood of radius *r*. Let *n*_*i,a*_(*r*) denote the number of neighboring cells of type *a*, and let *n*_*i*,·_(*r*) denote the total number of neighboring cells of any type within radius *r*. Anchors with no neighboring cells within radius *r* are excluded.

The observed local proportion of type-*a* cells around anchor *i* is

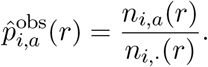

To account for tissue boundaries, we then compute the area of the radius-*r* disc around anchor *i* that lies inside the observed tissue region,

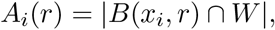

where *x*_*i*_ is the spatial coordinate of anchor *i, B*(*x*_*i*_, *r*) is the radius-*r* disc centered at *x*_*i*_, and *W* denotes the observed tissue window. Assuming homogeneity, the edge-corrected expected number of neighboring type-*a* cells around anchor *i* is

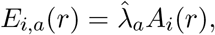

where 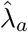 is the sample-wide density of cells of type *a*, calculated as the total number of cells of type *a* divided by the area of the observed tissue region. PANORAMIC then defines the anchor-level edge-corrected neighborhood enrichment score as

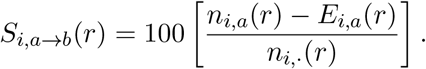

Positive values indicate that cells of type *a* are enriched within radius *r* of cells of type *b*, relative to the sample-wide density of type *a* and after accounting for boundary truncation. Negative values indicate depletion. The factor of 100 places the statistic on a percentage-point-like scale relative to the observed local neighborhood size.

For each sample *k*, radius *r*, and ordered pair *a* → *b*, PANORAMIC summarizes colocalization by averaging the anchor-level scores over all valid type-*b* anchors in the sample:

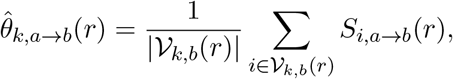

where 𝒱_*k,b*_(*r*) is the set of type-*b* anchor cells with at least one neighbor within radius *r*. This sample-level statistic is the effect size used in the downstream spatial bootstrap and multilevel meta-analysis. Thus, an effect size of 4 for *a* → *b* at radius *r* means that, on average, type-*a* cells are enriched by approximately four percentage points in the radius-*r* neighborhoods around type-*b* anchor cells, relative to the edge-corrected expected count for type-*a* cells in that sample.

Notably, this statistic is directional because neighborhoods are centered on one cell type and evaluated for enrichment of another. A high value of *a* → *b* means that type-*a* cells are enriched around type-*b* cells; it does not necessarily imply that type-*b* cells are enriched around type-*a* cells. Additional details on the edge-corrected neighborhood enrichment are provided in **Supplementary Methods S1**.

### Spatial bootstrapping and within-sample uncertainty

A single sample-level colocalization estimate can mask substantial within-sample spatial variability. In some tissues, spatial organization is relatively homogeneous and the estimate is stable across regions. In others, tissue compartmentalization, boundary effects, or localized cellular architecture can make the estimate sensitive to which regions of the tissue are sampled. PANORAMIC therefore treats each sample-level spatial statistic as an estimate with uncertainty, rather than as an error-free value.

To quantify within-sample uncertainty while preserving local spatial dependence, PANORAMIC uses an overlap-weighted block-based spatial bootstrap adapted from Loh-style spatial resampling [15, 16]. For each sample, the tissue window *W*_*k*_ is covered by a regular grid of square tiles. Each tile receives the overlap weight

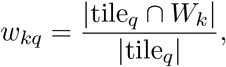

where *q* indexes tiles. Tiles which do not overlap with the tissue window are excluded, and the remaining tissue-overlapping tiles define the bootstrap sampling frame. Retained tiles are sampled with replacement using probabilities proportional to their overlap weights, so that tiles only partially contained in the observed tissue region are downweighted relative to tiles which fall completely within the observed tissue region (**Supplementary Methods S2**) in the resampling.

For each directional cell-type pair colocalization *f* in sample *k* at radius *r*, PANORAMIC first computes anchor-level neighborhood enrichment scores and assigns each valid anchor to a retained spatial tile. Within each tile, anchor-level scores are summarized to obtain tile-level means. A bootstrap replicate is then formed by sampling tissue-overlapping tiles with replacement and averaging the sampled tile-level summaries. This yields bootstrap replicate statistics

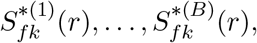

where *B* is the number of bootstrap replicates.

PANORAMIC estimates the sample-level effect as the mean of the bootstrap replicate statistics,

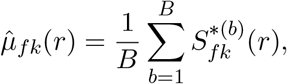

and the within-sample variance as

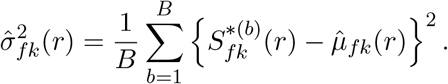

When *B* is big, 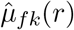 is expected to be close to the corresponding colocalization summary 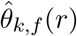 given earlier. These quantities define the sample-level estimate

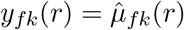

and the corresponding within-sample variance estimate

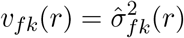

used in downstream multilevel meta-analysis. Consequently, PANORAMIC allows samples with unstable or spatially heterogeneous colocalization patterns to contribute less to pooled inference of a cell-type pair’s colocalization across samples and patients.

### Multilevel meta-analysis

To summarize colocalization within and between groups of samples, PANORAMIC fits feature-wise multilevel random-effects meta-analytic models using the metafor R package [25]. Here, *f* indexes a directional cell-type pair colocalization evaluated at a specified radius. For each cell-type pair *f* and sample *k*, the spatial bootstrap provides a sample-level effect estimate *y*_*fk*_ and a corresponding within-sample variance estimate *v*_*fk*_.

Let *c*(*k*) ∈ {1, · · ·, *C*} denote the biological group or condition of sample *k*, and let *m*(*k*) ∈ {1, · · ·, *M* } denote the patient from which sample *k* was obtained. For group-level estimation, PANORAMIC models

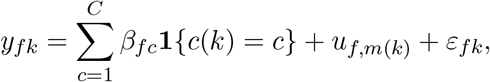

where *β*_*fc*_ is the expected colocalization level for feature *f* in group *c*,

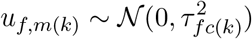

is a patient-level random effect capturing between-patient heterogeneity, and

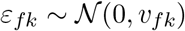

represents the estimated within-sample uncertainty from the spatial bootstrap. When multiple samples or tissue cores are available from the same patient, PANORAMIC can include an additional sample-level random effect nested within patient:

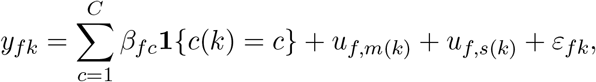

where *s*(*k*) denotes the sample or tissue region corresponding to patient *k* and

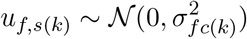

is the sample-level random effect capturing the additional between-sample heterogeneity within a patient. This hierarchical random-effects structure is selected according to the sampling design and the number of repeated samples per patient. This type of random-effects model can be fit using either moment- or likelihood-based method. The restricted maximum likelihood estimator (REML) is a common choice. In the end, estimators for group-level summary *β*_*fc*_ and variance components 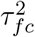 can be obtained along with their standard errors as the primary output of PANORAMIC.

For two-group comparisons, PANORAMIC estimates a contrast between the two group-specific pooled means,

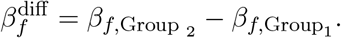

The Wald statistic is

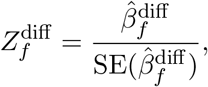

where 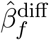 is the estimator of 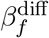 from PANORAMIC. Two-sided *p*-values are computed using the standard normal reference distribution, and Benjamini–Hochberg correction is applied across all tested cell-type pairs. This framework allows PANORAMIC to propagate spatial bootstrap uncertainty into group-level inference while also modeling heterogeneity across patients and repeated tissue samples. Additional details on the multilevel meta-analysis implementation are available in **Supplementary Methods S3**.

When sample-level clinical or demographic covariates are available and the number of biological replicates is sufficient, they can be included through meta-regression terms in the design matrix. However, as in standard meta-analysis, covariate-adjusted models require adequate sample size, and some variance components may not be identifiable for all covariate structures.

### Simulation studies

We evaluated PANORAMIC using two simulation studies designed to assess within-sample uncertainty estimation and multilevel inference under heterogeneous and progressively degraded sampling.

First, we evaluated whether the spatial bootstrap captures within-sample uncertainty in sample-level colocalization estimates. We simulated marked spatial point patterns in a 600 *µ*m × 600 *µ*m observation window under three spatial regimes: (i) spatially uniform cell types, (ii) cell types arranged with opposing spatial gradients, and (iii) clustered organization of cell types A–B (**Figure 2**). Simulations included five cell types, with overall cell densities scaled to match the average cell counts observed in the CRC TMA. For each simulated sample, PANORAMIC computed edge-corrected neighborhood enrichment at radius *r* = 25 *µ*m and estimated within-sample uncertainty using a spacial bootstrap with square tiles of side length 62.5 *µ*m. Bootstrap variance estimates were compared with a benchmark, which is the empirical variation of colocalization summaries across a large number of independently simulated samples. Additional simulation details are provided in **Supplementary Methods S4.1**.

**Figure 2.**
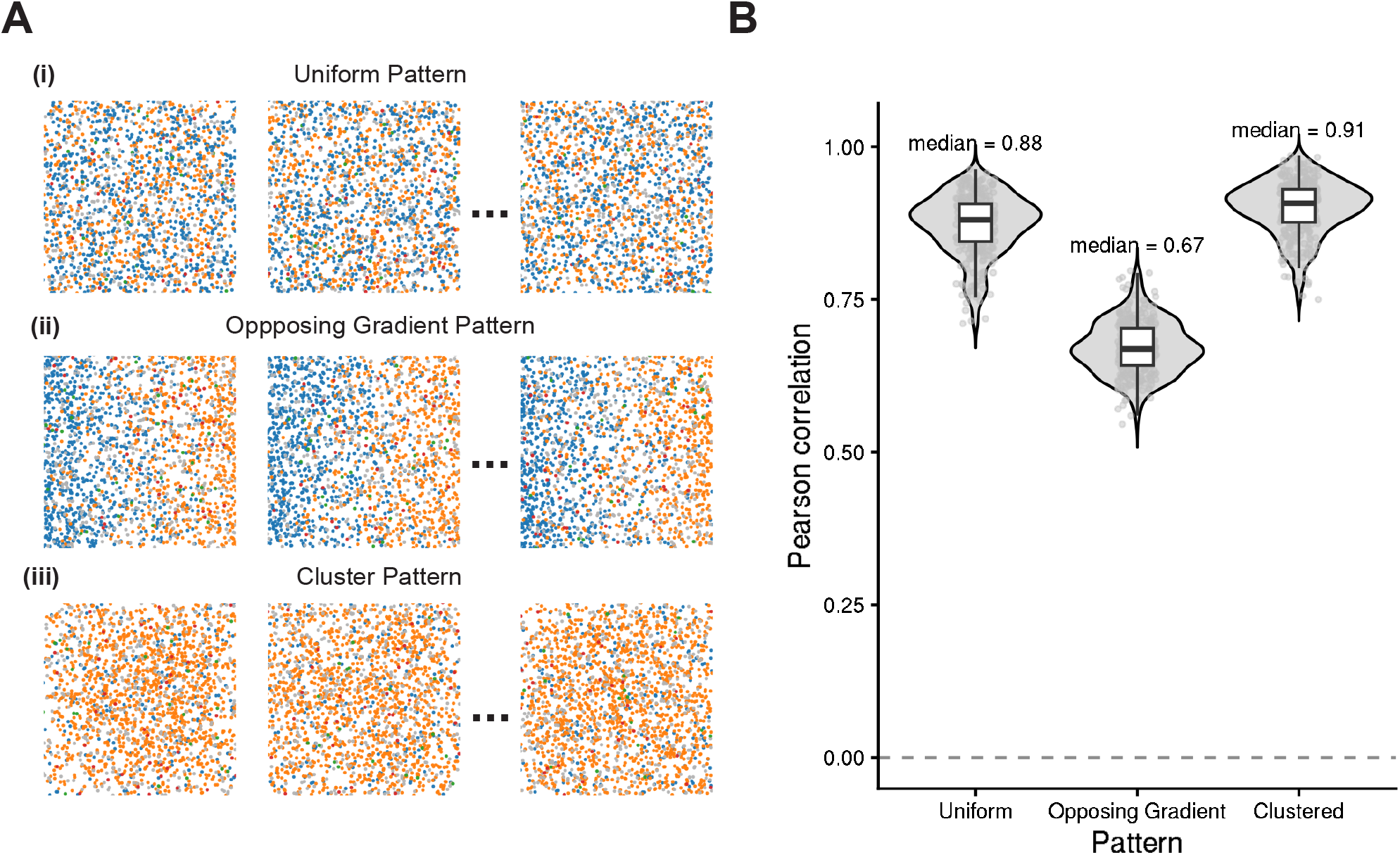
Validation of within-sample variance estimation via spatial bootstrapping. **(A)** Three spatial point process configurations were used for simulation: (i) uniform point pattern (complete spatial randomness), (ii) opposing gradient point pattern (spatial segregation with two cell types localized to opposite regions), and (iii) cluster point pattern (offspring occurring around parent cells). Cell types A, B, C are abundant; Rare 1 and Rare 2 represent low-frequency populations, distributed randomly. **(B)** For each configuration, 300 independent realizations were generated to establish ground truth within-sample variance. Spatial bootstrapping (100 iterations) was applied to each individual realization, and Pearson correlation between bootstrap-derived variance and ground truth was computed across all cell type pairs (25 ordered pairs). Violin plots show distribution of correlations across 300 realizations, with median values labeled. Spatial bootstrapping shows high concordance for uniform (median r = 0.88) and cluster (r = 0.91) point patterns, indicating accurate variance recovery for these spatial configurations. Concordance is moderate for the opposing gradient point pattern (r = 0.67).

Second, we evaluated PANORAMIC in a nested sampling setting with both patient-level and sample-level heterogeneity, variable numbers of samples per patient, and progressive sample degradation. We simulated clustered A–B spatial point patterns in which patient-specific latent effects modulated cluster intensity, abundance, and spread, with additional variation among repeated samples from the same patient. This clean nested simulation was used to assess recovery of the pooled mean *µ* and patient-level heterogeneity *τ*^2^ across patient-count regimes. We then extended the same nested simulation framework to include ordered degradation scenarios ranging from a clean baseline to increasingly severe contamination and quality-related distortion, including cell thinning, directional cell-type bias, and additional sample-level heterogeneity. The multilevel random-effects meta-analysis based on sample-level estimates and bootstrap variances was conducted, and the performance was summarized by comparing PANORAMIC and naive estimates of *µ* and *τ*^2^ across patient-count and degradation regimes. Additional simulation details can be found in **Supplementary Methods S4.2**.

### CRC TMA data processing and analysis

The colorectal cancer (CRC) tissue microarray (TMA) data were obtained from Schürch et al. [21]. The authors used multiplexed immunofluorescence (mIF) imaging to profile tissue regions from the invasive front of advanced-stage CRC tumors, with patient-level annotations including Crohn’s-like reaction (CLR) and diffuse inflammatory infiltration (DII) immune patterns. The original study profiled 140 tissue regions from 35 patients using 56 protein markers and identified conserved cellular neighborhoods associated with antitumoral immune organization and clinical outcome [21] (**Supplementary Figure S8.1**).

For PANORAMIC analysis, we used the released single-cell table containing cell-type annotations and spatial coordinates. Spatial coordinates were converted from pixels to microns using the conversion factor reported for this dataset. We removed cells with undefined cell-type labels and excluded rare cell types that were absent from more than 20% of images, following the filtering strategy used for this dataset in the spicyR analysis [2]. The resulting processed dataset contained 12 retained cell types (**Supplementary Methods S5.1**). For CLR patients, one core annotated as containing lymphoid aggregate architecture was randomly selected when available; for DII patients, one diffuse-infiltration core was randomly selected (**Supplementary Methods S5.2**).

Pairwise colocalization was quantified using the edge-corrected neighborhood enrichment statistic at radius *r* = 25 *µ*m. Within-sample uncertainty was estimated using the spatial block bootstrap with *B* = 100 bootstrap replicates and square tiles of side length 62.5 *µ*m, corresponding to 2.5 × *r*. Cell-type pairs were evaluated only when the relevant cell types had at least five cells in a given sample. Sample-level estimates and bootstrap variances were then propagated into feature-wise multilevel random-effects meta-analysis. For the comparison of CLR and DII tumors, PANORAMIC used patient-level random effects, group-specific fixed-effect means, group-median variance flooring, and restricted maximum likelihood (REML) estimation (**Figure 4**).

### Differential network and spatial compactness analysis

Pairwise differential colocalization identifies specific cell-type relationships that differ between conditions, but it does not directly reveal whether multiple cell types reorganize as a coordinated spatial module. PANORAMIC therefore uses differential colocalization statistics to construct weighted cell-type colocalization networks. In the CRC colocalization network analysis, we visualized the CLR-enriched network, representing cell-type pairs with stronger colocalization in CLR than in DII. Nodes represent cell types, and edges represent significant differential colocalization relationships. For visualization, the network was rendered as an undirected graph and partitioned using Leiden clustering [24] (**Figure 5**).

To complement the graph visualization with a sample-level spatial summary, we performed a spatial compactness analysis on the cell-type set highlighted by the CLR-enriched network,

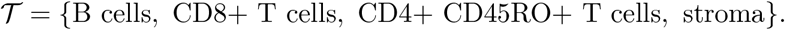

For each sample, we considered all cells whose labels belonged to 𝒯. For each such source cell, we computed the nearest-neighbor distance to each cell type in 𝒯, excluding the source cell itself when the source and target types were the same. Distances were capped at 25 *µ*m, matching the radius used in the primary colocalization analysis. These distances were then averaged across the target cell types in 𝒯, producing a per-cell within-set spacing summary.

To account for differences in overall cell density across samples, we normalized each per-cell within-set spacing summary by the nearest-neighbor distance from the same source cell to any other cell in the sample. The resulting relative spatial compactness values were averaged across cells to produce one spatial compactness score per sample. Smaller values indicate that the selected cell types are more tightly co-organized relative to the local cellular density baseline (**Supplementary Methods S5.5**).

Sample-level spatial compactness scores were compared between CLR and DII using Welch *t*-tests. The primary spatial compactness analysis used the mean relative spatial compactness score and tested whether CLR samples had lower relative compactness than DII samples. This analysis provides an independent sample-level summary of whether the cell types highlighted by the differential colocalization network are more tightly co-organized in CLR tumors.

### HNSCC TMA analysis

To assess whether PANORAMIC generalizes beyond colorectal cancer, we applied the workflow to a head and neck squamous cell carcinoma TMA cohort from Haist et al. [8]. Haist et al. used spatial proteomics, spatial transcriptomics, and experimental models to study lymph node colonization in head and neck cancer, reporting that nodal colonization is associated with tissue remodeling, interferon-*γ* signaling, enrichment of immunosuppressive myeloid cells and cancer-associated fibroblasts, and spatially organized fibroblast–myeloid niches linked to T cell dysfunction and Treg activation [8].

We used this cohort as a second real-data application to evaluate PANORAMIC in a setting with repeated tissue regions per patient and biologically distinct lymph node compartments. Specifically, we compared benign lymph node samples from node-negative patients with tumor-adjacent lymph node regions from patients with lymph node metastasis (**Supplementary Figure S9.1**). This comparison tests whether PANORAMIC can identify reproducible spatial reorganization associated with nodal metastatic involvement while accounting for multiple tissue regions from the same patient.

The HNSCC analysis used the same primary edge-corrected neighborhood enrichment statistic and spatial bootstrap design as the CRC analysis, with radius *r* = 25 *µ*m, *B* = 100 bootstrap replicates, square tiles of side length 62.5 *µ*m, and minimum cell-type counts of five cells per sample. Group comparisons were performed using feature-wise multilevel random-effects meta-analysis with patient-level or sample-level random effects selected according to the repeated-sampling structure.

## Results

### Intrasample simulations validate sample-level uncertainty estimation

We first evaluated whether PANORAMIC’s bootstrap variances track empirical within-sample variability. We simulated marked spatial point patterns under three regimes designed to represent distinct tissue organizations: spatially uniform cell types, opposing spatial gradients, and clustered A–B organization (**Figure 2A**).

For each regime, independent realizations were generated to estimate empirical variance for each ordered cell-type pair. PANORAMIC was then applied to individual realizations using *B* = 100 bootstrap replicates. Bootstrap-estimated variances were strongly correlated with empirical variances in the uniform and clustered settings, with median correlations of 0.88 and 0.91, respectively (**Figure 2B**). The opposing-gradient setting showed weaker, but still good, agreement, with a median correlation of 0.67.

These results indicate that the spatial bootstrap provides informative within-sample uncertainty estimates across multiple spatial regimes, with the strongest performance in settings where variability is driven by local spatial organization. The reduced agreement in the opposing-gradient setting suggests that broad tissue-scale compartmentalization is more difficult for local tile resampling to capture. This suggests that, when large-scale tissue domains dominate the spatial signal, users should consider pathologist-defined regions, tissue masks, or stratified analyses rather than relying on a single global tissue window.

### Intersample simulations show improved recovery of heterogeneity

We next evaluated PANORAMIC in hierarchical multi-sample simulations designed to assess recovery of the pooled mean colocalization and between-patient heterogeneity under progressive degradation. These simulations introduced patient-level variation, sample-level variation, variable numbers of samples per patient, and a degradation ladder ranging from baseline through mild, moderate, severe, and extreme contamination or quality distortion (**Figure 3B–C**).

**Figure 3.**
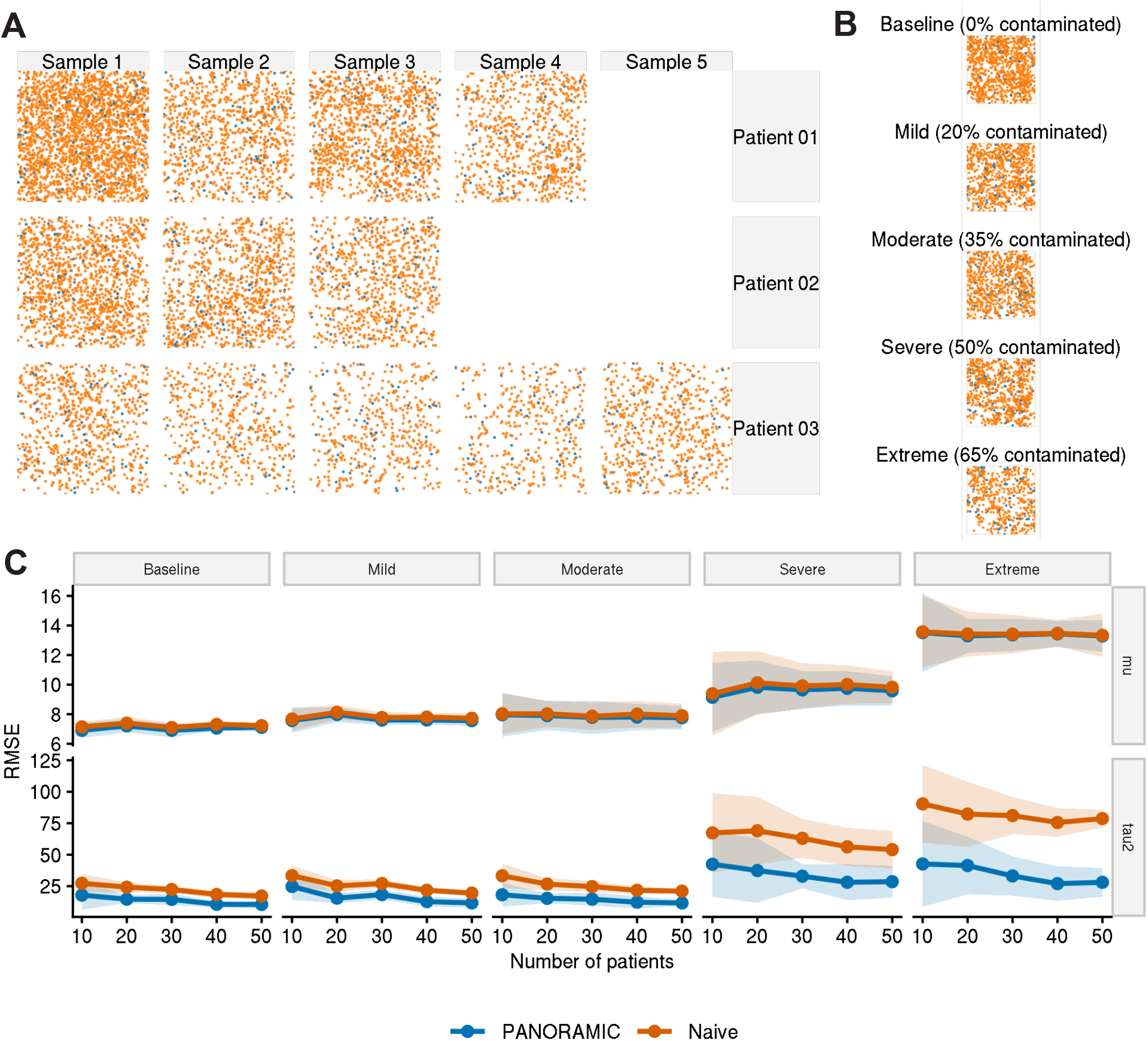
Validation of between-sample variance estimation using PANORAMIC. **(A)** Hierarchical simulation framework produces multiple samples per patient, selected via a negative binomial distribution. Three example patients shown here. **(B)** Progressive degradation of patient samples is performed to study the benefits of PANORAMIC under varying degrees of noise. **(C)** RMSE for the pooled mean *µ* and patient-level heterogeneity *τ*^2^ across patient-count settings *n*_patients_ = 10, …, 50. Confidence intervals are formed from 5 repeated simulations.

Across patient-count and degradation regimes, PANORAMIC and naive estimation produced broadly similar recovery of the pooled mean *µ*, particularly as the number of patients increased. The clearer and more persistent advantage of PANORAMIC was in recovery of patient-level heterogeneity. In the breakpoint stress test, the naive estimator increasingly conflated within-sample noise and degraded sample variation with true between-patient heterogeneity, whereas PANORAMIC maintained substantially better recovery of 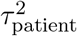, with scenario-specific root mean sqquared error (RMSE) reductions that were largest under severe and extreme degradation (**Figure 3C**).

These simulation results support the central statistical motivation for PANORAMIC: sample-specific uncertainty should be propagated into the level of the model where group and patient heterogeneity are estimated. This distinction is most important in heterogeneous, degraded, or small-to-moderate cohort settings, which are common in spatial omics studies.

### PANORAMIC identifies enriched B- and T-cell colocalizations in CRC tumors with Crohn’s-like reaction

We applied PANORAMIC to the colorectal cancer TMA cohort from Schürch et al., comparing tumors with Crohn’s-like reaction (CLR) against tumors with diffuse inflammatory infiltration (DII). The primary analysis used one randomly selected core per patient to avoid treating multiple TMA cores from the same patient as independent biological replicates. At *r* = 25 *µ*m, PANORAMIC tested 144 directional cell-type pairs and identified 13 differential cell-type pair colocalizations after Benjamini–Hochberg correction (**Figure 4**; **Supplementary Table S8.3**).

**Figure 4.**
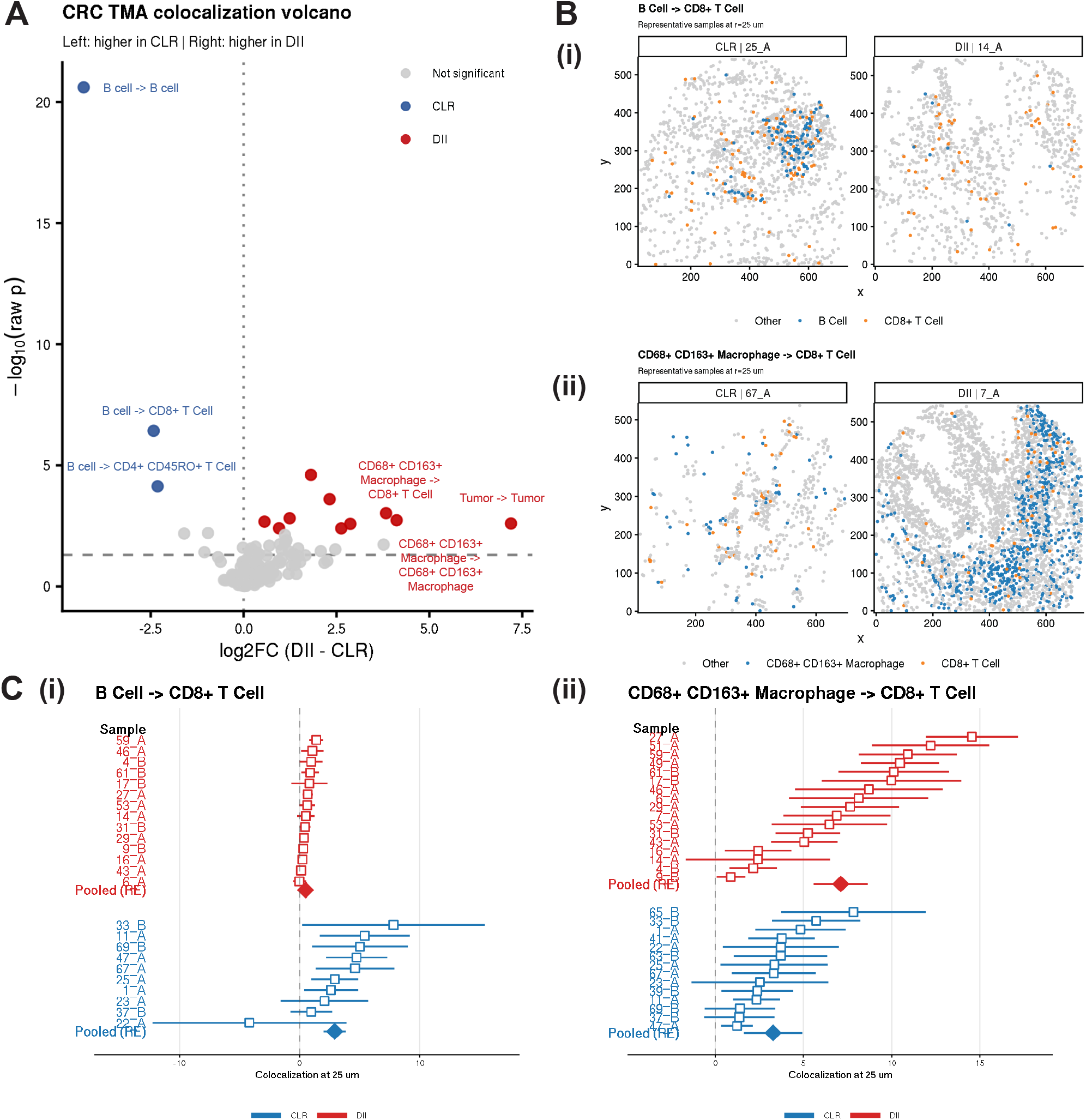
PANORAMIC identifies significantly differentially colocalized cell type pairs in CLR vs DII colorectal cancer. **(A)** Volcano plot showing differential colocalization between colorectal cancer phenotypes CLR and DII at 25 *µ*m. Each point represents a cell-type pair, with the effect size on the x-axis defined as the difference in pooled colocalization means (DII-CLR) and y-axis as the p-values from PANORAMIC’s Wald test. Points are colored by significance. 13 cell type pairs show significant colocalization differences across groups (3 in CLR, 10 in DII) after multiple testing correction. **(B)** Representative CLR and DII tissue cores for (i) B cell → CD8+ T cell colocalization and (ii) CD68+ CD163+ macrophage → CD8+ T cell colocalization. The selected cores were chosen from samples closest to their respective pooled group mean from the meta-analysis. Colored points highlight the relevant cell type populations; gray points denote other cell types. **(C)** Forest plots display the sample level colocalization estimates (white squares) and the uncertainty estimated through the bootstrap (red and blue horizontal bars). The pooled estimate from meta-analysis for each group is indicated by the red and blue diamonds.

The strongest CLR-enriched signals involved B-cell and T-cell organization, including B cell → B cell, B cell → CD8+ T cell, and B cell → CD4+ CD45RO+ T cell colocalization. These findings are consistent with tighter B- and T-cell organization in CLR tumors and align with the expected tertiary-lymphoid-like immune architecture described in this phenotype. Selected macrophage- and tumor-associated colocalizations were stronger in DII; for example, CD68+ CD163+ macrophage → CD8+ T cell colocalization was higher in DII.

To support interpretation, we visualized representative TMA cores closest to the group-level colocalization means for selected cell-type pair colocalization, rather than visually extreme examples (**Figure 4B**). We also generated forest plots for selected features to show sample-level estimates, bootstrap-derived uncertainty, and pooled group effects (**Figure 4C**). These plots illustrate a practical benefit of PANORAMIC: the same analysis reports both the estimated colocalization effect and its uncertainty across samples and patients.

### Differential colocalization network reveals coordinated CLR organization

Pairwise colocalization analyses identify individual cell-type relationships, but do not directly show whether multiple cell types reorganize as a coordinated spatial module. To summarize higher-order structure, we constructed a weighted differential colocalization network from the CRC contrast statistics and focused on the CLR-enriched component. The resulting undirected network contained 12 nodes, 21 edges, and 3 Leiden clusters (**Figure 5A**).

**Figure 5.**
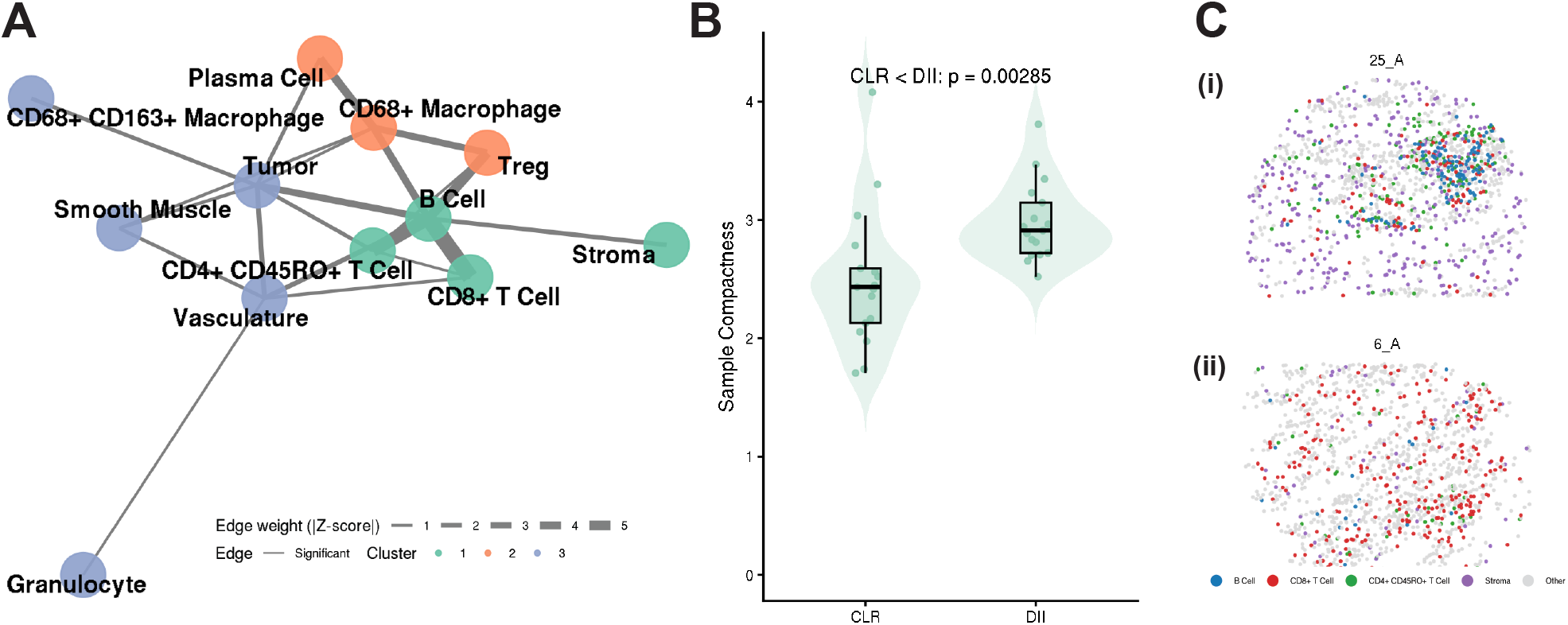
Colocalization network reveals coordinated immune subnetworks in CLR. **(A)** Differential colocalization network constructed from PANORAMIC *Z*-scores for CLR-enriched associations. Edges are weighted by the magnitude of Wald *Z*-scores from differential testing between CLR and DII groups, and Leiden clustering identifies three subnetworks. **(B)** For cluster 1 (green), spatial compactness scores of the relevant cell types were computed for each sample. CLR samples have significantly lower relative compactness scores than DII samples, indicating tighter spatial organization of the selected cell-type set (*p* ≈ 0.003, one-sided *t*-test). **(C)** Representative CLR and DII cores illustrate compact immune clustering in a sample from a CLR patient (i) versus a sample from a patient with DII (ii). The chosen samples were those closest to the median compactness score per group.

One cluster highlighted a CLR-associated immune/stromal module involving B cells, CD8+ T cells, CD4+ CD45RO+ T cells, and stroma. To test whether this network signal corresponded to tighter spatial organization at the sample level, we performed a compactness analysis on this cell-type set. CLR samples had lower relative compactness values than DII samples (**Figure 5B**). The one-sided Welch test for the alternative CLR *<* DII yielded *p* = 0.00285.

Representative CLR and DII samples near their respective group means showed the same qualitative pattern: the selected immune/stromal cell types appeared more tightly organized in CLR and more dispersed in DII (**Figure 5C**). Together, the network and spatial compactness analyses suggest that the CLR-associated colocalization signal is not limited to isolated pairwise associations, but is consistent with broader coordinated immune organization.

### Sensitivity analyses support scale stability and contextualize benchmark performance

We evaluated the scale sensitivity of the CRC results by rerunning PANORAMIC across radii from 25 to 150 *µ*m. Several leading differential features persisted across multiple radii, whereas others were significant only at a subset of spatial scales (**Supplementary Figure S8.5**). These results indicate that the CLR–DII contrast is not driven solely by a single arbitrary neighborhood radius, while also showing that the strength and composition of the signal depend on spatial scale.

We also compared PANORAMIC with spicyR and two-sample Welch *t*-tests for every cell-type pair (**Supplementary Tables S8.6, S8.7**). In the CRC benchmark, PANORAMIC recovered a broader set of biologically coherent B- and T-cell-associated CLR signals than the benchmark workflows.

As an additional diagnostic, we examined empirical *p*-value distributions across tested cell-type pairs (**Supplementary Figure S8.8**). The PANORAMIC histogram showed an approximately uniform background with enrichment of small *p*-values consistent with the pattern from a small number of non-zero differences among a large number of nulls, whereas benchmark workflows showed less interpretable distributions.

### HNSCC analysis demonstrates portability to a second TMA cohort

We then applied PANORAMIC to a head and neck squamous cell carcinoma tissue microarray cohort from Haist et al. [8]. This analysis compared benign lymph node samples from node-negative patients with tumor-adjacent lymph node regions from patients with lymph node metastasis. Unlike the primary CRC analysis, this setting included repeated tissue regions per patient and therefore evaluated PANORAMIC in a nested sampling design.

At *r* = 25 *µ*m, PANORAMIC tested 324 directional cell-type pairs and identified one cell-type pair colocalization at the 5% FDR-significance level, NK cells → *γδ* T cells (**Supplementary Table S9.2**). This analysis shows that the workflow transfers to a second mIF TMA cohort with a different tissue context, comparison structure, and repeated-sampling design.

## Discussion

Spatial omics studies increasingly compare tissue organization across patients, tissue regions, and clinical conditions, but many analysis workflows still treat sample-level spatial summaries as error-free quantities. This can obscure the distinction between uncertainty within a tissue sample and biological heterogeneity across patients or samples. PANORAMIC addresses this gap by combining an interpretable local colocalization statistic, spatial bootstrap estimation of within-sample uncertainty, and multilevel random-effects meta-analysis. The resulting framework allows each tissue sample to contribute both an estimated spatial effect and an uncertainty estimate, while the cohort-level model accounts for heterogeneity across patients and repeated tissue regions.

Three features of PANORAMIC are most important for interpretation. First, the default statistic used here, edge-corrected neighborhood enrichment, measures local deviation from expected cell-type composition around anchor cells. It should be interpreted as a local spatial enrichment statistic, not as proof of direct cell–cell interaction. Second, the spatial bootstrap estimates sample-specific uncertainty by resampling local tissue regions rather than individual cells. This preserves local spatial dependence and accounts for boundary truncation conditional on the supplied tissue region. Third, the multilevel meta-analysis propagates these sample-specific uncertainty estimates into group-level fixed effect estimates, differential tests, and heterogeneity estimates. Together, these components make PANORAMIC most useful when the scientific question concerns reproducible spatial organization across samples rather than cell-level associations within a single image.

The simulation studies support two complementary conclusions. The first is a calibration conclusion: bootstrap-derived within-sample uncertainty estimates were strongly aligned with empirical variance in uniform and clustered local spatial regimes, but less strongly aligned when broad opposing gradients dominated the tissue architecture. This indicates that local tile resampling is informative for many neighborhood-scale spatial patterns, while also highlighting the need for tissue masks, region definitions, or stratified analyses when large-scale tissue compartments drive the signal. The second is a heterogeneity-estimation conclusion: PANORAMIC gave its clearest advantage over naive estimators for estimating patient-level heterogeneity, especially under contamination and sample-degradation scenarios. This distinction is important because the main value of PANORAMIC is not only generating an efficient point estimate of colocalization, but separating within-sample spatial uncertainty from between-patient variability.

In the CRC TMA application, PANORAMIC identified differential colocalization patterns that distinguish CLR from DII tumors and are consistent with the known immune architecture of the dataset. B-cell enriched and B/T-cell colocalization patterns were stronger in CLR, consistent with organized lymphoid aggregate biology, whereas selected macrophage- and tumor-associated patterns were stronger in DII. The representative tissue cores and forest plots illustrate a practical benefit of PANORAMIC: the analysis reports group-level effects while retaining information about sample-level variability and uncertainty. The differential network and compactness analyses further suggest that the CLR-associated signal is not limited to isolated pairwise associations, but is consistent with broader coordinated immune/stromal organization.

The supplementary benchmark analyses provide context for interpreting these findings. Comparisons with spicyR and Welch *t*-test baselines should not be interpreted as feature-identical comparisons because the methods use different spatial summaries and uncertainty models. However, under matched CRC analysis settings, PANORAMIC recovered a broader set of CLR-enriched B- and T-cell-associated signals and produced empirical distributions of p-values consistent with a mixture of null and strong alternative signals. These diagnostics do not prove formal false discovery rate control, but they support the empirical calibration and biological coherence of the CRC results. The HNSCC TMA analysis further demonstrates that the same workflow can be applied to a second mIF cohort with repeated tissue regions per patient, although we interpret that analysis primarily as a portability and nested-design demonstration rather than as a comprehensive biological study of metastatic lymph node remodeling.

PANORAMIC has several limitations. First, while it propagates uncertainty for a chosen spatial statistic, it does not remove the need to select a biologically appropriate definition of colocalization. Different statistics capture different spatial features, and results should be interpreted relative to the statistic and radius used. Second, radius choice remains biologically important and affects the construction of tiles resampled in the spatial bootstrap. Multi-radius analyses can assess scale stability, but they cannot replace prior knowledge about the spatial scale of the biological process. Third, PANORAMIC depends on accurate cell-type annotations and tissue-window definitions. Highly irregular, perforated, necrotic, or compartmentalized tissues may require careful tissue masks or stratified analyses using pathologist-defined regions. Fourth, computational cost increases with the number of cell types, tested radii, bootstrap replicates, samples, and random-effects complexity. Finally, as with any meta-analytic approach, small effective sample sizes limit the precision of between-patient heterogeneity estimates even when within-sample uncertainty is modeled correctly.

Despite these limitations, PANORAMIC provides a practical framework for cohort-level spatial colocalization analysis. By carrying forward both sample-level effects and uncertainty estimates, PANORAMIC supports pooled estimates, heterogeneity summaries, forest plots, differential tests, and differential colocalization networks that are more transparent than sample-level point estimates alone. Future work can extend this uncertainty-propagation principle to additional spatial summaries, covariate-adjusted meta-regression, and broader spatial omics settings, including spatial transcriptomic and multimodal workflows where cell types, cell states, or spatial domains can be represented as marked point patterns.

## Data Availability

The colorectal cancer tissue microarray data analyzed in this study were previously published by Schürch et al. [21]. The released single-cell CODEX data and associated metadata are available from the original data repository (https://data.mendeley.com/datasets/mpjzbtfgfr/1). The head and neck squamous cell carcinoma tissue microarray data analyzed in the supplementary application were previously published by Haist et al. [8] and are available through the data resources associated with that publication (https://data.mendeley.com/datasets/t4k5kj3cxr/1). No new experimental data were generated for this study.

## Code Availability

The PANORAMIC framework is released as an open-source R package at https://bioconductor.org/packages/devel/bioc/html/panoramic.html and archived on Zenodo with DOI: 10.5281/zenodo.19927197. All scripts used to preprocess data, run simulations, conduct the CRC and HNSCC analyses, perform benchmarking, and generate manuscript figures are available at https://github.com/chang-jacob/panoramic-paper. The analysis repository includes the code needed to reproduce the results from the publicly available input datasets.

## Supplementary Methods

### S1 Edge-corrected neighborhood enrichment colocalization statistic

The default colocatization statistic for PANORAMIC is the edge-corrected neighborhood enrichment statistic, as described in the main text, using the generic tissue window *W*. Here, we define *W*_*k*_ as the observed tissue window of sample *k*, with area |*W*_*k*_|, and describe the implementation details used for self-pairs and boundary correction.

For an ordered pair *a* → *b*, PANORAMIC uses the same direction convention as in the main text: *a* → *b* denotes enrichment of type-*a* cells around type-*b* anchor cells. For self-pairs (i.e. *a* = *b*), the anchor cell itself is excluded from both the observed neighborhood count and the expected target-cell count. We denote the corresponding leave-one-anchor density by 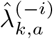, where the superscript (−*i*) indicates that anchor cell *i* is omitted when *a* = *b*:

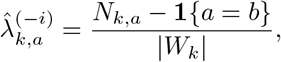

where *N*_*k,a*_ is the number of cells of type *a* in sample *k*. For non-self pairs (i.e. *a* ≠ *b*), the anchor cell is not a type-*a* target cell, so this reduces to the usual sample-wide density *N*_*k,a*_*/*|*W*_*k*_|. For self-pairs (i.e. *a* = *b*), this reduces to (*N*_*k,a*_ − 1)*/* |*W*_*k*_|, preventing the anchor cell from contributing to its own expected target-cell density.

For anchor cell *i*, the expected count is

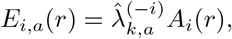

where

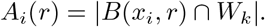

Thus, the expected number of target cells is computed using the available in-tissue area *A*_*i*_(*r*), rather than the full disc area *πr*^2^. This gives anchors near tissue boundaries smaller expected target-cell counts.

Equivalently, the anchor-level statistic from the main text can be written as

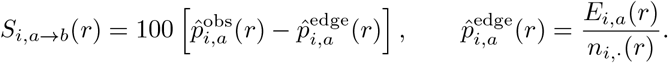

Here, 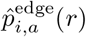 is an edge-corrected expected target-cell count divided by the *n*_*i*._(*r*).

The edge-corrected neighborhood enrichment statistic is conceptually related to graph-based neighborhood enrichment and co-occurrence analyses used in spatial omics software, including Squidpy and Giotto [1, 2]. PANORAMIC differs in three main ways: neighborhoods are defined by metric balls at fixed physical radii; expected target-cell counts are adjusted for available tissue area around each anchor; and sample-level estimates are paired with spatial-bootstrap variance estimates for multilevel random-effects meta-analysis. The boundary correction also draws on the broader spatial point-pattern literature, where edge correction is needed because neighborhoods near the observation-window boundary are only partially observed [3, 4].

### S2 Spatial bootstrap implementation

For each feature *f*, sample *k*, and radius *r*, let *S*_*i,f,k*_(*r*) denote the anchor-level neighborhood enrichment score for valid anchor *i*. In the analyses in this manuscript, a feature *f* corresponds to an ordered cell-type pair *a* → *b* evaluated at a specified radius.

PANORAMIC overlays the sample window *W*_*k*_ with a regular grid of square tiles. Let 𝒢_*k*_ denote the set of tiles with positive overlap with the observed tissue window. Each retained tile receives the overlap weight

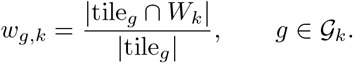

Tiles with zero overlap with *W*_*k*_ are excluded from the bootstrap sampling frame.

Each valid anchor is assigned to one retained tile according to its spatial coordinate. Let 𝒜_*g,f,k*_ denote the set of valid anchors for feature *f* assigned to tile *g*. For each retained tile, PANORAMIC computes a tile-level summary

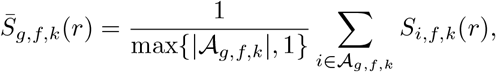

with the convention that tiles containing no valid anchors for feature *f* contribute a zero tile-level sum. This convention keeps the bootstrap sampling frame tied to the observed tissue window rather than redefining the set of sampled tiles separately for each feature.

Bootstrap resampling probabilities are proportional to the overlap weights,

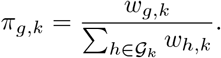

For bootstrap replicate *b*, PANORAMIC samples tiles

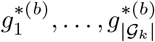

with replacement from 𝒢_*k*_ using probabilities *π*_*g,k*_. The replicate statistic is

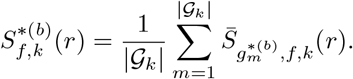

Thus, overlap weights affect how often each tile is sampled.

The bootstrap-centered sample-level estimate is the mean of the replicate statistics,

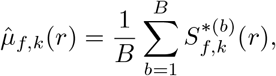

and the within-sample variance estimate is

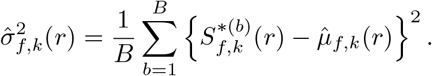

These define the sample-level effect estimate and sampling variance used in the downstream meta-analysis:

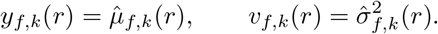

This procedure is a PANORAMIC-specific adaptation of Loh-style spatial resampling [5, 6]. Rather than resampling individual cells, which would ignore local spatial dependence, the method resamples local spatial regions represented by tiles.

### S3 Multilevel meta-analysis implementation

PANORAMIC fits each feature independently using `metafor::rma.mv`. The input for feature *f* consists of sample-level estimates *y*_*fk*_ and corresponding sampling variances *v*_*fk*_ obtained from the spatial bootstrap. The estimated variances *v*_*fk*_ are passed to the meta-analytic model as known sampling variances, while random-effect terms model residual heterogeneity not explained by the within-sample bootstrap variance.

When a grouping variable is supplied, PANORAMIC estimates group-specific pooled means using fixed-effect moderators with no intercept. In matrix notation, the patient-level model can be written as

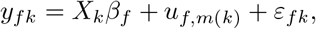

where *X*_*k*_ encodes group membership or other sample-level moderators for sample *k, m*(*k*) denotes the patient from which sample *k* was obtained,

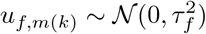

is a patient-level random effect, and

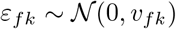

is the known sampling-error term from the spatial bootstrap.

For analyses with repeated regions or cores from the same patient, PANORAMIC can fit a nested patient-plus-sample random-effects structure,

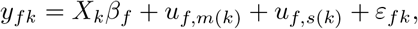

where *s*(*k*) denotes the sample or tissue region corresponding to observation *k* and

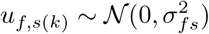

is the sample-level random effect within a patient. In the package implementation, these two structures correspond to `tau_structure = “patient”` and `tau_structure = “patient_sample”`, respectively. The nested model is implemented as a random intercept for samples nested within patients.

For two-group comparisons, PANORAMIC reports the contrast between the two estimated group means. In the package output, `beta_diff` is computed as the second group minus the first group, where group order is determined by the factor levels of the grouping variable. The standard error of the contrast is computed from the fitted coefficient covariance matrix, and the Wald statistic is

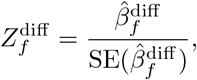

where 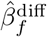 is a point estimator of 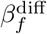 and 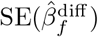 is the estimated standard error of 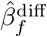. Two-sided *p*-values are computed using the standard normal reference distribution, and Benjamini–Hochberg correction is applied across tested features.

PANORAMIC also supports variance flooring for non-positive or numerically unstable within-sample variance estimates. With `vi_floor = “median”`, non-positive *v*_*fk*_ values are replaced by the median positive variance for that feature. With `vi_floor = “group_median”`, the replacement is computed within each group. With `vi_floor = “none”`, sample-feature combinations with non-positive variances are dropped before model fitting. Variance flooring prevents samples with zero or near-zero bootstrap variance from receiving infinite or numerically dominant inverse-variance weight in the meta-analysis.

### S4 Simulation design

#### S4.1 Intrasample variance calibration simulation

To evaluate whether PANORAMIC’s spatial bootstrap provides informative within-sample variance estimates, we simulated marked spatial point patterns under three regimes: uniform random mixing, opposing gradients, and clustered organization. Each simulated sample occupied a 600 *µ*m × 600 *µ*m square window and contained five cell types: A, B, C, Rare 1, and Rare 2. Baseline cell-type intensities were scaled so that the expected total number of simulated cells matched the average number of cells in the CRC TMA cores.

The “uniform regime” generated each cell type from an independent homogeneous Poisson process. The “opposing-gradient” regime generated types A and B from opposing inhomogeneous Poisson processes, producing large-scale compositional heterogeneity without direct local attraction. The “clustered regime” generated *B* → *A* spatial structure using a parent-offspring process, with A cells corresponding to parent points and B cells generated around parent locations; the remaining cell types were generated from independent homogeneous Poisson processes.

For each regime, we generated 300 simulated samples. PANORAMIC was applied to each sample using edge-corrected neighborhood enrichment at radius *r* = 25 *µ*m, block bootstrap uncertainty estimation with *B* = 100 replicates, and square tiles of side length 62.5 *µ*m. For each feature, empirical variance was estimated across independent simulation replicates and compared with the mean PANORAMIC bootstrap variance. We summarized variance calibration using Pearson correlations.

#### S4.2 Intersample degradation simulation

To evaluate PANORAMIC under degraded multi-sample sampling, we simulated nested patient-level datasets in which each observed sample was an *B* → *A* clustered point pattern in a 600 *µ*m × 600 *µ*m square window with target total cell count approximately 1900. Patient-level latent effects modified the parent-process intensity, offspring abundance, and cluster spread, inducing between-patient heterogeneity in colocalization. Additional sample-level log-normal perturbations induced within-patient variability across repeated samples. Clean simulated samples were also subjected to global thinning to mimic incomplete or variable tissue sampling, while enforcing minimum total-cell and per-type cell-count thresholds.

We evaluated *n*_patients_ ∈ {10, 20, 30, 40, 50}, with 5 simulation replicates per patient-count setting. The number of samples per patient was drawn from a negative-binomial distribution with mean 2.2 and constrained to lie between 1 and 4. For each patient, 100 independent clean truth realizations were generated without bootstrap resampling to approximate the patient-specific expected colocalization effect and within-patient variance structure. Observed samples were then analyzed with PANORAMIC using edge-corrected neighborhood enrichment at *r* = 25 *µ*m, block bootstrap uncertainty estimation with *B* = 100 replicates, and baseline tile size 62.5 *µ*m. Sample-level effects and variances were passed to REML-based multilevel meta-analysis with patient- and sample-level random effects, and estimates were compared with the corresponding simulation truth.

To assess robustness to controlled departures from clean sampling, we introduced a degradation ladder ranging from a clean baseline to mild, moderate, severe, and extreme degradation. In each scenario, contamination was assigned at the patient level and inherited by that patient’s observed samples. Degradation combined quality-dependent perturbations in four components: increased sample-level heterogeneity, asymmetric thinning by cell type, directional compositional bias favoring one cell type, and inflation of the effective bootstrap tile scale. Minimum total-cell and per-type cell-count thresholds were enforced after degradation.

For each scenario and patient-count regime, we summarized PANORAMIC performance using the root mean squared error (RMSE) of the estimated pooled mean effect and patient-level heterogeneity relative to the simulated truth. Confidence intervals for RMSE were computed across the 5 independent simulation replicates. Simulation outputs also included feature-level estimates, replicate-level performance summaries, and breakpoint-style diagnostics describing how estimation accuracy changed with increasing degradation severity.

### S5 CRC TMA preprocessing and analysis settings

#### S5.1 CRC data source and preprocessing

The CRC TMA dataset was originally generated by Schürch et al. using CODEX multiplexed immunofluorescence imaging of formalin-fixed paraffin-embedded colorectal cancer tissue microarrays sampled from the invasive tumor front [7]. The released single-cell table includes spatial coordinates, cell-type annotations, cellular neighborhood assignments, marker measurements, and patient-level metadata. The original study profiled 140 tissue regions from 35 advanced-stage CRC patients using 56 protein markers and identified nine conserved cellular neighborhoods in the immune tumor microenvironment.

Patients were annotated according to immune infiltration patterns at the invasive tumor front. Patients with lymphoid aggregate architecture consistent with Crohn’s-like reaction were annotated as CLR, whereas patients with diffuse inflammatory infiltration were annotated as DII. For CLR patients, cores included samples with lymphoid aggregate samples with diffuse regions; for DII patients, only cores with diffuse infiltration regions are sampled.

Starting from the released single-cell table, we retained cells with defined cell-type annotations and spatial coordinates. Spatial coordinates were converted from pixels to microns using the dataset-specific conversion factor 0.37742 *µ*m per pixel. We removed undefined cells and excluded cell types that were absent from more than 20% of images, mirroring the preprocessing strategy used for this dataset in the spicyR study [8]. After filtering, the processed table contained 228,811 cells across 35 patients, 140 spots, and 12 retained cell types.

#### S5.2 One-core-per-patient subsampling

For the primary CRC analysis, we restricted the dataset to one core per patient to preserve patient-level biological replication. We first retained cores whose annotations matched the group-specific comparison of interest: lymphoid aggregate cores for CLR patients and diffuse infiltration cores for DII patients. From this filtered set, one core was randomly selected per patient using a fixed random seed.

#### S5.3 PANORAMIC CRC analysis settings

The primary PANORAMIC run used the edge-corrected neighborhood enrichment statistic at *r* = 25 *µ*m, *B* = 100 spatial bootstrap replicates, and square bootstrap tiles with side length 62.5 *µ*m, corresponding to 2.5 × *r*. The analysis used the block bootstrap and a convex tissue window. Features were required to have at least five relevant cells per sample; otherwise, that sample-feature combination was marked as missing. Feature-wise group comparisons between CLR and DII were fit using multilevel random-effects meta-analysis with patient-level random effects, group-specific fixed-effect means, group-median variance flooring, and REML estimation. Multiple testing correction was performed using the Benjamini–Hochberg procedure across tested features.

#### S5.4 CRC differential colocalization network analysis

To summarize coordinated spatial changes across multiple cell types, we constructed a differential colocalization network from the PANORAMIC CRC differential testing results. The network was based on the primary one-core-per-patient CRC analysis using edge-corrected neighborhood enrichment at *r* = 25 *µ*m, *B* = 100 bootstrap replicates, and multilevel meta-analysis.

Each node in the network represents a retained cell type. Candidate edges correspond to ordered cell-type pairs tested by PANORAMIC. For the main CRC network visualization, we focused on CLR-enriched associations. Because the differential effect in the primary output was defined as DII minus CLR, CLR-enriched associations corresponded to negative differential effects and negative Wald *Z*-statistics. We therefore constructed the displayed network from the CLR-up component using significant features with Benjamini– Hochberg adjusted *p <* 0.05 and *Z <* 0.

Although PANORAMIC tests ordered cell-type pairs, the network was rendered as an undirected graph for visualization of higher-order cell-type organization. Edge weights were derived from the magnitude of the Wald *Z*-statistics from the differential colocalization tests. Leiden clustering was then applied to the resulting graph to identify groups of cell types participating in coordinated differential colocalization patterns [9]. In the plotted network, we used a Leiden resolution parameter of 1.2. Non-significant edges were optionally retained at low visual weight to preserve network context, but interpretation focused on the FDR-significant CLR-enriched edges.

The CLR-enriched network highlighted an immune/stromal module involving B cells, CD8+ T cells, CD4+ CD45RO+ T cells, and stroma. We used this cell-type set for the compactness analysis described below, which provides a sample-level spatial summary of whether the network-highlighted cell types were more tightly co-organized in CLR than in DII samples.

#### S5.5 CRC spatial compactness analysis

Let 𝒯 denote the set of cell types selected from the differential colocalization network. In the CRC analysis, we used

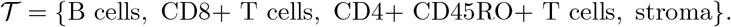

For sample *k*, let ℐ_*k*, 𝒯_ denote the set of cells whose labels belong to 𝒯. For each source cell *i* ∈ ℐ_*k*,𝒯_ and each target type *t* ∈ 𝒯, define

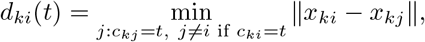

where *x*_*ki*_ is the coordinate of cell *i* ∈ ℐ_*k*,T_ and ∥ *x*_*ki*_ − *x*_*kj*_ ∥ reprensents the distance between cell *i* and cell *j*. When no target cell of type *t* was available, the distance was set to infinity. Distances were capped at *r*_0_ = 25 *µ*m:

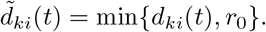

The within-set spacing for source cell *i* was defined as the average capped nearest-neighbor distance to the selected target types,

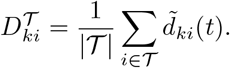

To account for differences in overall cell density, we computed a local baseline distance

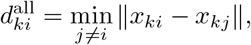

the nearest-neighbor distance from cell *i* to any other cell in the sample. The relative compactness value for cell *i* was

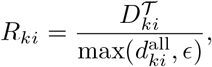

where *ϵ* = 10^−8^ was used for numerical stability. The sample-level relative compactness score was then computed as

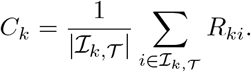

Smaller values of *C*_*k*_ indicate that the selected cell types are closer to one another relative to the local nearest-neighbor spacing of the sample.

For each ordered pair of selected cell types, we also recorded sample-level mean, median, and standard deviation of capped nearest-neighbor distances. Pairwise distances and spatial compactness summaries were compared between CLR and DII samples using Welch *t*-tests. The primary spatial compactness comparison used *C*_*k*_, the mean relative compactness score, and tested the one-sided alternative that CLR samples have lower relative compactness than DII samples.

### S6 HNSCC TMA preprocessing and analysis settings

#### S6.1 HNSCC data source and comparison definition

The HNSCC tissue microarray cohort was obtained from Haist et al., who investigated how lymph node colonization remodels primary tumors and tumor-draining lymph nodes in head and neck squamous cell carcinoma [10]. The dataset includes spatially resolved profiling of primary tumors and lymph node tissues, including tumor-involved, tumor-adjacent, and benign lymph node regions. Haist et al. reported that nodal colonization was associated with interferon-*γ* signaling, enrichment of immunosuppressive myeloid cells and cancer-associated fibroblasts, and spatially organized myeloid–fibroblast niches associated with T cell dysfunction and Treg activation [10].

For PANORAMIC analysis, we focused on a lymph node comparison designed to evaluate spatial remodeling associated with metastatic involvement. We compared benign lymph node samples from node-negative patients with tumor-adjacent (i.e., cancer-free) lymph node regions from patients with lymph node metastasis. This setting provides a nested design in which multiple tissue regions may be observed for the same patient, allowing evaluation of PANORAMIC’s multilevel modeling framework beyond the one-core-per-patient CRC analysis.

#### S6.2 PANORAMIC HNSCC analysis settings

The HNSCC analysis used the same primary local spatial statistic and bootstrap settings as the CRC analysis: edge-corrected neighborhood enrichment at radius *r* = 25 *µ*m, *B* = 100 spatial bootstrap replicates, square bootstrap tiles of side length 62.5 *µ*m, and a minimum of five relevant cells per sample-feature combination. Feature-wise group comparisons were fit using multilevel random-effects meta-analysis with patient-level or patient-plus-sample random effects, selected according to the repeated-sampling structure available for each feature. Multiple testing correction was performed using the Benjamini–Hochberg procedure across tested ordered cell-type pairs.

We included this analysis primarily to evaluate PANORAMIC in a second real multiplexed imaging dataset with multiple regions per patient. Therefore, the HNSCC analysis is presented as a nested-design demonstration and sensitivity application rather than as a comprehensive biological analysis of metastatic lymph node remodeling.

#### S6.3 HNSCC benchmark settings

To compare PANORAMIC with an existing image-level differential localization workflow, we also analyzed the HNSCC comparison using spicyR. The spicyR benchmark used the same HNSCC comparison, retained cell types, and sample inclusion criteria where possible. Because PANORAMIC and spicyR use different colocalization metrics and modeling assumptions, this benchmark should be interpreted as a comparison of differential spatial-analysis workflows rather than as a feature-identical test of the same estimand.

P-value distributions from the HNSCC analyses were compared to evaluate whether the workflows produced broadly different empirical significance patterns in this second dataset. These comparisons are descriptive and are intended to complement the CRC benchmark, where the primary biological signal and label-permutation analyses are reported.

## Supplementary Results

### S7 Simulation supplementary results

#### S7.1 Progressive degradation and breakpoint results

**Figure S1:**
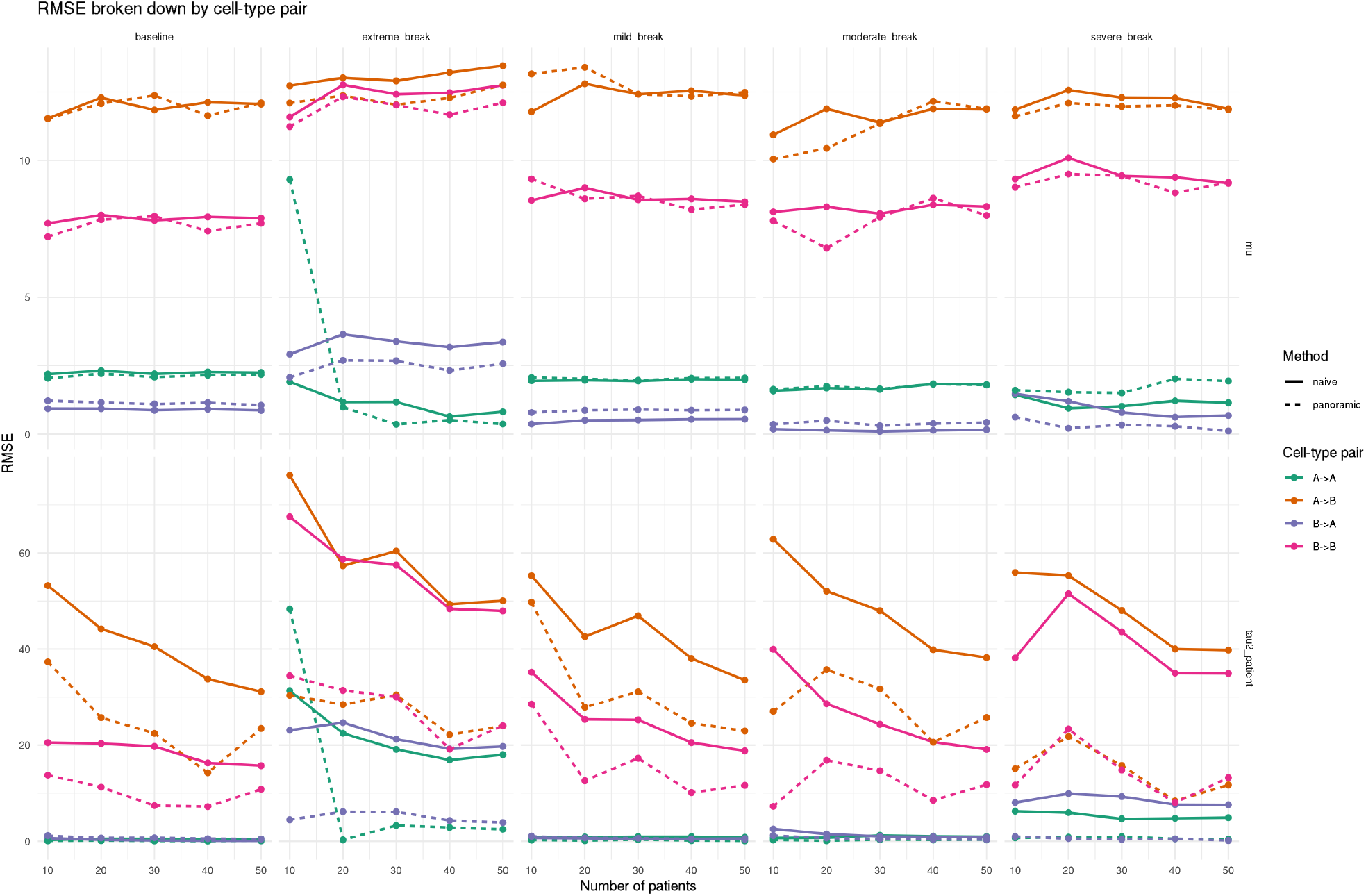
Breakpoint simulation performance by cell-type pair. Root mean squared error (RMSE) is shown for recovery of the pooled mean colocalization effect *µ* and patient-level heterogeneity 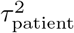 across patient-count regimes and degradation scenarios. Columns correspond to the baseline simulation and progressively degraded settings, from mild through extreme breakpoint contamination. Colors denote ordered cell-type pairs, and line type denotes the estimator: naive estimation versus PANORAMIC. PANORAMIC provides the clearest improvement for estimating patient-level heterogeneity, particularly for A→B and B→B relationships under degraded sampling, whereas improvements for the pooled mean *µ* are smaller and more cell-type-pair dependent. These results support the main simulation finding that PANORAMIC’s primary advantage is separating within-sample uncertainty from between-patient heterogeneity under sample degradation.

### S8 CRC supplementary results

#### S8.1 CRC TMA overview

**Figure S2:**
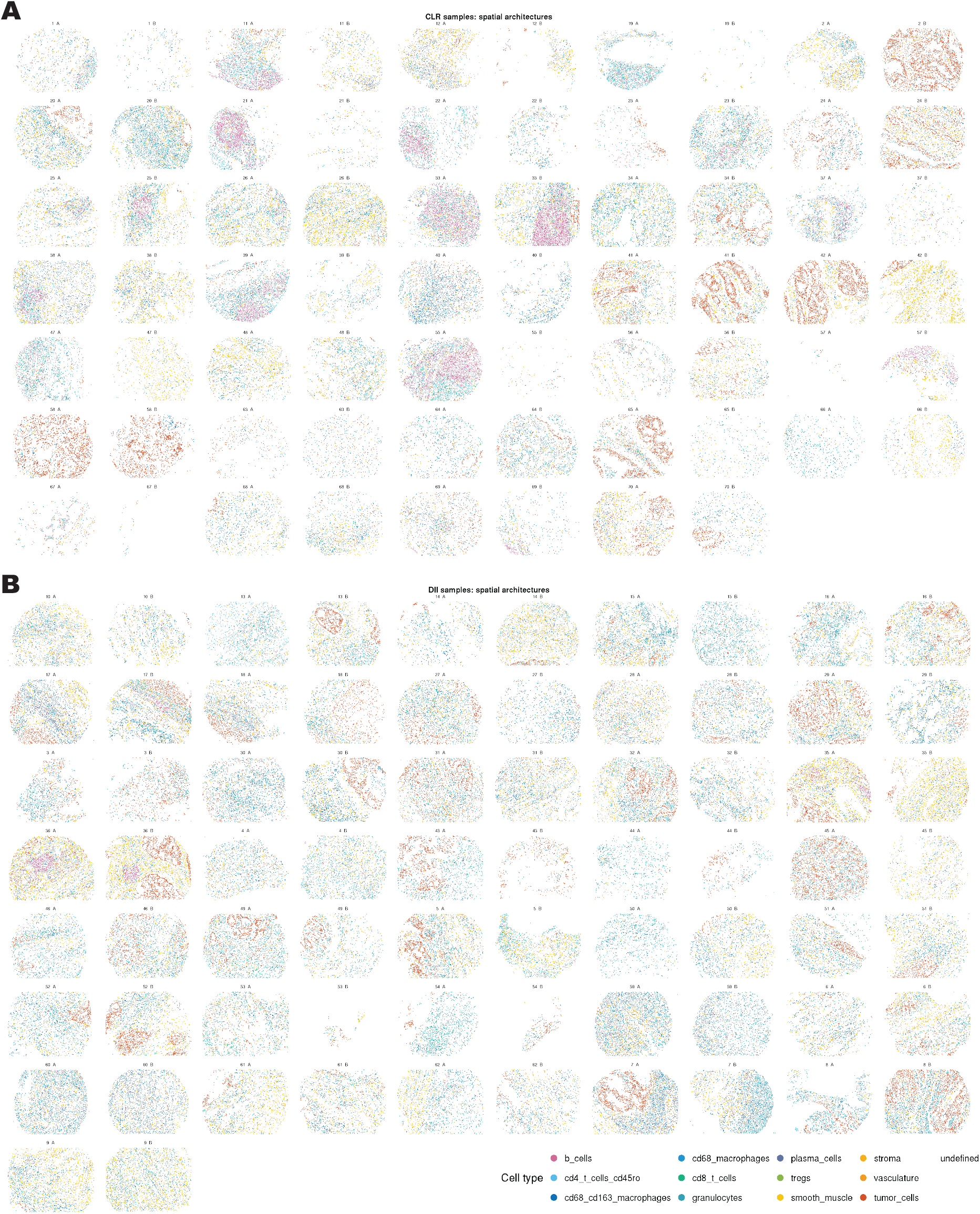
Visual overview of the CRC TMA from Schürch et al.[7]. (A) 68 cores from patients identified to have Crohn’s-like reaction (CLR). (B) 72 cores from patients identified to have diffuse inflammatory infiltration (DII).

#### S8.2 CRC cell-type composition

**Figure S3:**
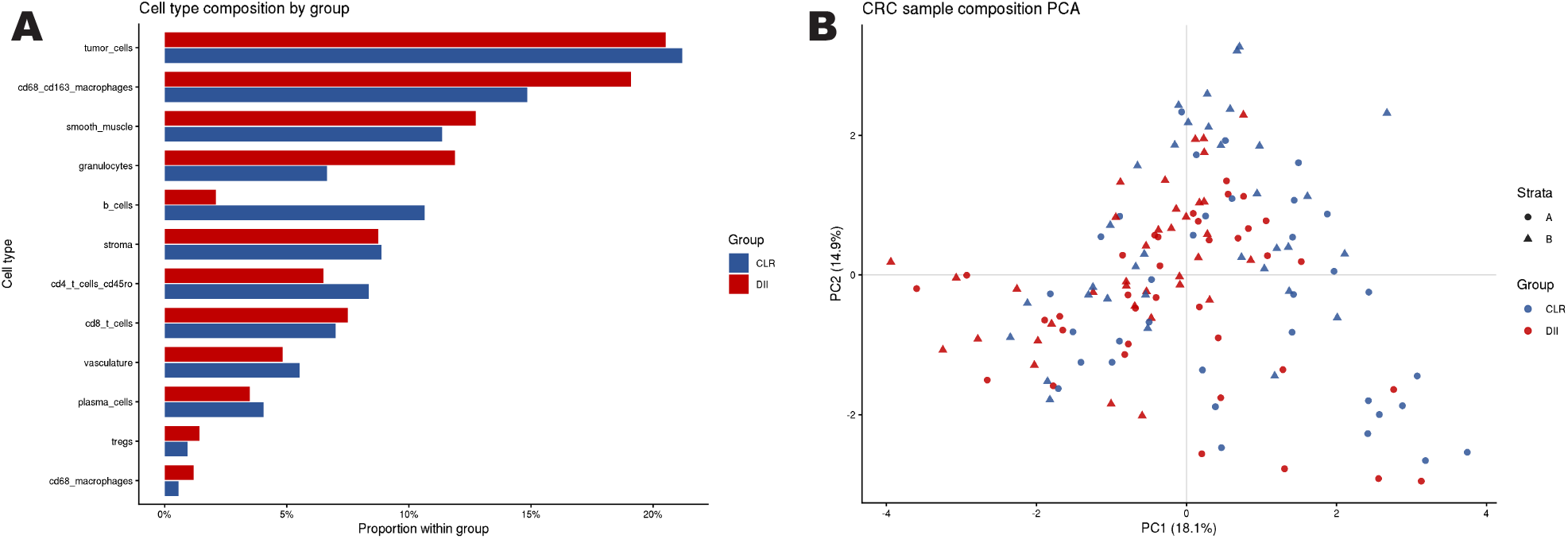
Overview of CRC TMA cell-type proportions. (A) Proportions of cell types between cores from CLR and DII patients. (B) PC1 and PC2 generated from the matrix of cell-type proportions per core.

#### S8.3 Full PANORAMIC CRC results at 25 *µ*m

**Table S1:**
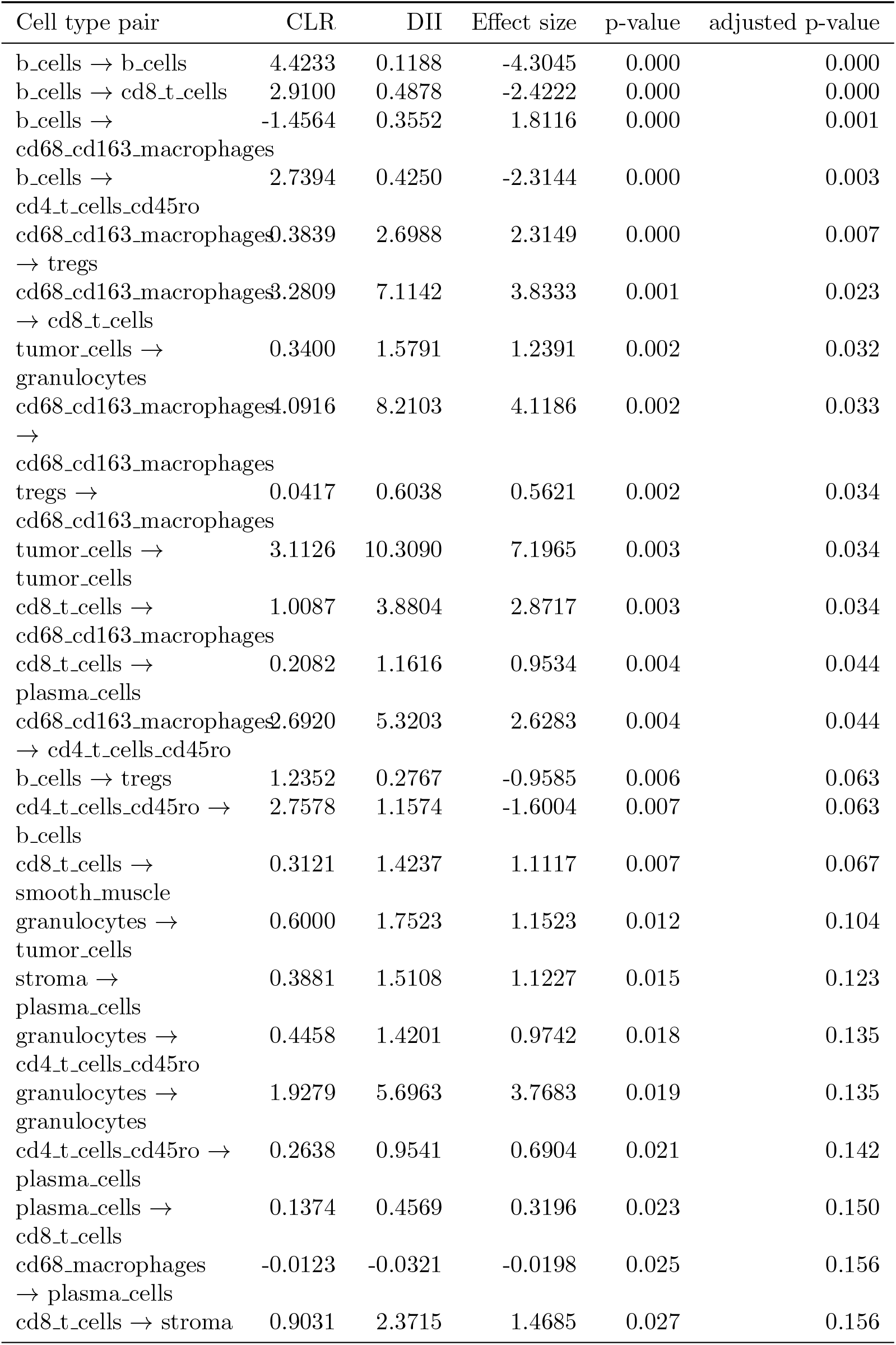
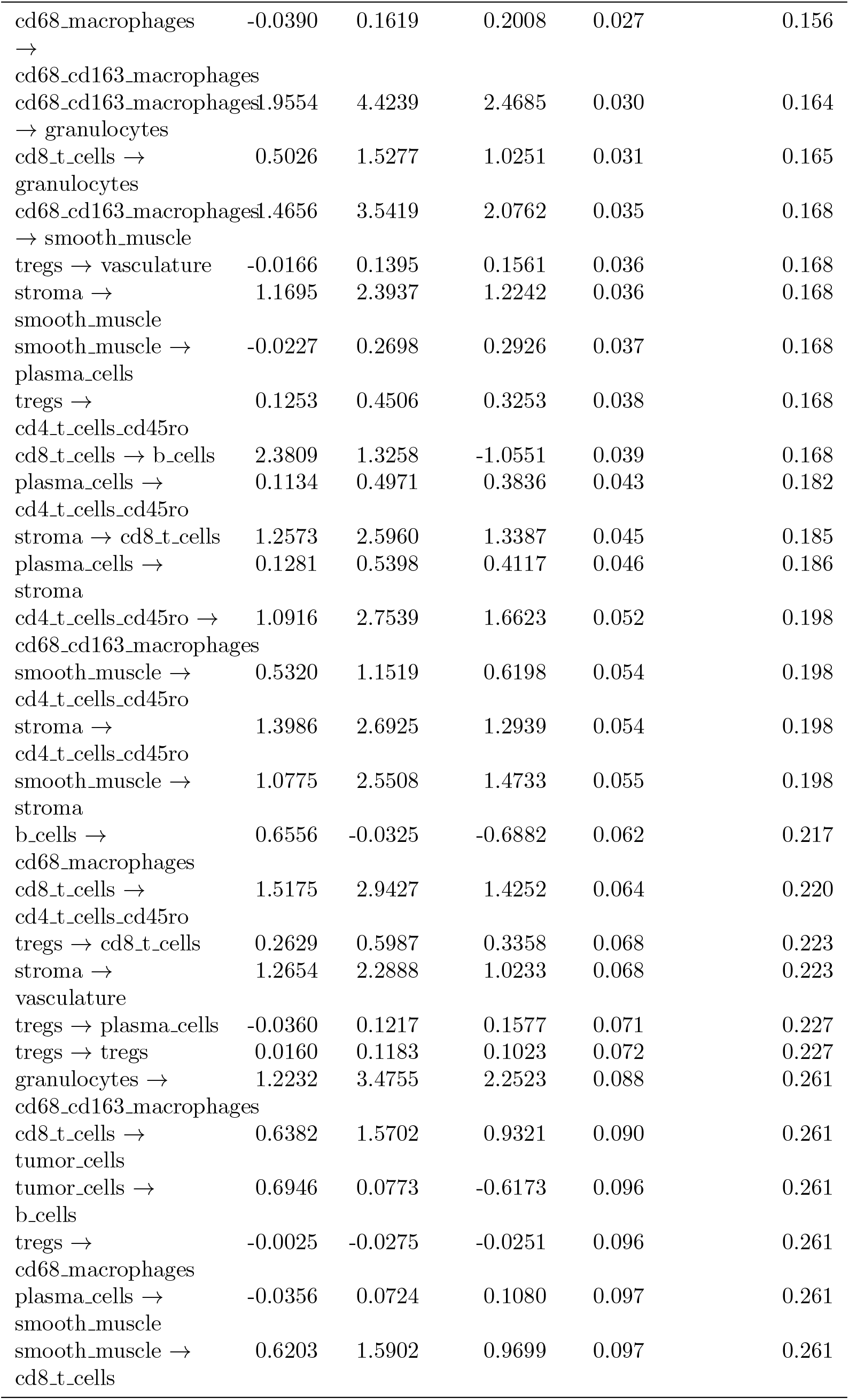
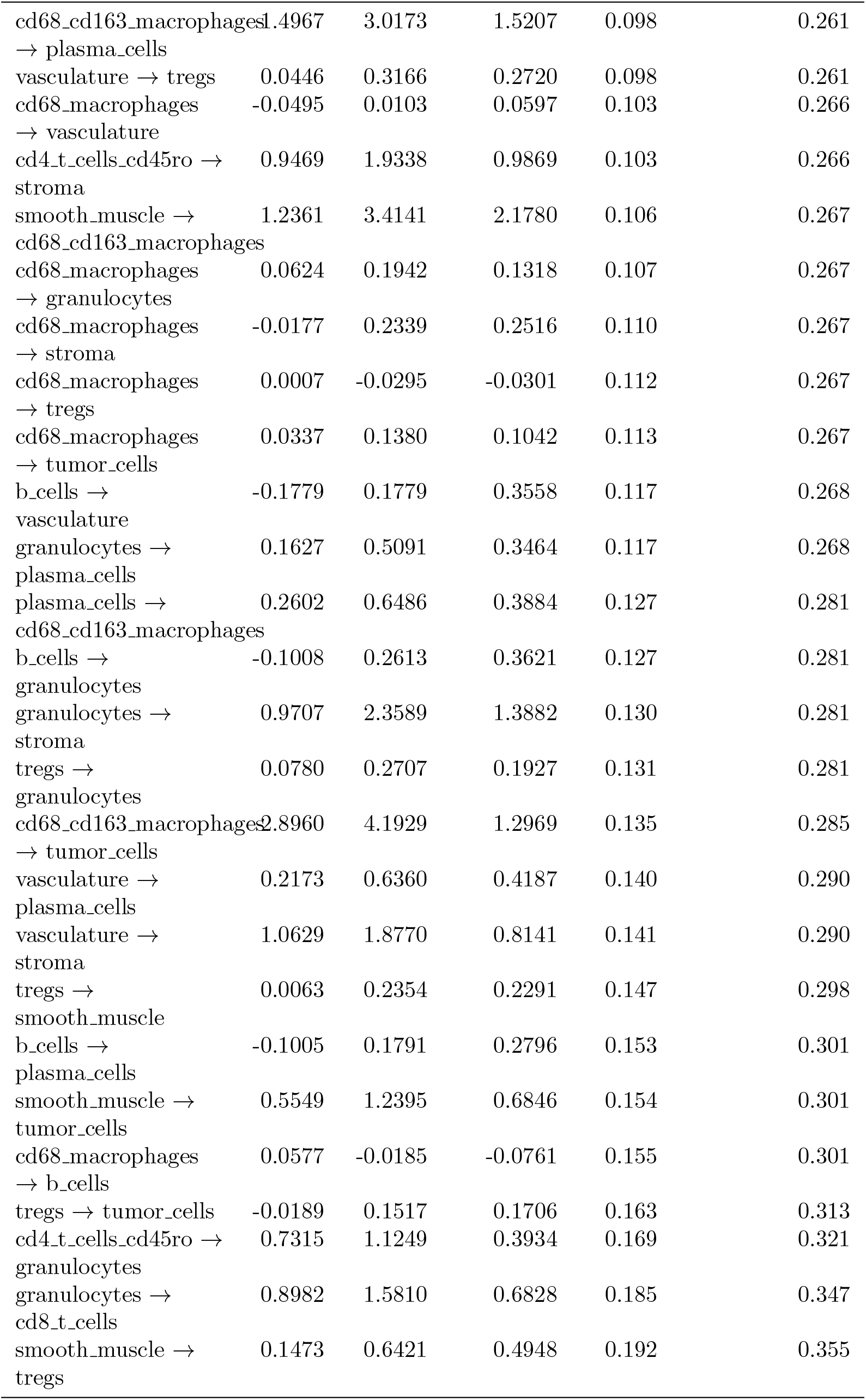
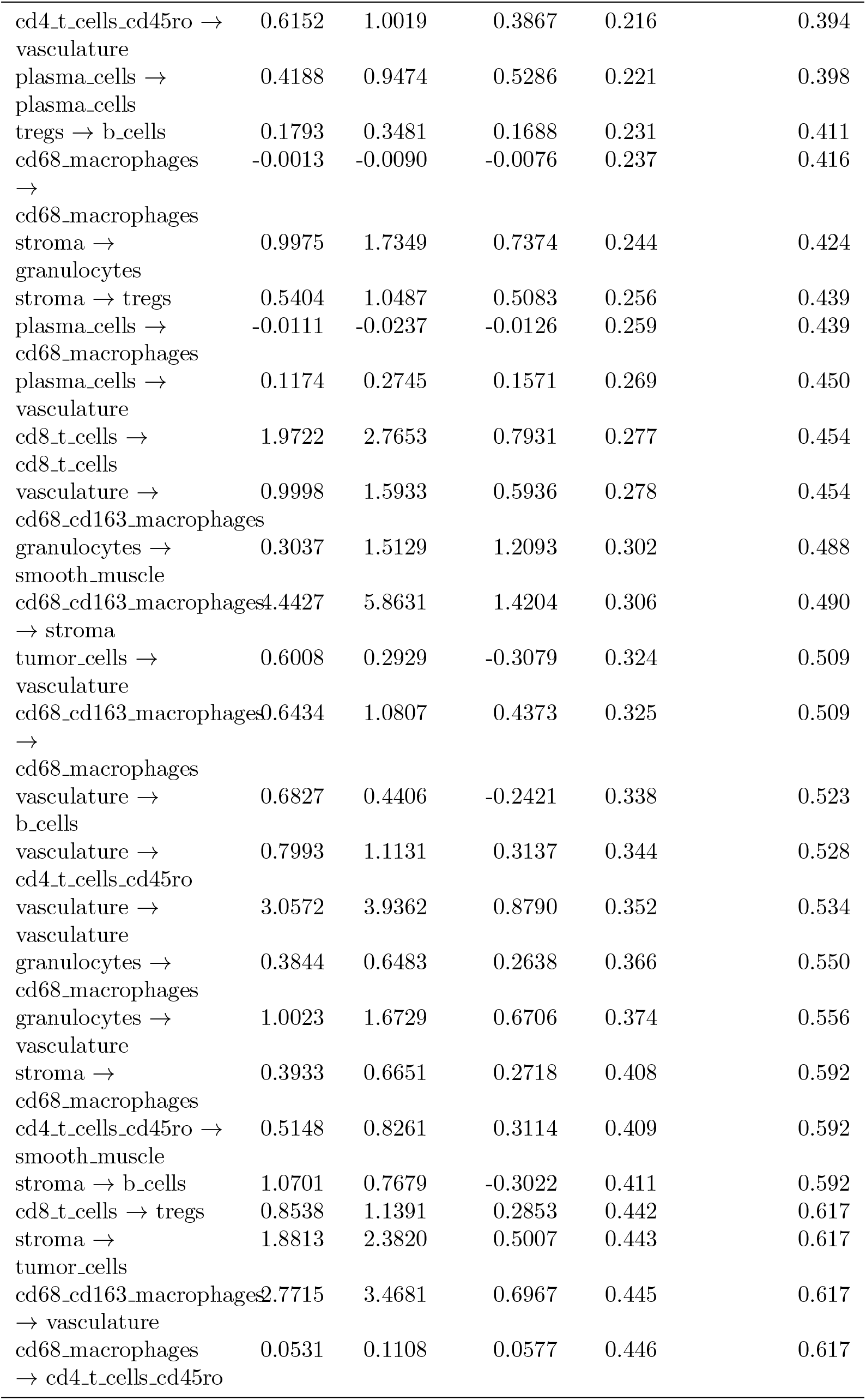
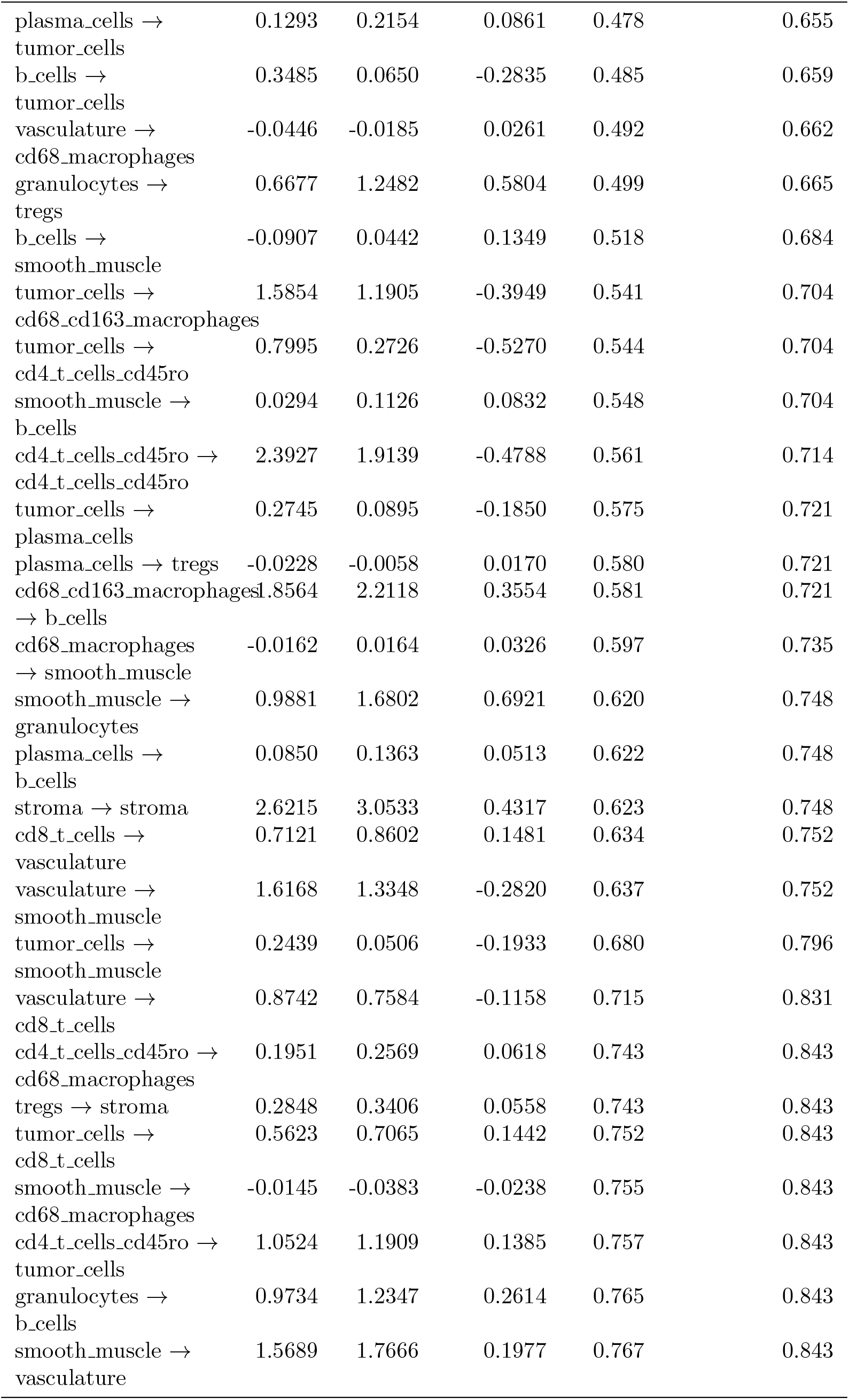
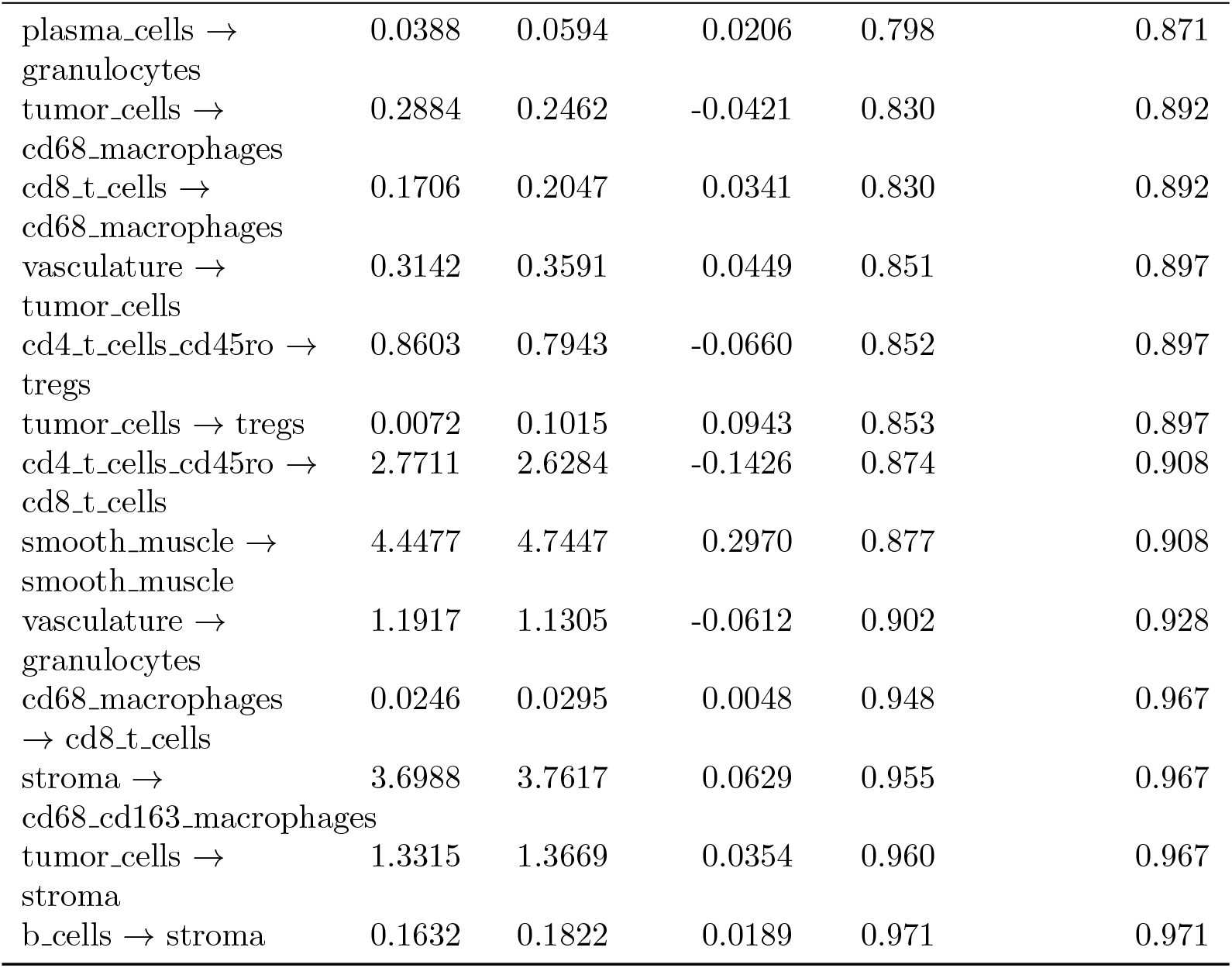
CRC TMA PANORAMIC results, using the neighborhood enrichment statistic at 25 micron.

#### S8.4 CRC runtime

**Table S2:** Runtime for the primary CRC PANORAMIC analysis on Stanford Sherlock HPC using 10 CPUs.

| Analysis step | Runtime (min) |
| --- | --- |
| Data preparation | 0.92 |
| Spatial statistic and bootstrap estimation | 1.37 |
| Multilevel meta-analysis | 3.67 |
| Total | 5.96 |

#### S8.5 CRC radius stability

**Figure S4:**
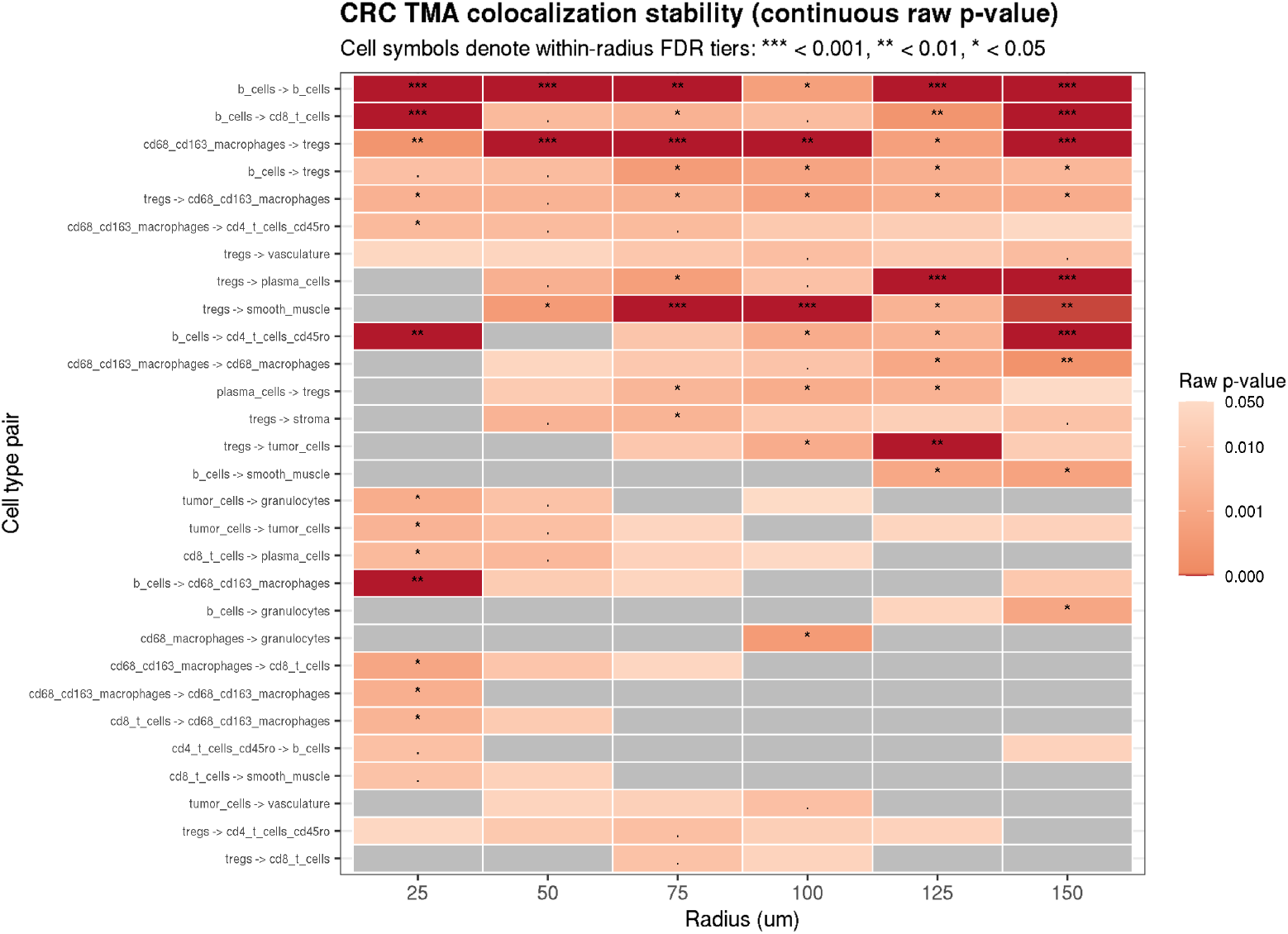
Stability of PANORAMIC results across different radii. The plotted cell pairs are those with results at or near statistical significance at any of the radii. Gray boxes indicate raw p-values ≥ 0.05, while shades of red indicate raw p-values achieving statistical significance *p <* 0.05. Asterisks indicate the p-values that retain statistical significance after performing Benjamini-Hochberg multiple testing correction.

#### S8.6 CRC spicyR benchmark

**Table S3:**
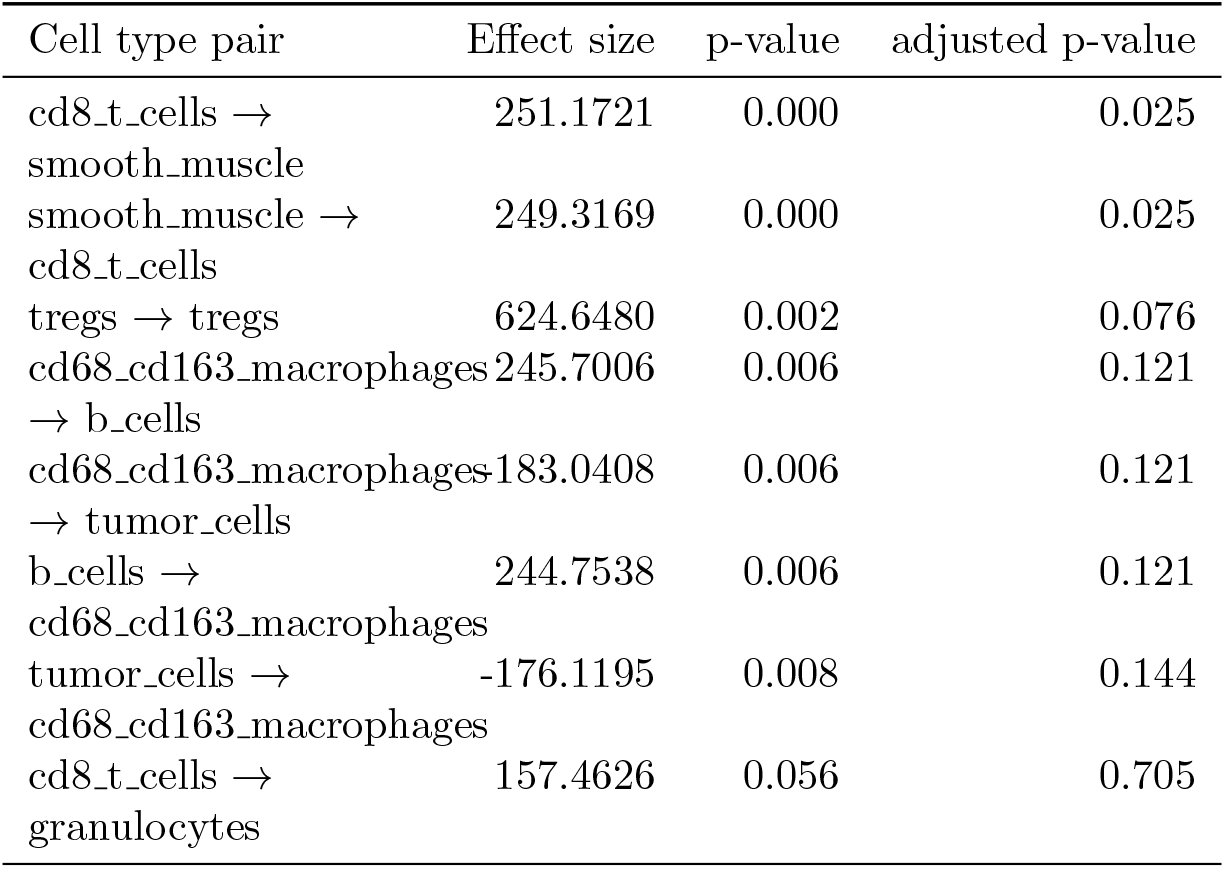
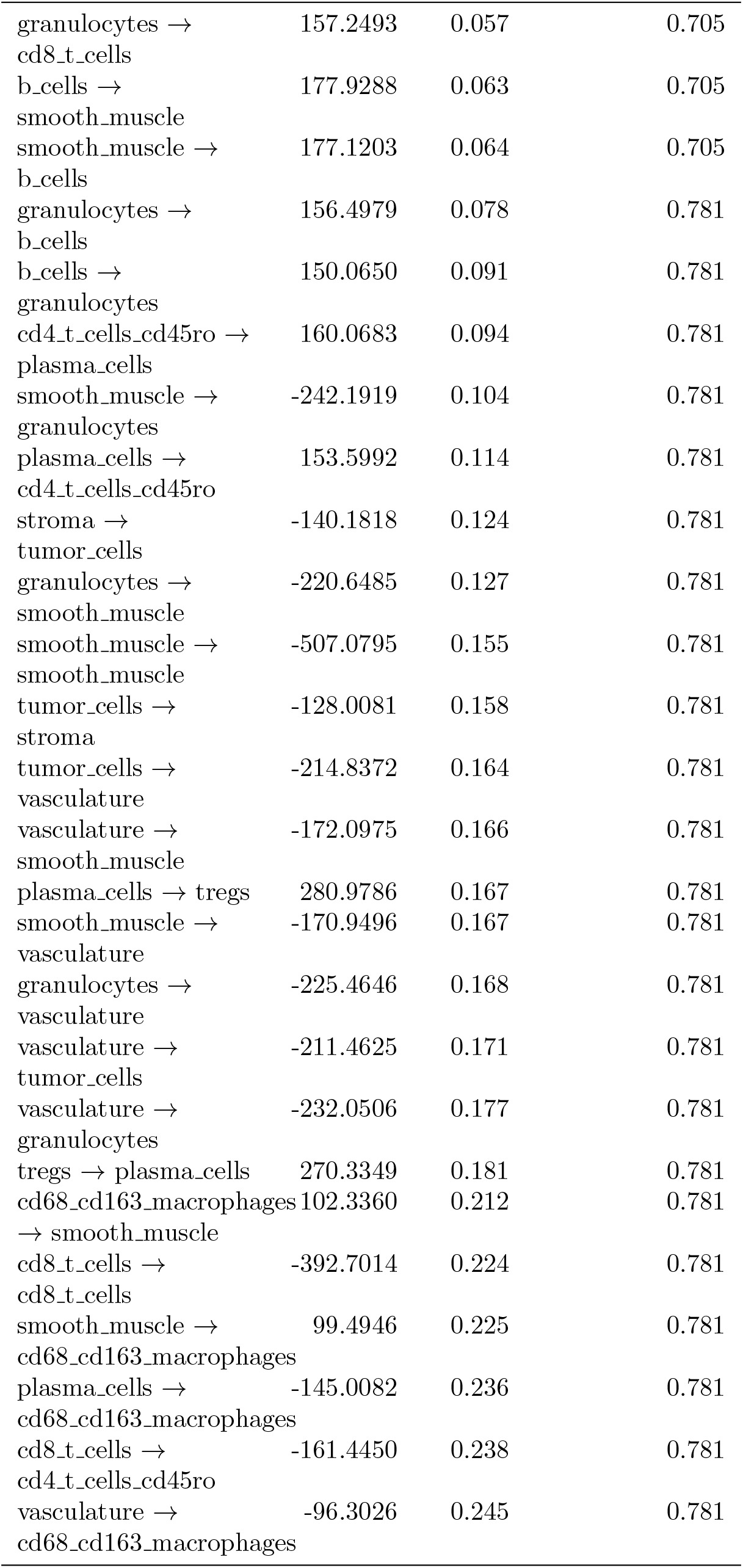
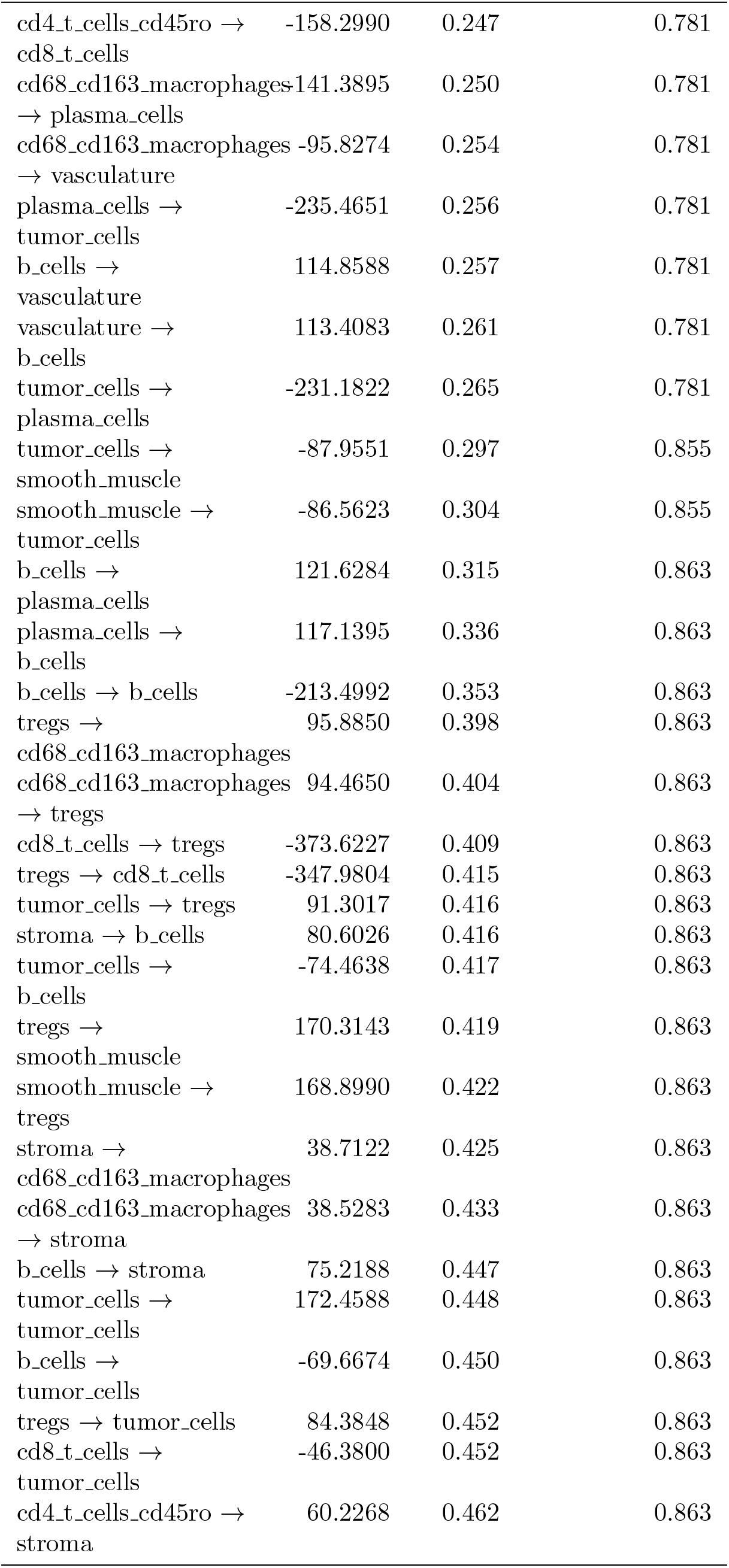
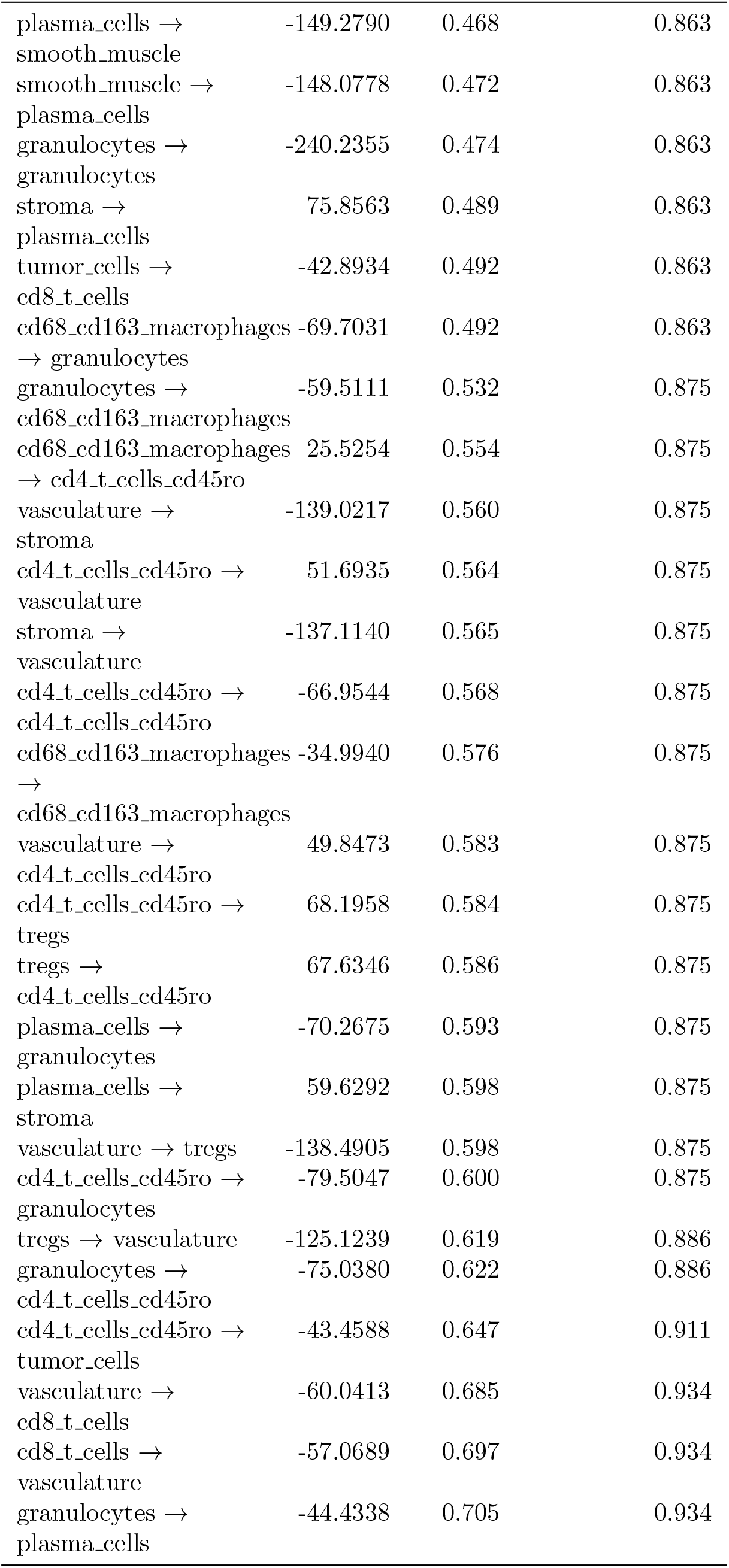
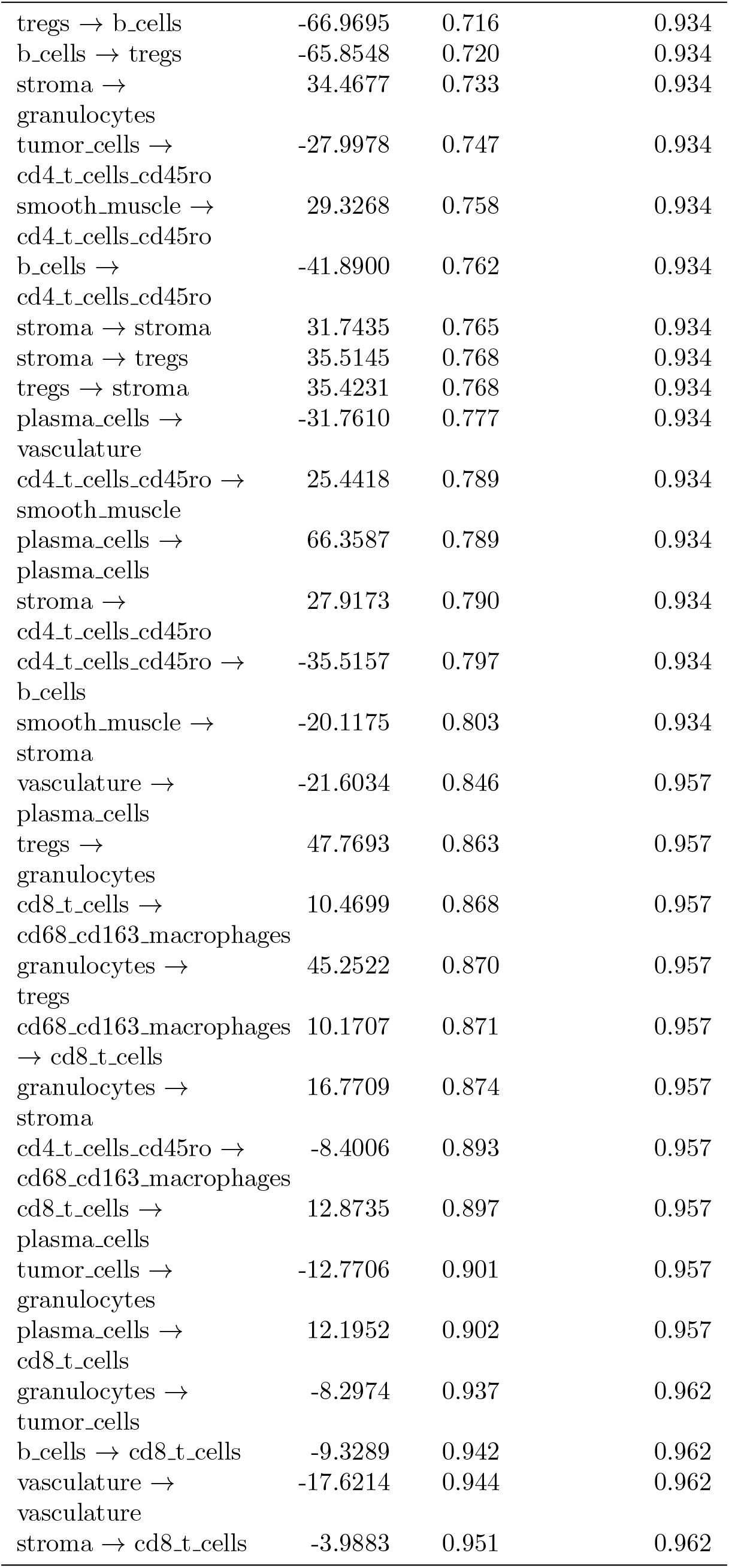
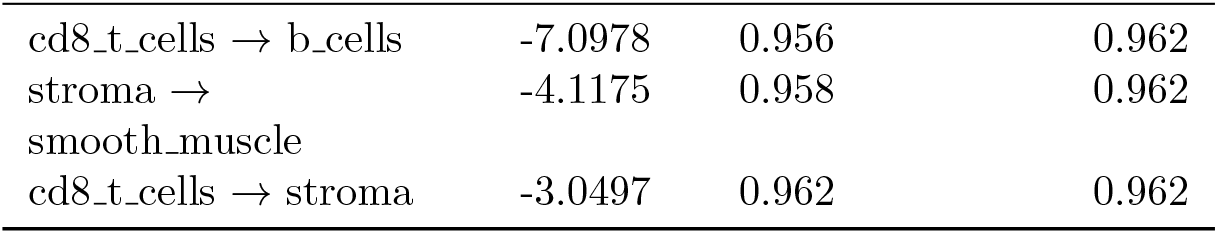
CRC TMA spicyR results (benchmark context).

#### S8.7 CRC Welch *t*-test benchmark

**Table S4:**
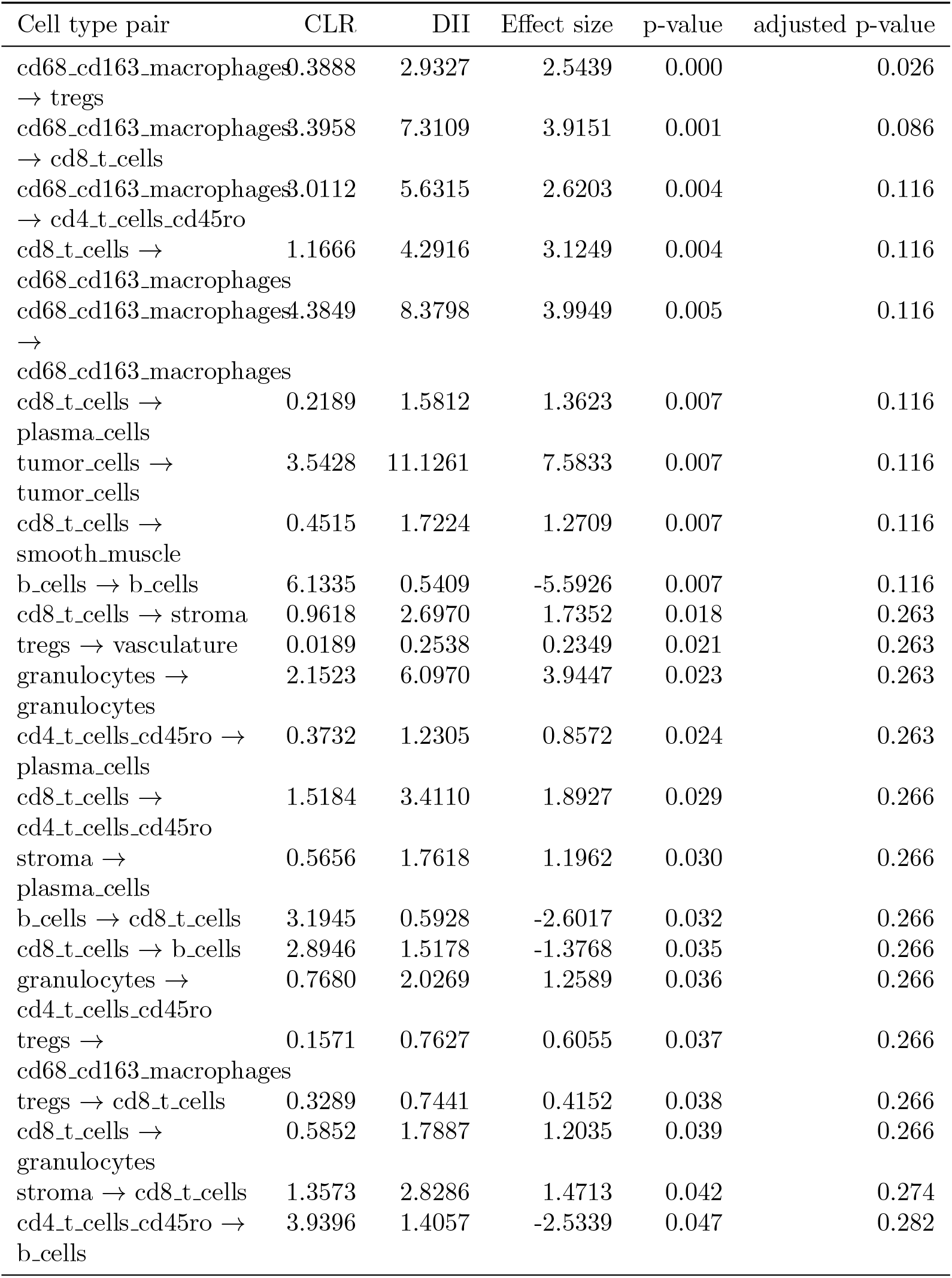
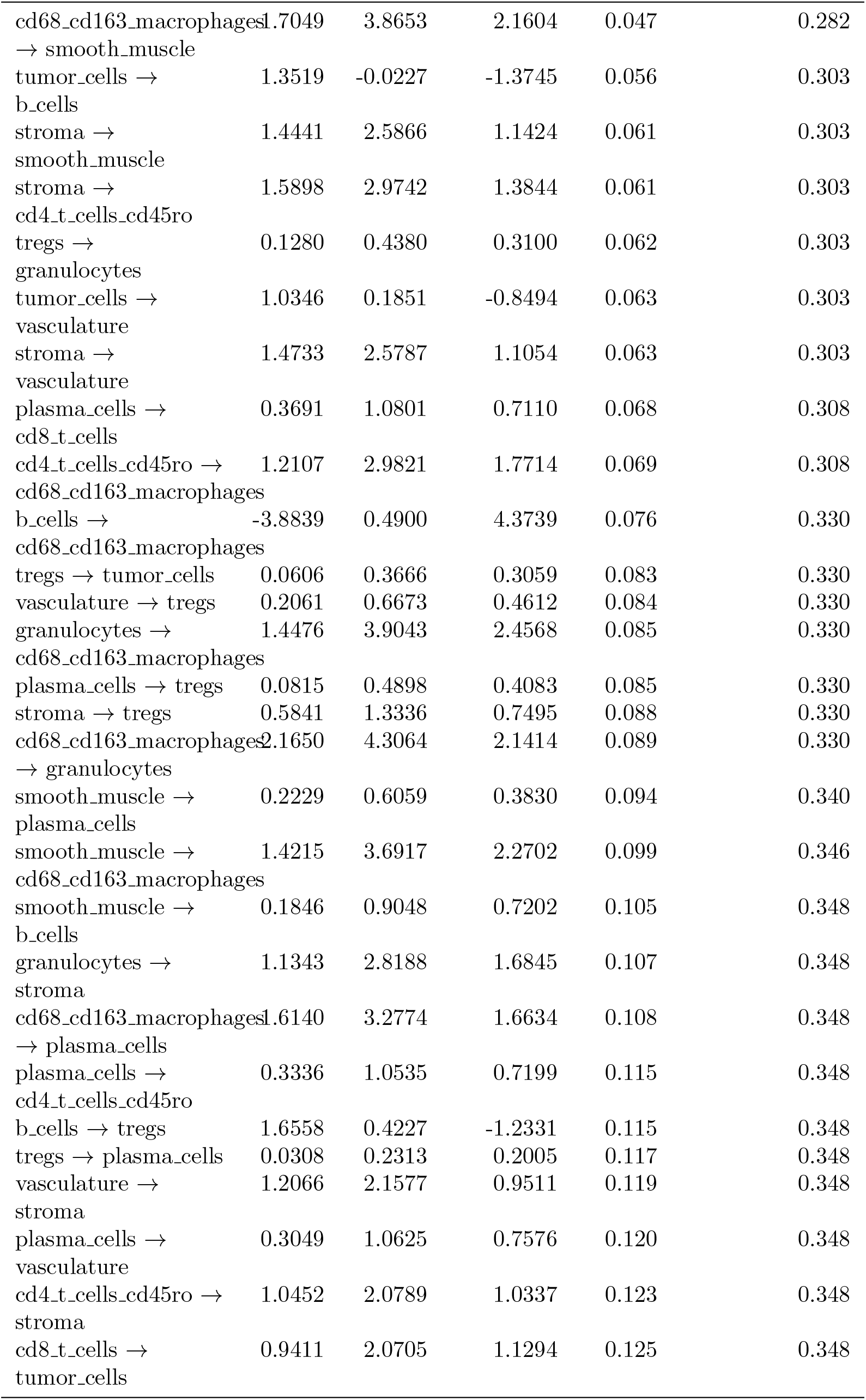
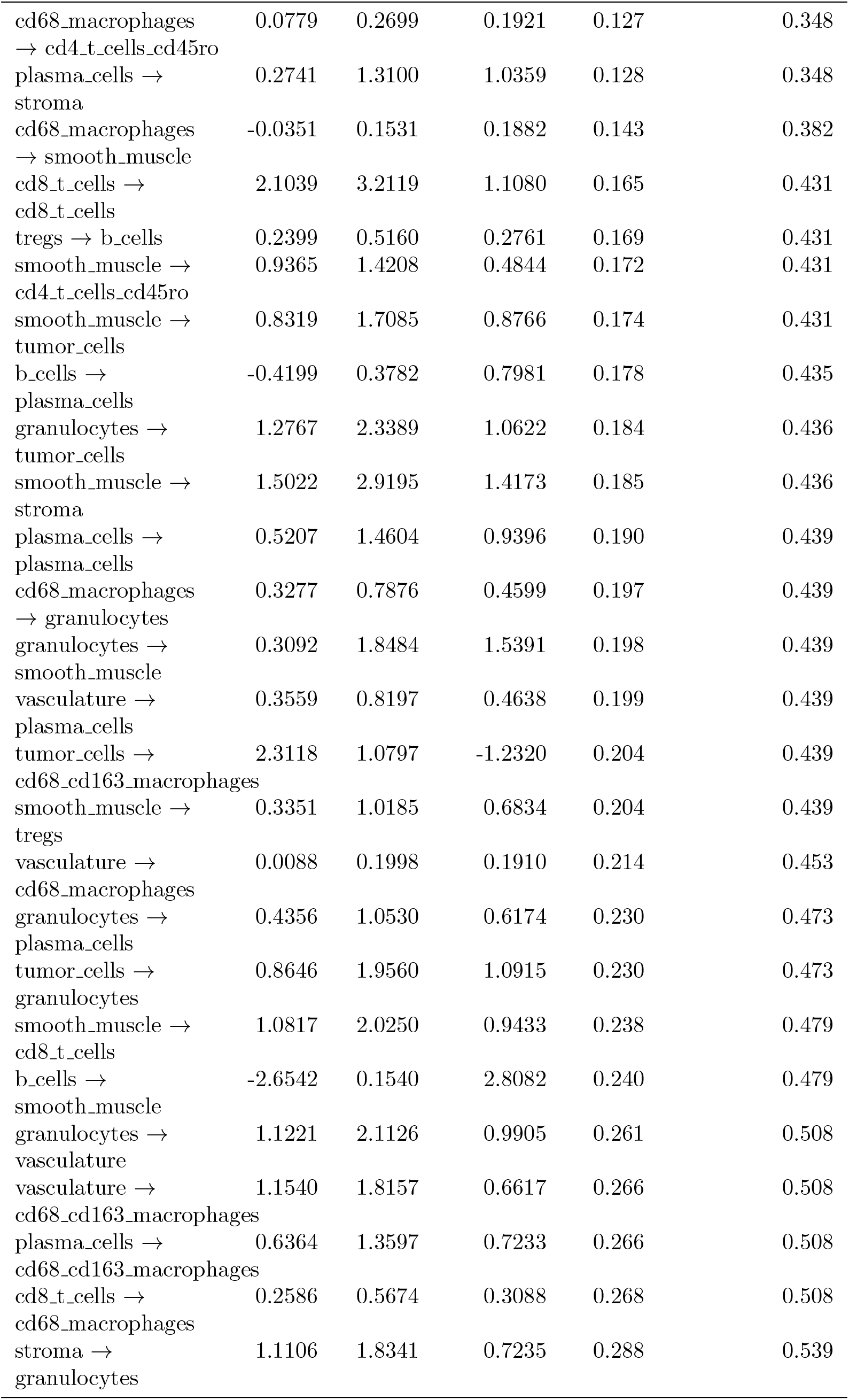
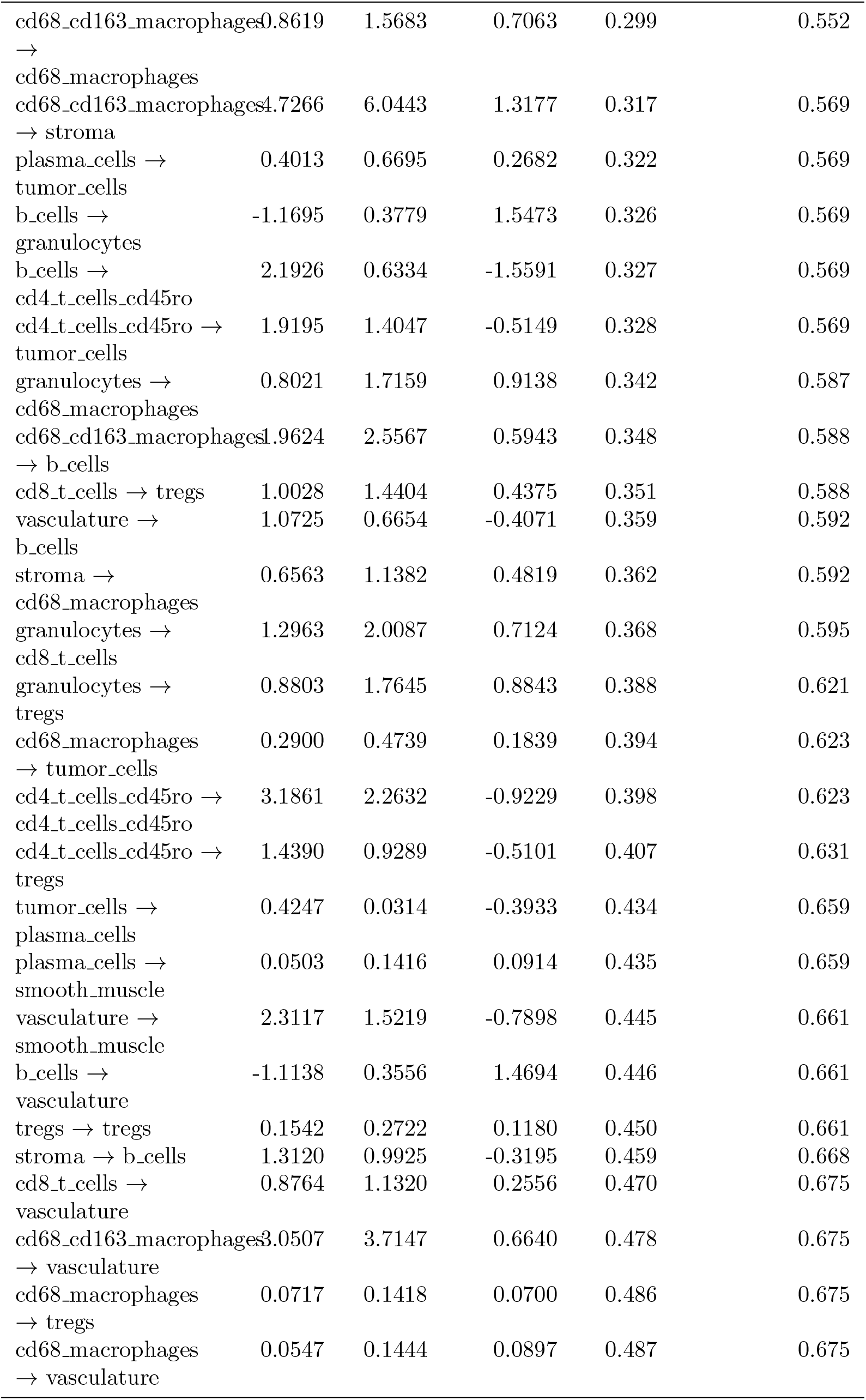
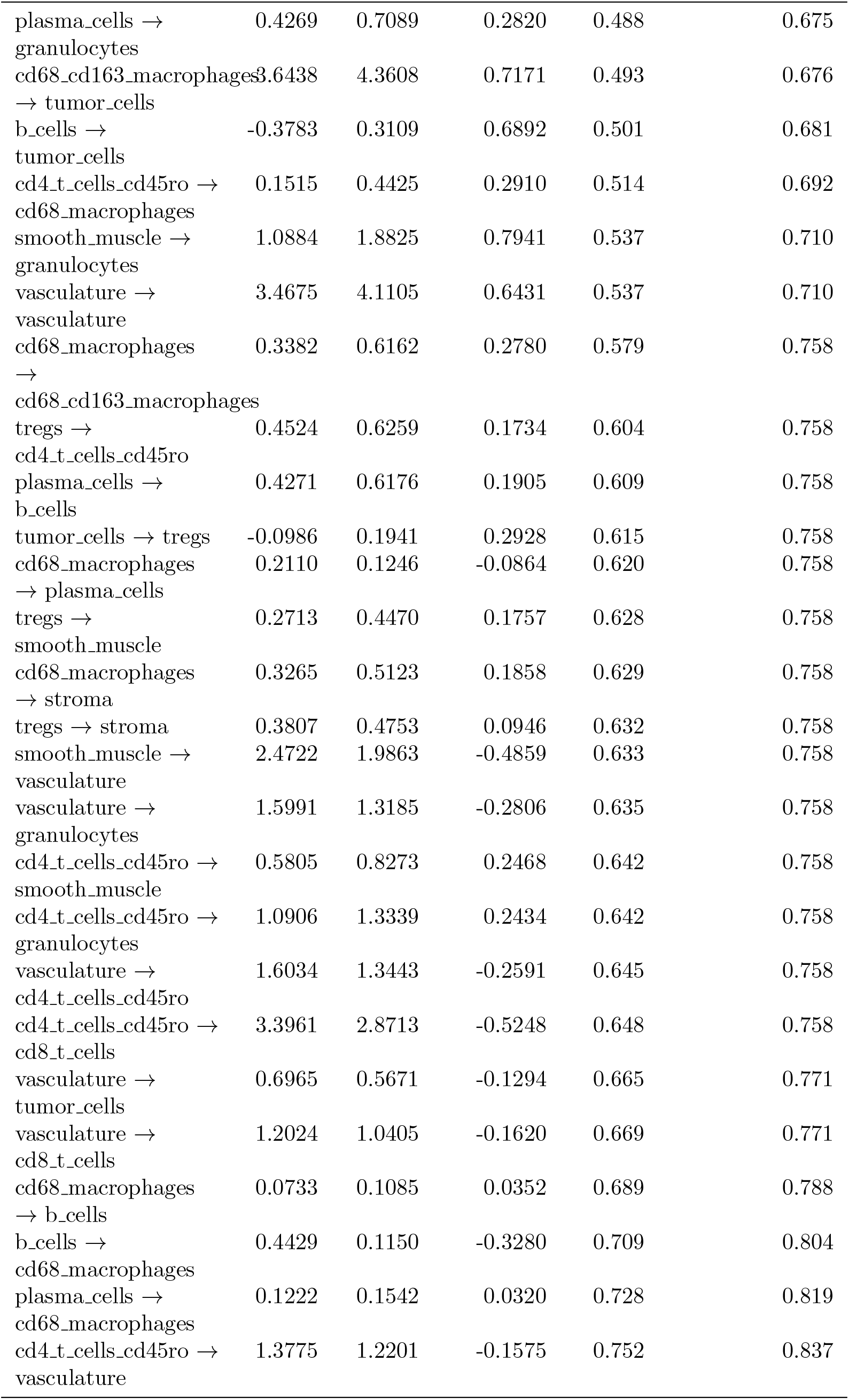
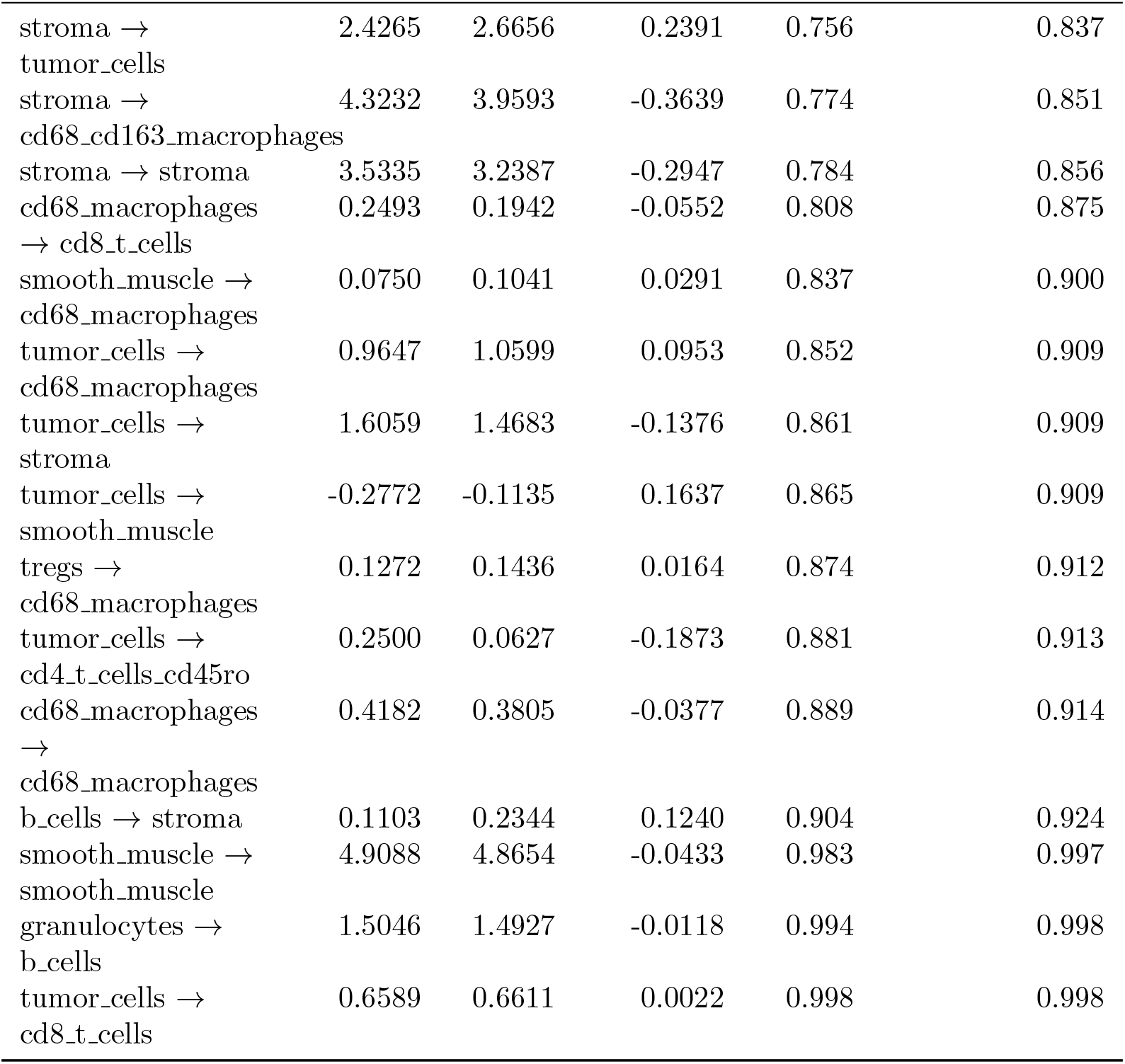
CRC TMA t-test results (benchmark context).

| Cell type pair | CLR | DII | Effect size | p-value | adjusted p-value |
| --- | --- | --- | --- | --- | --- |
| cd68_cd163_macrophages | 0.3888 | 2.9327 | 2.5439 | 0.000 | 0.026 |
| → tregs |  |  |  |  |  |
| cd68_cd163_macrophages | 3.3958 | 7.3109 | 3.9151 | 0.001 | 0.086 |
| → cd8.t.cells |  |  |  |  |  |
| cd68_cd163_macrophages | 3.0112 | 5.6315 | 2.6203 | 0.004 | 0.116 |
| → cd4.t.cells_cd45ro |  |  |  |  |  |
| cd8.t.cells → | 1.1666 | 4.2916 | 3.1249 | 0.004 | 0.116 |
| cd68_cd163_macrophages |  |  |  |  |  |
| cd68_cd163_macrophages | 4.3849 | 8.3798 | 3.9949 | 0.005 | 0.116 |
| → |  |  |  |  |  |
| cd68_cd163_macrophages |  |  |  |  |  |
| cd8.t.cells → | 0.2189 | 1.5812 | 1.3623 | 0.007 | 0.116 |
| plasma.cells |  |  |  |  |  |
| tumor.cells → | 3.5428 | 11.1261 | 7.5833 | 0.007 | 0.116 |
| tumor.cells |  |  |  |  |  |
| cd8.t.cells → | 0.4515 | 1.7224 | 1.2709 | 0.007 | 0.116 |
| smooth_muscle |  |  |  |  |  |
| b.cells → b.cells | 6.1335 | 0.5409 | -5.5926 | 0.007 | 0.116 |
| cd8.t.cells → stroma | 0.9618 | 2.6970 | 1.7352 | 0.018 | 0.263 |
| tregs → vasculature | 0.0189 | 0.2538 | 0.2349 | 0.021 | 0.263 |
| granulocytes → | 2.1523 | 6.0970 | 3.9447 | 0.023 | 0.263 |
| granulocytes |  |  |  |  |  |
| cd4.t.cells_cd45ro → | 0.3732 | 1.2305 | 0.8572 | 0.024 | 0.263 |
| plasma.cells |  |  |  |  |  |
| cd8.t.cells → | 1.5184 | 3.4110 | 1.8927 | 0.029 | 0.266 |
| cd4.t.cells_cd45ro |  |  |  |  |  |
| stroma → | 0.5656 | 1.7618 | 1.1962 | 0.030 | 0.266 |
| plasma.cells |  |  |  |  |  |
| b.cells → cd8.t.cells | 3.1945 | 0.5928 | -2.6017 | 0.032 | 0.266 |
| cd8.t.cells → b.cells | 2.8946 | 1.5178 | -1.3768 | 0.035 | 0.266 |
| granulocytes → | 0.7680 | 2.0269 | 1.2589 | 0.036 | 0.266 |
| cd4.t.cells_cd45ro |  |  |  |  |  |
| tregs → | 0.1571 | 0.7627 | 0.6055 | 0.037 | 0.266 |
| cd68_cd163_macrophages |  |  |  |  |  |
| tregs → cd8.t.cells | 0.3289 | 0.7441 | 0.4152 | 0.038 | 0.266 |
| cd8.t.cells → | 0.5852 | 1.7887 | 1.2035 | 0.039 | 0.266 |
| granulocytes |  |  |  |  |  |
| stroma → cd8.t.cells | 1.3573 | 2.8286 | 1.4713 | 0.042 | 0.274 |
| cd4.t.cells_cd45ro → | 3.9396 | 1.4057 | -2.5339 | 0.047 | 0.282 |
| b.cells |  |  |  |  |  |
| cd68_cd163_macrophages | 1.7049 | 3.8653 | 2.1604 | 0.047 | 0.282 |
| → smooth_muscle |  |  |  |  |  |
| tumor_cells → | 1.3519 | -0.0227 | -1.3745 | 0.056 | 0.303 |
| b_cells |  |  |  |  |  |
| stroma → | 1.4441 | 2.5866 | 1.1424 | 0.061 | 0.303 |
| smooth_muscle |  |  |  |  |  |
| stroma → | 1.5898 | 2.9742 | 1.3844 | 0.061 | 0.303 |
| cd4_t_cells_cd45ro |  |  |  |  |  |
| tregs → | 0.1280 | 0.4380 | 0.3100 | 0.062 | 0.303 |
| granulocytes |  |  |  |  |  |
| tumor_cells → | 1.0346 | 0.1851 | -0.8494 | 0.063 | 0.303 |
| vasculature |  |  |  |  |  |
| stroma → | 1.4733 | 2.5787 | 1.1054 | 0.063 | 0.303 |
| vasculature |  |  |  |  |  |
| plasma_cells → | 0.3691 | 1.0801 | 0.7110 | 0.068 | 0.308 |
| cd8_t_cells |  |  |  |  |  |
| cd4_t_cells_cd45ro → | 1.2107 | 2.9821 | 1.7714 | 0.069 | 0.308 |
| cd68_cd163_macrophages |  |  |  |  |  |
| b_cells → | -3.8839 | 0.4900 | 4.3739 | 0.076 | 0.330 |
| cd68_cd163_macrophages |  |  |  |  |  |
| tregs → tumor_cells | 0.0606 | 0.3666 | 0.3059 | 0.083 | 0.330 |
| vasculature → tregs | 0.2061 | 0.6673 | 0.4612 | 0.084 | 0.330 |
| granulocytes → | 1.4476 | 3.9043 | 2.4568 | 0.085 | 0.330 |
| cd68_cd163_macrophages |  |  |  |  |  |
| plasma_cells → tregs | 0.0815 | 0.4898 | 0.4083 | 0.085 | 0.330 |
| stroma → tregs | 0.5841 | 1.3336 | 0.7495 | 0.088 | 0.330 |
| cd68_cd163_macrophages | 2.1650 | 4.3064 | 2.1414 | 0.089 | 0.330 |
| → granulocytes |  |  |  |  |  |
| smooth_muscle → | 0.2229 | 0.6059 | 0.3830 | 0.094 | 0.340 |
| plasma_cells |  |  |  |  |  |
| smooth_muscle → | 1.4215 | 3.6917 | 2.2702 | 0.099 | 0.346 |
| cd68_cd163_macrophages |  |  |  |  |  |
| smooth_muscle → | 0.1846 | 0.9048 | 0.7202 | 0.105 | 0.348 |
| b_cells |  |  |  |  |  |
| granulocytes → | 1.1343 | 2.8188 | 1.6845 | 0.107 | 0.348 |
| stroma |  |  |  |  |  |
| cd68_cd163_macrophages | 1.6140 | 3.2774 | 1.6634 | 0.108 | 0.348 |
| → plasma_cells |  |  |  |  |  |
| plasma_cells → | 0.3336 | 1.0535 | 0.7199 | 0.115 | 0.348 |
| cd4_t_cells_cd45ro |  |  |  |  |  |
| b_cells → tregs | 1.6558 | 0.4227 | -1.2331 | 0.115 | 0.348 |
| tregs → plasma_cells | 0.0308 | 0.2313 | 0.2005 | 0.117 | 0.348 |
| vasculature → | 1.2066 | 2.1577 | 0.9511 | 0.119 | 0.348 |
| stroma |  |  |  |  |  |
| plasma_cells → | 0.3049 | 1.0625 | 0.7576 | 0.120 | 0.348 |
| vasculature |  |  |  |  |  |
| cd4_t_cells_cd45ro → | 1.0452 | 2.0789 | 1.0337 | 0.123 | 0.348 |
| stroma |  |  |  |  |  |
| cd8_t_cells → | 0.9411 | 2.0705 | 1.1294 | 0.125 | 0.348 |
| tumor_cells |  |  |  |  |  |
| cd68_macrophages<br>→ cd4_t_cells_cd45ro | 0.0779 | 0.2699 | 0.1921 | 0.127 | 0.348 |
| plasma_cells →<br>stroma | 0.2741 | 1.3100 | 1.0359 | 0.128 | 0.348 |
| cd68_macrophages<br>→ smooth_muscle | -0.0351 | 0.1531 | 0.1882 | 0.143 | 0.382 |
| cd8_t_cells →<br>cd8_t_cells | 2.1039 | 3.2119 | 1.1080 | 0.165 | 0.431 |
| tregs → b_cells | 0.2399 | 0.5160 | 0.2761 | 0.169 | 0.431 |
| smooth_muscle →<br>cd4_t_cells_cd45ro | 0.9365 | 1.4208 | 0.4844 | 0.172 | 0.431 |
| smooth_muscle →<br>tumor_cells | 0.8319 | 1.7085 | 0.8766 | 0.174 | 0.431 |
| b_cells →<br>plasma_cells | -0.4199 | 0.3782 | 0.7981 | 0.178 | 0.435 |
| granulocytes →<br>tumor_cells | 1.2767 | 2.3389 | 1.0622 | 0.184 | 0.436 |
| smooth_muscle →<br>stroma | 1.5022 | 2.9195 | 1.4173 | 0.185 | 0.436 |
| plasma_cells →<br>plasma_cells | 0.5207 | 1.4604 | 0.9396 | 0.190 | 0.439 |
| cd68_macrophages<br>→ granulocytes | 0.3277 | 0.7876 | 0.4599 | 0.197 | 0.439 |
| granulocytes →<br>smooth_muscle | 0.3092 | 1.8484 | 1.5391 | 0.198 | 0.439 |
| vasculature →<br>plasma_cells | 0.3559 | 0.8197 | 0.4638 | 0.199 | 0.439 |
| tumor_cells →<br>cd68_cd163_macrophages | 2.3118 | 1.0797 | -1.2320 | 0.204 | 0.439 |
| smooth_muscle →<br>tregs | 0.3351 | 1.0185 | 0.6834 | 0.204 | 0.439 |
| vasculature →<br>cd68_macrophages | 0.0088 | 0.1998 | 0.1910 | 0.214 | 0.453 |
| granulocytes →<br>plasma_cells | 0.4356 | 1.0530 | 0.6174 | 0.230 | 0.473 |
| tumor_cells →<br>granulocytes | 0.8646 | 1.9560 | 1.0915 | 0.230 | 0.473 |
| smooth_muscle →<br>cd8_t_cells | 1.0817 | 2.0250 | 0.9433 | 0.238 | 0.479 |
| b_cells →<br>smooth_muscle | -2.6542 | 0.1540 | 2.8082 | 0.240 | 0.479 |
| granulocytes →<br>vasculature | 1.1221 | 2.1126 | 0.9905 | 0.261 | 0.508 |
| vasculature →<br>cd68_cd163_macrophages | 1.1540 | 1.8157 | 0.6617 | 0.266 | 0.508 |
| plasma_cells →<br>cd68_cd163_macrophages | 0.6364 | 1.3597 | 0.7233 | 0.266 | 0.508 |
| cd8_t_cells →<br>cd68_macrophages | 0.2586 | 0.5674 | 0.3088 | 0.268 | 0.508 |
| stroma →<br>granulocytes | 1.1106 | 1.8341 | 0.7235 | 0.288 | 0.539 |
| cd68_cd163_macrophages | 0.8619 | 1.5683 | 0.7063 | 0.299 | 0.552 |
| →<br>cd68_macrophages |  |  |  |  |  |
| cd68_cd163_macrophages | 4.7266 | 6.0443 | 1.3177 | 0.317 | 0.569 |
| → stroma |  |  |  |  |  |
| plasma_cells →<br>tumor_cells | 0.4013 | 0.6695 | 0.2682 | 0.322 | 0.569 |
| b_cells →<br>granulocytes | -1.1695 | 0.3779 | 1.5473 | 0.326 | 0.569 |
| b_cells →<br>cd4_t_cells_cd45ro | 2.1926 | 0.6334 | -1.5591 | 0.327 | 0.569 |
| cd4_t_cells_cd45ro →<br>tumor_cells | 1.9195 | 1.4047 | -0.5149 | 0.328 | 0.569 |
| granulocytes →<br>cd68_macrophages | 0.8021 | 1.7159 | 0.9138 | 0.342 | 0.587 |
| cd68_cd163_macrophages | 1.9624 | 2.5567 | 0.5943 | 0.348 | 0.588 |
| → b_cells |  |  |  |  |  |
| cd8_t_cells → tregs | 1.0028 | 1.4404 | 0.4375 | 0.351 | 0.588 |
| vasculature →<br>b_cells | 1.0725 | 0.6654 | -0.4071 | 0.359 | 0.592 |
| stroma →<br>cd68_macrophages | 0.6563 | 1.1382 | 0.4819 | 0.362 | 0.592 |
| granulocytes →<br>cd8_t_cells | 1.2963 | 2.0087 | 0.7124 | 0.368 | 0.595 |
| granulocytes →<br>tregs | 0.8803 | 1.7645 | 0.8843 | 0.388 | 0.621 |
| cd68_macrophages<br>→ tumor_cells | 0.2900 | 0.4739 | 0.1839 | 0.394 | 0.623 |
| cd4_t_cells_cd45ro →<br>cd4_t_cells_cd45ro | 3.1861 | 2.2632 | -0.9229 | 0.398 | 0.623 |
| cd4_t_cells_cd45ro →<br>tregs | 1.4390 | 0.9289 | -0.5101 | 0.407 | 0.631 |
| tumor_cells →<br>plasma_cells | 0.4247 | 0.0314 | -0.3933 | 0.434 | 0.659 |
| plasma_cells →<br>smooth_muscle | 0.0503 | 0.1416 | 0.0914 | 0.435 | 0.659 |
| vasculature →<br>smooth_muscle | 2.3117 | 1.5219 | -0.7898 | 0.445 | 0.661 |
| b_cells →<br>vasculature | -1.1138 | 0.3556 | 1.4694 | 0.446 | 0.661 |
| tregs → tregs | 0.1542 | 0.2722 | 0.1180 | 0.450 | 0.661 |
| stroma → b_cells | 1.3120 | 0.9925 | -0.3195 | 0.459 | 0.668 |
| cd8_t_cells →<br>vasculature | 0.8764 | 1.1320 | 0.2556 | 0.470 | 0.675 |
| cd68_cd163_macrophages | 3.0507 | 3.7147 | 0.6640 | 0.478 | 0.675 |
| → vasculature |  |  |  |  |  |
| cd68_macrophages<br>→ tregs | 0.0717 | 0.1418 | 0.0700 | 0.486 | 0.675 |
| cd68_macrophages<br>→ vasculature | 0.0547 | 0.1444 | 0.0897 | 0.487 | 0.675 |
| plasma_cells →<br>granulocytes | 0.4269 | 0.7089 | 0.2820 | 0.488 | 0.675 |
| cd68_cd163_macrophages → tumor_cells | 3.6438 | 4.3608 | 0.7171 | 0.493 | 0.676 |
| b_cells →<br>tumor_cells | -0.3783 | 0.3109 | 0.6892 | 0.501 | 0.681 |
| cd4_t_cells_cd45ro →<br>cd68_macrophages | 0.1515 | 0.4425 | 0.2910 | 0.514 | 0.692 |
| smooth_muscle →<br>granulocytes | 1.0884 | 1.8825 | 0.7941 | 0.537 | 0.710 |
| vasculature →<br>vasculature | 3.4675 | 4.1105 | 0.6431 | 0.537 | 0.710 |
| cd68_macrophages →<br>cd68_cd163_macrophages | 0.3382 | 0.6162 | 0.2780 | 0.579 | 0.758 |
| tregs →<br>cd4_t_cells_cd45ro | 0.4524 | 0.6259 | 0.1734 | 0.604 | 0.758 |
| plasma_cells →<br>b_cells | 0.4271 | 0.6176 | 0.1905 | 0.609 | 0.758 |
| tumor_cells → tregs | -0.0986 | 0.1941 | 0.2928 | 0.615 | 0.758 |
| cd68_macrophages → plasma_cells | 0.2110 | 0.1246 | -0.0864 | 0.620 | 0.758 |
| tregs →<br>smooth_muscle | 0.2713 | 0.4470 | 0.1757 | 0.628 | 0.758 |
| cd68_macrophages → stroma | 0.3265 | 0.5123 | 0.1858 | 0.629 | 0.758 |
| tregs → stroma | 0.3807 | 0.4753 | 0.0946 | 0.632 | 0.758 |
| smooth_muscle →<br>vasculature | 2.4722 | 1.9863 | -0.4859 | 0.633 | 0.758 |
| vasculature →<br>granulocytes | 1.5991 | 1.3185 | -0.2806 | 0.635 | 0.758 |
| cd4_t_cells_cd45ro →<br>smooth_muscle | 0.5805 | 0.8273 | 0.2468 | 0.642 | 0.758 |
| cd4_t_cells_cd45ro →<br>granulocytes | 1.0906 | 1.3339 | 0.2434 | 0.642 | 0.758 |
| vasculature →<br>cd4_t_cells_cd45ro | 1.6034 | 1.3443 | -0.2591 | 0.645 | 0.758 |
| cd4_t_cells_cd45ro →<br>cd8_t_cells | 3.3961 | 2.8713 | -0.5248 | 0.648 | 0.758 |
| vasculature →<br>tumor_cells | 0.6965 | 0.5671 | -0.1294 | 0.665 | 0.771 |
| vasculature →<br>cd8_t_cells | 1.2024 | 1.0405 | -0.1620 | 0.669 | 0.771 |
| cd68_macrophages → b_cells | 0.0733 | 0.1085 | 0.0352 | 0.689 | 0.788 |
| b_cells →<br>cd68_macrophages | 0.4429 | 0.1150 | -0.3280 | 0.709 | 0.804 |
| plasma_cells →<br>cd68_macrophages | 0.1222 | 0.1542 | 0.0320 | 0.728 | 0.819 |
| cd4_t_cells_cd45ro →<br>vasculature | 1.3775 | 1.2201 | -0.1575 | 0.752 | 0.837 |
| stroma → tumor_cells | 2.4265 | 2.6656 | 0.2391 | 0.756 | 0.837 |
| stroma → cd68_cd163_macrophages | 4.3232 | 3.9593 | -0.3639 | 0.774 | 0.851 |
| stroma → stroma | 3.5335 | 3.2387 | -0.2947 | 0.784 | 0.856 |
| cd68_macrophages → cd8_t_cells | 0.2493 | 0.1942 | -0.0552 | 0.808 | 0.875 |
| smooth_muscle → cd68_macrophages | 0.0750 | 0.1041 | 0.0291 | 0.837 | 0.900 |
| tumor_cells → cd68_macrophages | 0.9647 | 1.0599 | 0.0953 | 0.852 | 0.909 |
| tumor_cells → stroma | 1.6059 | 1.4683 | -0.1376 | 0.861 | 0.909 |
| tumor_cells → smooth_muscle | -0.2772 | -0.1135 | 0.1637 | 0.865 | 0.909 |
| tregs → cd68_macrophages | 0.1272 | 0.1436 | 0.0164 | 0.874 | 0.912 |
| tumor_cells → cd4_t_cells_cd45ro | 0.2500 | 0.0627 | -0.1873 | 0.881 | 0.913 |
| cd68_macrophages → cd68_macrophages | 0.4182 | 0.3805 | -0.0377 | 0.889 | 0.914 |
| b_cells → stroma | 0.1103 | 0.2344 | 0.1240 | 0.904 | 0.924 |
| smooth_muscle → smooth_muscle | 4.9088 | 4.8654 | -0.0433 | 0.983 | 0.997 |
| granulocytes → b_cells | 1.5046 | 1.4927 | -0.0118 | 0.994 | 0.998 |
| tumor_cells → cd8_t_cells | 0.6589 | 0.6611 | 0.0022 | 0.998 | 0.998 |

#### S8.8 CRC *p*-value distribution comparison

**Figure S5:**
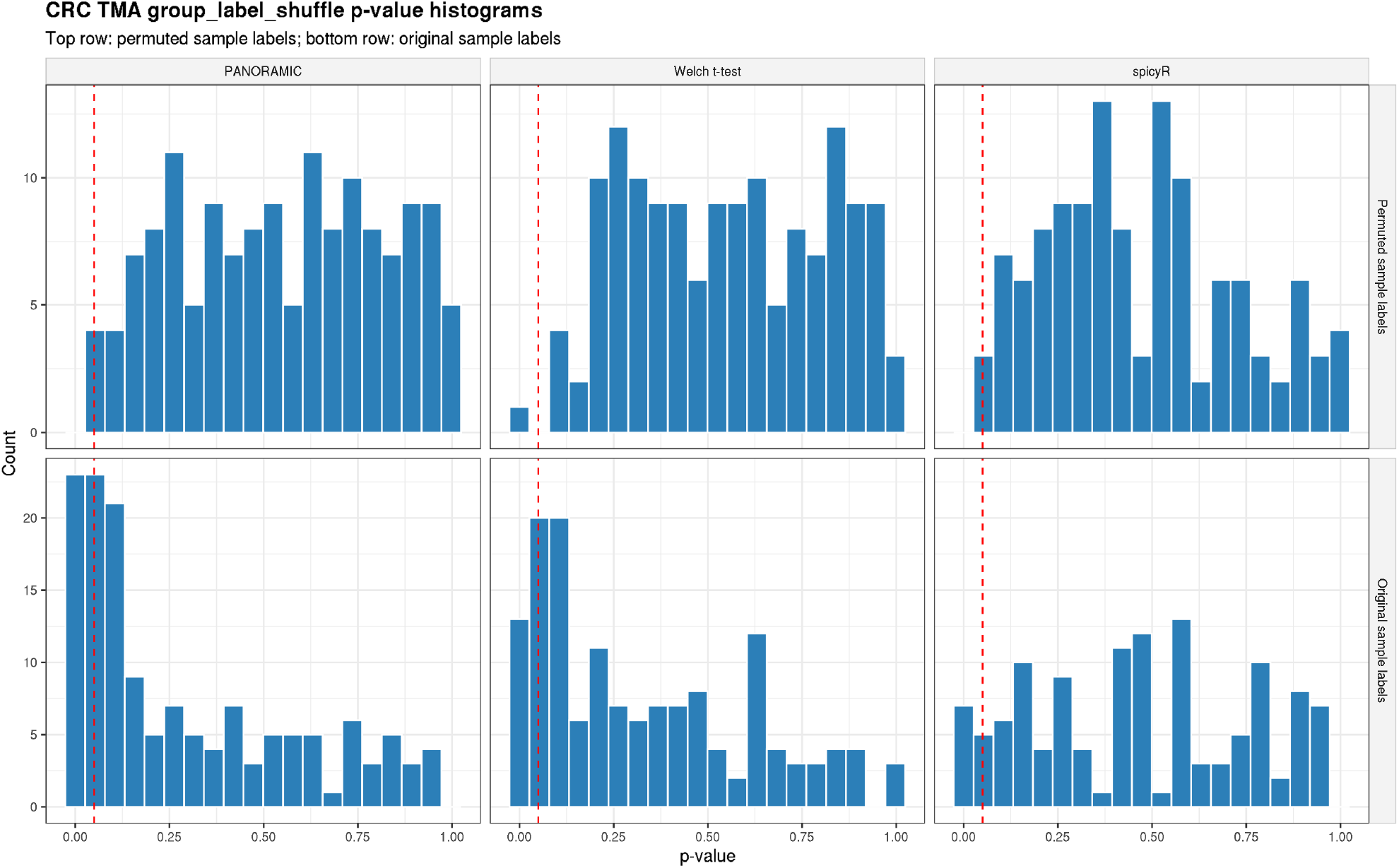
P-value distributions for CRC differential colocalization analyses at *r* = 25 *µ*m. Top row: p-value histograms of cell-type pair colocalization when sample labels (CLR or DII) are permuted. Bottom row: p-value histograms with correctly labeled data. PANORAMIC has improved sensitivity to colocalization differences over t-testing and spicyR.

#### S8.9 Alternative spatial statistic: cross-*L*

**Table S5:**
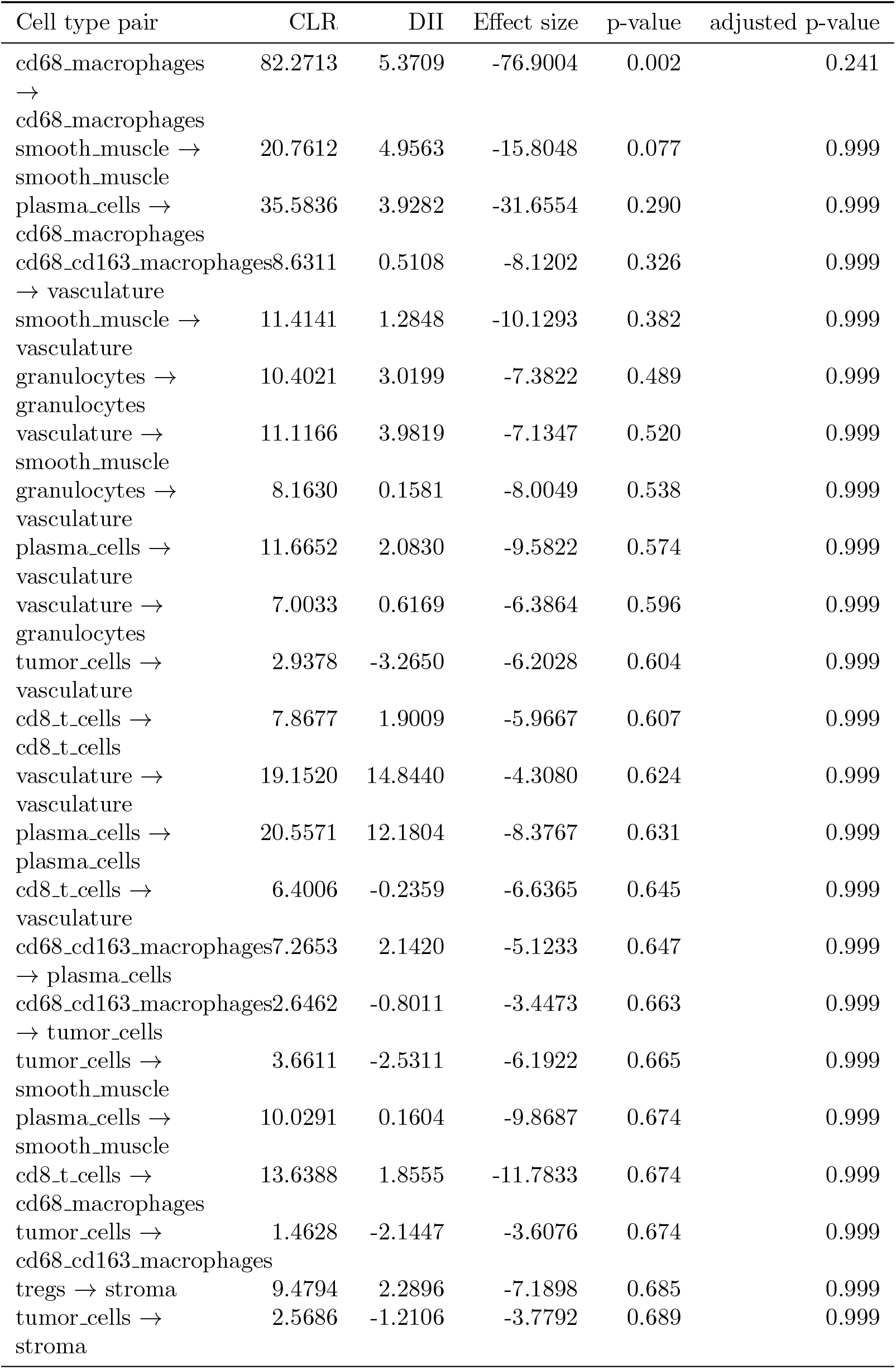
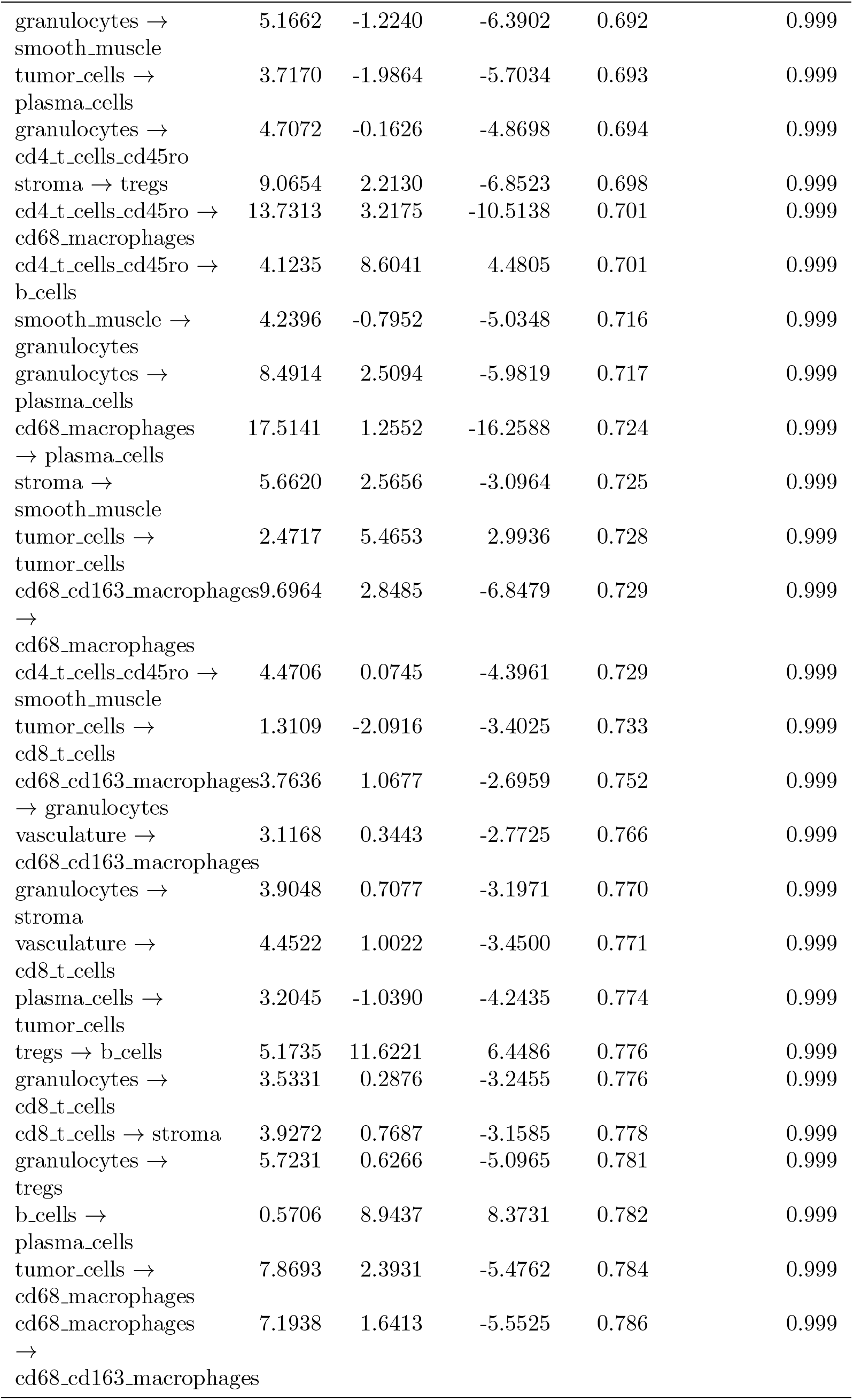
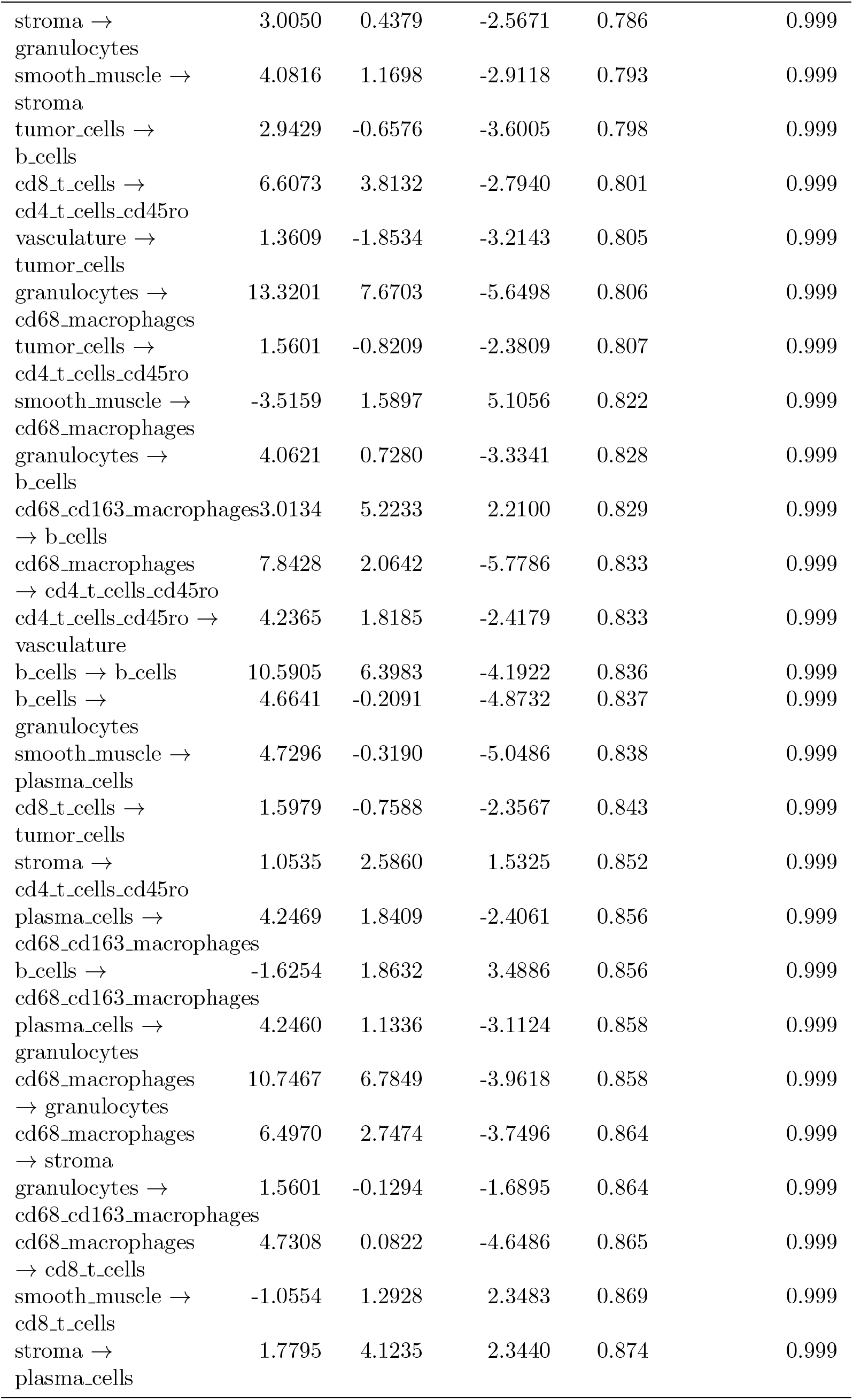
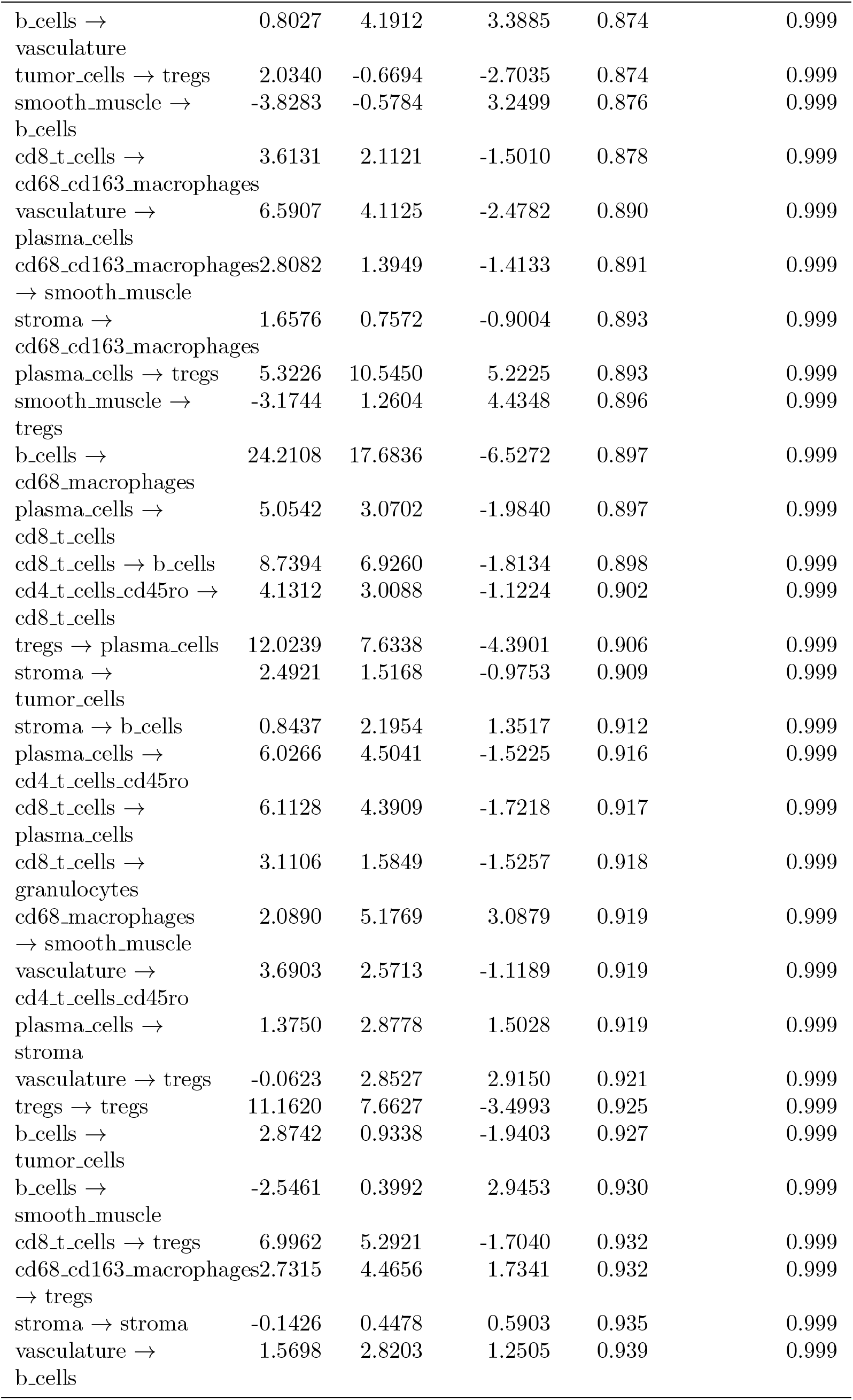
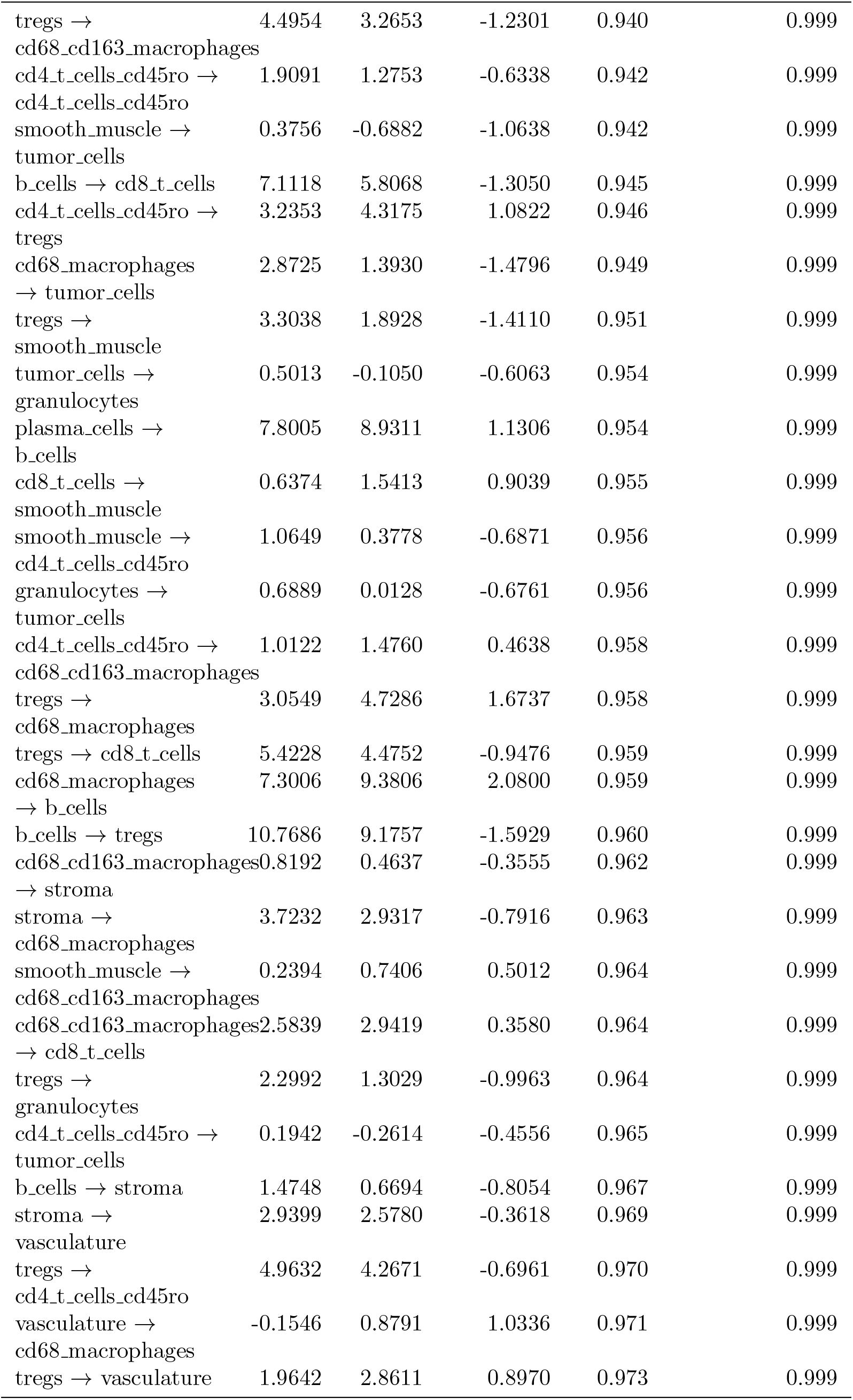
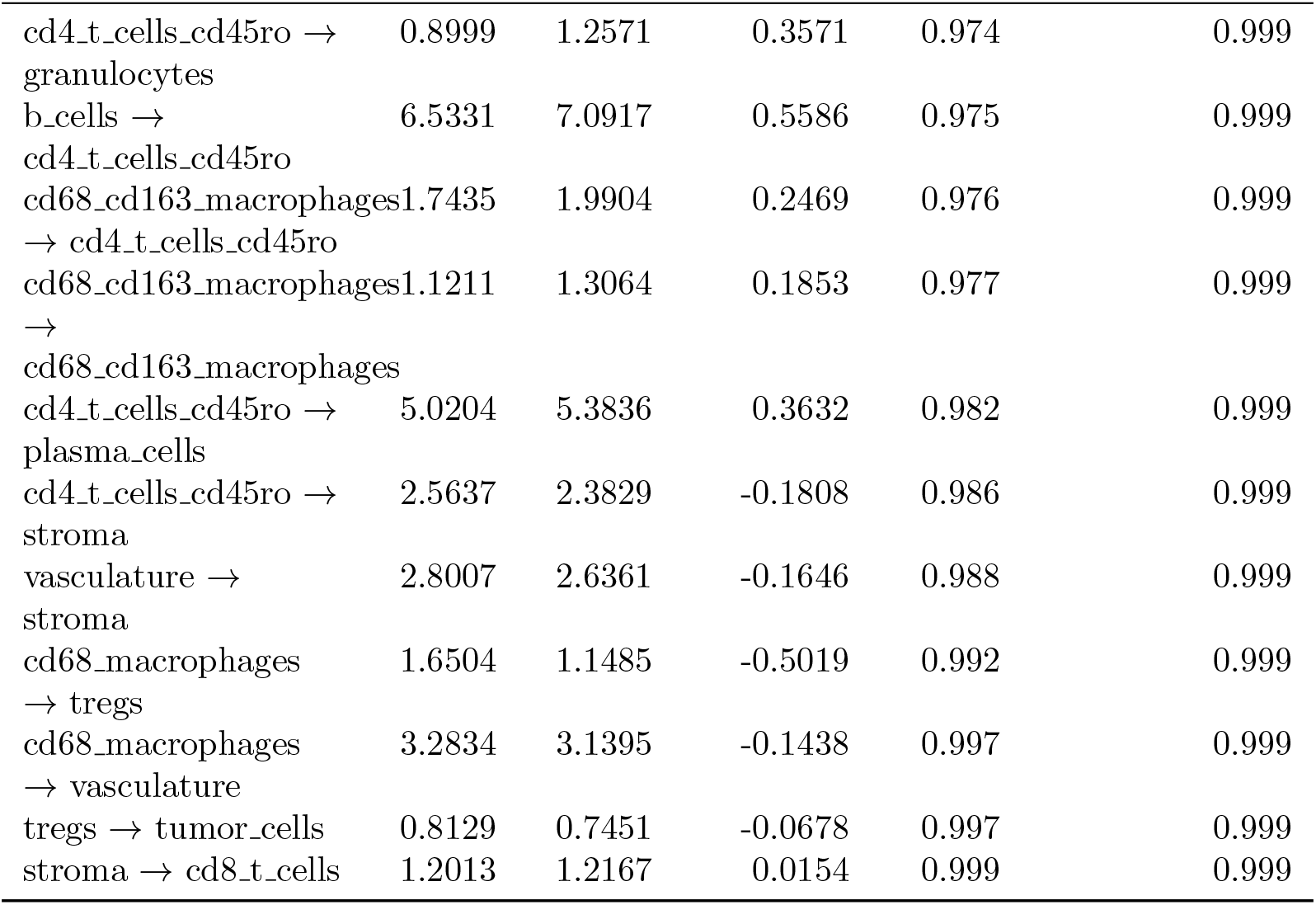
CRC TMA PANORAMIC results.

| Cell type pair | CLR | DII | Effect size | p-value | adjusted p-value |
| --- | --- | --- | --- | --- | --- |
| cd68_macrophages<br>→<br>cd68_macrophages | 82.2713 | 5.3709 | -76.9004 | 0.002 | 0.241 |
| smooth_muscle →<br>smooth_muscle | 20.7612 | 4.9563 | -15.8048 | 0.077 | 0.999 |
| plasma_cells →<br>cd68_macrophages | 35.5836 | 3.9282 | -31.6554 | 0.290 | 0.999 |
| cd68_cd163_macrophages<br>→ vasculature | 8.6311 | 0.5108 | -8.1202 | 0.326 | 0.999 |
| smooth_muscle →<br>vasculature | 11.4141 | 1.2848 | -10.1293 | 0.382 | 0.999 |
| granulocytes →<br>granulocytes | 10.4021 | 3.0199 | -7.3822 | 0.489 | 0.999 |
| vasculature →<br>smooth_muscle | 11.1166 | 3.9819 | -7.1347 | 0.520 | 0.999 |
| granulocytes →<br>vasculature | 8.1630 | 0.1581 | -8.0049 | 0.538 | 0.999 |
| plasma_cells →<br>vasculature | 11.6652 | 2.0830 | -9.5822 | 0.574 | 0.999 |
| vasculature →<br>granulocytes | 7.0033 | 0.6169 | -6.3864 | 0.596 | 0.999 |
| tumor_cells →<br>vasculature | 2.9378 | -3.2650 | -6.2028 | 0.604 | 0.999 |
| cd8_t_cells →<br>cd8_t_cells | 7.8677 | 1.9009 | -5.9667 | 0.607 | 0.999 |
| vasculature →<br>vasculature | 19.1520 | 14.8440 | -4.3080 | 0.624 | 0.999 |
| plasma_cells →<br>plasma_cells | 20.5571 | 12.1804 | -8.3767 | 0.631 | 0.999 |
| cd8_t_cells →<br>vasculature | 6.4006 | -0.2359 | -6.6365 | 0.645 | 0.999 |
| cd68_cd163_macrophages<br>→ plasma_cells | 7.2653 | 2.1420 | -5.1233 | 0.647 | 0.999 |
| cd68_cd163_macrophages<br>→ tumor_cells | 2.6462 | -0.8011 | -3.4473 | 0.663 | 0.999 |
| tumor_cells →<br>smooth_muscle | 3.6611 | -2.5311 | -6.1922 | 0.665 | 0.999 |
| plasma_cells →<br>smooth_muscle | 10.0291 | 0.1604 | -9.8687 | 0.674 | 0.999 |
| cd8_t_cells →<br>cd68_macrophages | 13.6388 | 1.8555 | -11.7833 | 0.674 | 0.999 |
| tumor_cells →<br>cd68_cd163_macrophages | 1.4628 | -2.1447 | -3.6076 | 0.674 | 0.999 |
| tregs → stroma | 9.4794 | 2.2896 | -7.1898 | 0.685 | 0.999 |
| tumor_cells →<br>stroma | 2.5686 | -1.2106 | -3.7792 | 0.689 | 0.999 |
| granulocytes → smooth_muscle | 5.1662 | -1.2240 | -6.3902 | 0.692 | 0.999 |
| tumor_cells → plasma_cells | 3.7170 | -1.9864 | -5.7034 | 0.693 | 0.999 |
| granulocytes → cd4_t_cells_cd45ro | 4.7072 | -0.1626 | -4.8698 | 0.694 | 0.999 |
| stroma → tregs | 9.0654 | 2.2130 | -6.8523 | 0.698 | 0.999 |
| cd4_t_cells_cd45ro → cd68_macrophages | 13.7313 | 3.2175 | -10.5138 | 0.701 | 0.999 |
| cd4_t_cells_cd45ro → b_cells | 4.1235 | 8.6041 | 4.4805 | 0.701 | 0.999 |
| smooth_muscle → granulocytes | 4.2396 | -0.7952 | -5.0348 | 0.716 | 0.999 |
| granulocytes → plasma_cells | 8.4914 | 2.5094 | -5.9819 | 0.717 | 0.999 |
| cd68_macrophages → plasma_cells | 17.5141 | 1.2552 | -16.2588 | 0.724 | 0.999 |
| stroma → smooth_muscle | 5.6620 | 2.5656 | -3.0964 | 0.725 | 0.999 |
| tumor_cells → tumor_cells | 2.4717 | 5.4653 | 2.9936 | 0.728 | 0.999 |
| cd68_cd163_macrophages → cd68_macrophages | 9.6964 | 2.8485 | -6.8479 | 0.729 | 0.999 |
| cd4_t_cells_cd45ro → smooth_muscle | 4.4706 | 0.0745 | -4.3961 | 0.729 | 0.999 |
| tumor_cells → cd8_t_cells | 1.3109 | -2.0916 | -3.4025 | 0.733 | 0.999 |
| cd68_cd163_macrophages → granulocytes | 3.7636 | 1.0677 | -2.6959 | 0.752 | 0.999 |
| vasculature → cd68_cd163_macrophages | 3.1168 | 0.3443 | -2.7725 | 0.766 | 0.999 |
| granulocytes → stroma | 3.9048 | 0.7077 | -3.1971 | 0.770 | 0.999 |
| vasculature → cd8_t_cells | 4.4522 | 1.0022 | -3.4500 | 0.771 | 0.999 |
| plasma_cells → tumor_cells | 3.2045 | -1.0390 | -4.2435 | 0.774 | 0.999 |
| tregs → b_cells | 5.1735 | 11.6221 | 6.4486 | 0.776 | 0.999 |
| granulocytes → cd8_t_cells | 3.5331 | 0.2876 | -3.2455 | 0.776 | 0.999 |
| cd8_t_cells → stroma | 3.9272 | 0.7687 | -3.1585 | 0.778 | 0.999 |
| granulocytes → tregs | 5.7231 | 0.6266 | -5.0965 | 0.781 | 0.999 |
| b_cells → plasma_cells | 0.5706 | 8.9437 | 8.3731 | 0.782 | 0.999 |
| tumor_cells → cd68_macrophages | 7.8693 | 2.3931 | -5.4762 | 0.784 | 0.999 |
| cd68_macrophages → cd68_cd163_macrophages | 7.1938 | 1.6413 | -5.5525 | 0.786 | 0.999 |
| stroma → | 3.0050 | 0.4379 | -2.5671 | 0.786 | 0.999 |
| granulocytes |  |  |  |  |  |
| smooth_muscle → | 4.0816 | 1.1698 | -2.9118 | 0.793 | 0.999 |
| stroma |  |  |  |  |  |
| tumor_cells → | 2.9429 | -0.6576 | -3.6005 | 0.798 | 0.999 |
| b_cells |  |  |  |  |  |
| cd8_t_cells → | 6.6073 | 3.8132 | -2.7940 | 0.801 | 0.999 |
| cd4_t_cells_cd45ro |  |  |  |  |  |
| vasculature → | 1.3609 | -1.8534 | -3.2143 | 0.805 | 0.999 |
| tumor_cells |  |  |  |  |  |
| granulocytes → | 13.3201 | 7.6703 | -5.6498 | 0.806 | 0.999 |
| cd68_macrophages |  |  |  |  |  |
| tumor_cells → | 1.5601 | -0.8209 | -2.3809 | 0.807 | 0.999 |
| cd4_t_cells_cd45ro |  |  |  |  |  |
| smooth_muscle → | -3.5159 | 1.5897 | 5.1056 | 0.822 | 0.999 |
| cd68_macrophages |  |  |  |  |  |
| granulocytes → | 4.0621 | 0.7280 | -3.3341 | 0.828 | 0.999 |
| b_cells |  |  |  |  |  |
| cd68_cd163_macrophages | 3.0134 | 5.2233 | 2.2100 | 0.829 | 0.999 |
| → b_cells |  |  |  |  |  |
| cd68_macrophages | 7.8428 | 2.0642 | -5.7786 | 0.833 | 0.999 |
| → cd4_t_cells_cd45ro |  |  |  |  |  |
| cd4_t_cells_cd45ro → | 4.2365 | 1.8185 | -2.4179 | 0.833 | 0.999 |
| vasculature |  |  |  |  |  |
| b_cells → b_cells | 10.5905 | 6.3983 | -4.1922 | 0.836 | 0.999 |
| b_cells → | 4.6641 | -0.2091 | -4.8732 | 0.837 | 0.999 |
| granulocytes |  |  |  |  |  |
| smooth_muscle → | 4.7296 | -0.3190 | -5.0486 | 0.838 | 0.999 |
| plasma_cells |  |  |  |  |  |
| cd8_t_cells → | 1.5979 | -0.7588 | -2.3567 | 0.843 | 0.999 |
| tumor_cells |  |  |  |  |  |
| stroma → | 1.0535 | 2.5860 | 1.5325 | 0.852 | 0.999 |
| cd4_t_cells_cd45ro |  |  |  |  |  |
| plasma_cells → | 4.2469 | 1.8409 | -2.4061 | 0.856 | 0.999 |
| cd68_cd163_macrophages |  |  |  |  |  |
| b_cells → | -1.6254 | 1.8632 | 3.4886 | 0.856 | 0.999 |
| cd68_cd163_macrophages |  |  |  |  |  |
| plasma_cells → | 4.2460 | 1.1336 | -3.1124 | 0.858 | 0.999 |
| granulocytes |  |  |  |  |  |
| cd68_macrophages | 10.7467 | 6.7849 | -3.9618 | 0.858 | 0.999 |
| → granulocytes |  |  |  |  |  |
| cd68_macrophages | 6.4970 | 2.7474 | -3.7496 | 0.864 | 0.999 |
| → stroma |  |  |  |  |  |
| granulocytes → | 1.5601 | -0.1294 | -1.6895 | 0.864 | 0.999 |
| cd68_cd163_macrophages |  |  |  |  |  |
| cd68_macrophages | 4.7308 | 0.0822 | -4.6486 | 0.865 | 0.999 |
| → cd8_t_cells |  |  |  |  |  |
| smooth_muscle → | -1.0554 | 1.2928 | 2.3483 | 0.869 | 0.999 |
| cd8_t_cells |  |  |  |  |  |
| stroma → | 1.7795 | 4.1235 | 2.3440 | 0.874 | 0.999 |
| plasma_cells |  |  |  |  |  |
| b_cells → vasculature | 0.8027 | 4.1912 | 3.3885 | 0.874 | 0.999 |
| tumor_cells → tregs | 2.0340 | -0.6694 | -2.7035 | 0.874 | 0.999 |
| smooth_muscle → b_cells | -3.8283 | -0.5784 | 3.2499 | 0.876 | 0.999 |
| cd8_t_cells → cd68_cd163_macrophages | 3.6131 | 2.1121 | -1.5010 | 0.878 | 0.999 |
| vasculature → plasma_cells | 6.5907 | 4.1125 | -2.4782 | 0.890 | 0.999 |
| cd68_cd163_macrophages → smooth_muscle | 2.8082 | 1.3949 | -1.4133 | 0.891 | 0.999 |
| stroma → cd68_cd163_macrophages | 1.6576 | 0.7572 | -0.9004 | 0.893 | 0.999 |
| plasma_cells → tregs | 5.3226 | 10.5450 | 5.2225 | 0.893 | 0.999 |
| smooth_muscle → tregs | -3.1744 | 1.2604 | 4.4348 | 0.896 | 0.999 |
| b_cells → cd68_macrophages | 24.2108 | 17.6836 | -6.5272 | 0.897 | 0.999 |
| plasma_cells → cd8_t_cells | 5.0542 | 3.0702 | -1.9840 | 0.897 | 0.999 |
| cd8_t_cells → b_cells | 8.7394 | 6.9260 | -1.8134 | 0.898 | 0.999 |
| cd4_t_cells_cd45ro → cd8_t_cells | 4.1312 | 3.0088 | -1.1224 | 0.902 | 0.999 |
| tregs → plasma_cells | 12.0239 | 7.6338 | -4.3901 | 0.906 | 0.999 |
| stroma → tumor_cells | 2.4921 | 1.5168 | -0.9753 | 0.909 | 0.999 |
| stroma → b_cells | 0.8437 | 2.1954 | 1.3517 | 0.912 | 0.999 |
| plasma_cells → cd4_t_cells_cd45ro | 6.0266 | 4.5041 | -1.5225 | 0.916 | 0.999 |
| cd8_t_cells → plasma_cells | 6.1128 | 4.3909 | -1.7218 | 0.917 | 0.999 |
| cd8_t_cells → granulocytes | 3.1106 | 1.5849 | -1.5257 | 0.918 | 0.999 |
| cd68_macrophages → smooth_muscle | 2.0890 | 5.1769 | 3.0879 | 0.919 | 0.999 |
| vasculature → cd4_t_cells_cd45ro | 3.6903 | 2.5713 | -1.1189 | 0.919 | 0.999 |
| plasma_cells → stroma | 1.3750 | 2.8778 | 1.5028 | 0.919 | 0.999 |
| vasculature → tregs | -0.0623 | 2.8527 | 2.9150 | 0.921 | 0.999 |
| tregs → tregs | 11.1620 | 7.6627 | -3.4993 | 0.925 | 0.999 |
| b_cells → tumor_cells | 2.8742 | 0.9338 | -1.9403 | 0.927 | 0.999 |
| b_cells → smooth_muscle | -2.5461 | 0.3992 | 2.9453 | 0.930 | 0.999 |
| cd8_t_cells → tregs | 6.9962 | 5.2921 | -1.7040 | 0.932 | 0.999 |
| cd68_cd163_macrophages → tregs | 2.7315 | 4.4656 | 1.7341 | 0.932 | 0.999 |
| stroma → stroma | -0.1426 | 0.4478 | 0.5903 | 0.935 | 0.999 |
| vasculature → b_cells | 1.5698 | 2.8203 | 1.2505 | 0.939 | 0.999 |
| tregs → | 4.4954 | 3.2653 | -1.2301 | 0.940 | 0.999 |
| cd68_cd163_macrophages |  |  |  |  |  |
| cd4_t.cells_cd45ro → | 1.9091 | 1.2753 | -0.6338 | 0.942 | 0.999 |
| cd4_t.cells_cd45ro |  |  |  |  |  |
| smooth_muscle → | 0.3756 | -0.6882 | -1.0638 | 0.942 | 0.999 |
| tumor_cells |  |  |  |  |  |
| b_cells → cd8_t.cells | 7.1118 | 5.8068 | -1.3050 | 0.945 | 0.999 |
| cd4_t.cells_cd45ro → | 3.2353 | 4.3175 | 1.0822 | 0.946 | 0.999 |
| tregs |  |  |  |  |  |
| cd68_macrophages | 2.8725 | 1.3930 | -1.4796 | 0.949 | 0.999 |
| → tumor_cells |  |  |  |  |  |
| tregs → | 3.3038 | 1.8928 | -1.4110 | 0.951 | 0.999 |
| smooth_muscle |  |  |  |  |  |
| tumor_cells → | 0.5013 | -0.1050 | -0.6063 | 0.954 | 0.999 |
| granulocytes |  |  |  |  |  |
| plasma_cells → | 7.8005 | 8.9311 | 1.1306 | 0.954 | 0.999 |
| b_cells |  |  |  |  |  |
| cd8_t.cells → | 0.6374 | 1.5413 | 0.9039 | 0.955 | 0.999 |
| smooth_muscle |  |  |  |  |  |
| smooth_muscle → | 1.0649 | 0.3778 | -0.6871 | 0.956 | 0.999 |
| cd4_t.cells_cd45ro |  |  |  |  |  |
| granulocytes → | 0.6889 | 0.0128 | -0.6761 | 0.956 | 0.999 |
| tumor_cells |  |  |  |  |  |
| cd4_t.cells_cd45ro → | 1.0122 | 1.4760 | 0.4638 | 0.958 | 0.999 |
| cd68_cd163_macrophages |  |  |  |  |  |
| tregs → | 3.0549 | 4.7286 | 1.6737 | 0.958 | 0.999 |
| cd68_macrophages |  |  |  |  |  |
| tregs → cd8_t.cells | 5.4228 | 4.4752 | -0.9476 | 0.959 | 0.999 |
| cd68_macrophages | 7.3006 | 9.3806 | 2.0800 | 0.959 | 0.999 |
| → b_cells |  |  |  |  |  |
| b_cells → tregs | 10.7686 | 9.1757 | -1.5929 | 0.960 | 0.999 |
| cd68_cd163_macrophages | 0.8192 | 0.4637 | -0.3555 | 0.962 | 0.999 |
| → stroma |  |  |  |  |  |
| stroma → | 3.7232 | 2.9317 | -0.7916 | 0.963 | 0.999 |
| cd68_macrophages |  |  |  |  |  |
| smooth_muscle → | 0.2394 | 0.7406 | 0.5012 | 0.964 | 0.999 |
| cd68_cd163_macrophages |  |  |  |  |  |
| cd68_cd163_macrophages | 2.5839 | 2.9419 | 0.3580 | 0.964 | 0.999 |
| → cd8_t.cells |  |  |  |  |  |
| tregs → | 2.2992 | 1.3029 | -0.9963 | 0.964 | 0.999 |
| granulocytes |  |  |  |  |  |
| cd4_t.cells_cd45ro → | 0.1942 | -0.2614 | -0.4556 | 0.965 | 0.999 |
| tumor_cells |  |  |  |  |  |
| b_cells → stroma | 1.4748 | 0.6694 | -0.8054 | 0.967 | 0.999 |
| stroma → | 2.9399 | 2.5780 | -0.3618 | 0.969 | 0.999 |
| vasculature |  |  |  |  |  |
| tregs → | 4.9632 | 4.2671 | -0.6961 | 0.970 | 0.999 |
| cd4_t.cells_cd45ro |  |  |  |  |  |
| vasculature → | -0.1546 | 0.8791 | 1.0336 | 0.971 | 0.999 |
| cd68_macrophages |  |  |  |  |  |
| tregs → vasculature | 1.9642 | 2.8611 | 0.8970 | 0.973 | 0.999 |
| cd4_t_cells_cd45ro → granulocytes | 0.8999 | 1.2571 | 0.3571 | 0.974 | 0.999 |
| b_cells → cd4_t_cells_cd45ro | 6.5331 | 7.0917 | 0.5586 | 0.975 | 0.999 |
| cd68_cd163_macrophages → cd4_t_cells_cd45ro | 1.7435 | 1.9904 | 0.2469 | 0.976 | 0.999 |
| cd68_cd163_macrophages → cd68_cd163_macrophages | 1.1211 | 1.3064 | 0.1853 | 0.977 | 0.999 |
| cd4_t_cells_cd45ro → plasma_cells | 5.0204 | 5.3836 | 0.3632 | 0.982 | 0.999 |
| cd4_t_cells_cd45ro → stroma | 2.5637 | 2.3829 | -0.1808 | 0.986 | 0.999 |
| vasculature → stroma | 2.8007 | 2.6361 | -0.1646 | 0.988 | 0.999 |
| cd68_macrophages → tregs | 1.6504 | 1.1485 | -0.5019 | 0.992 | 0.999 |
| cd68_macrophages → vasculature | 3.2834 | 3.1395 | -0.1438 | 0.997 | 0.999 |
| tregs → tumor_cells | 0.8129 | 0.7451 | -0.0678 | 0.997 | 0.999 |
| stroma → cd8_t_cells | 1.2013 | 1.2167 | 0.0154 | 0.999 | 0.999 |

**Figure S6:**
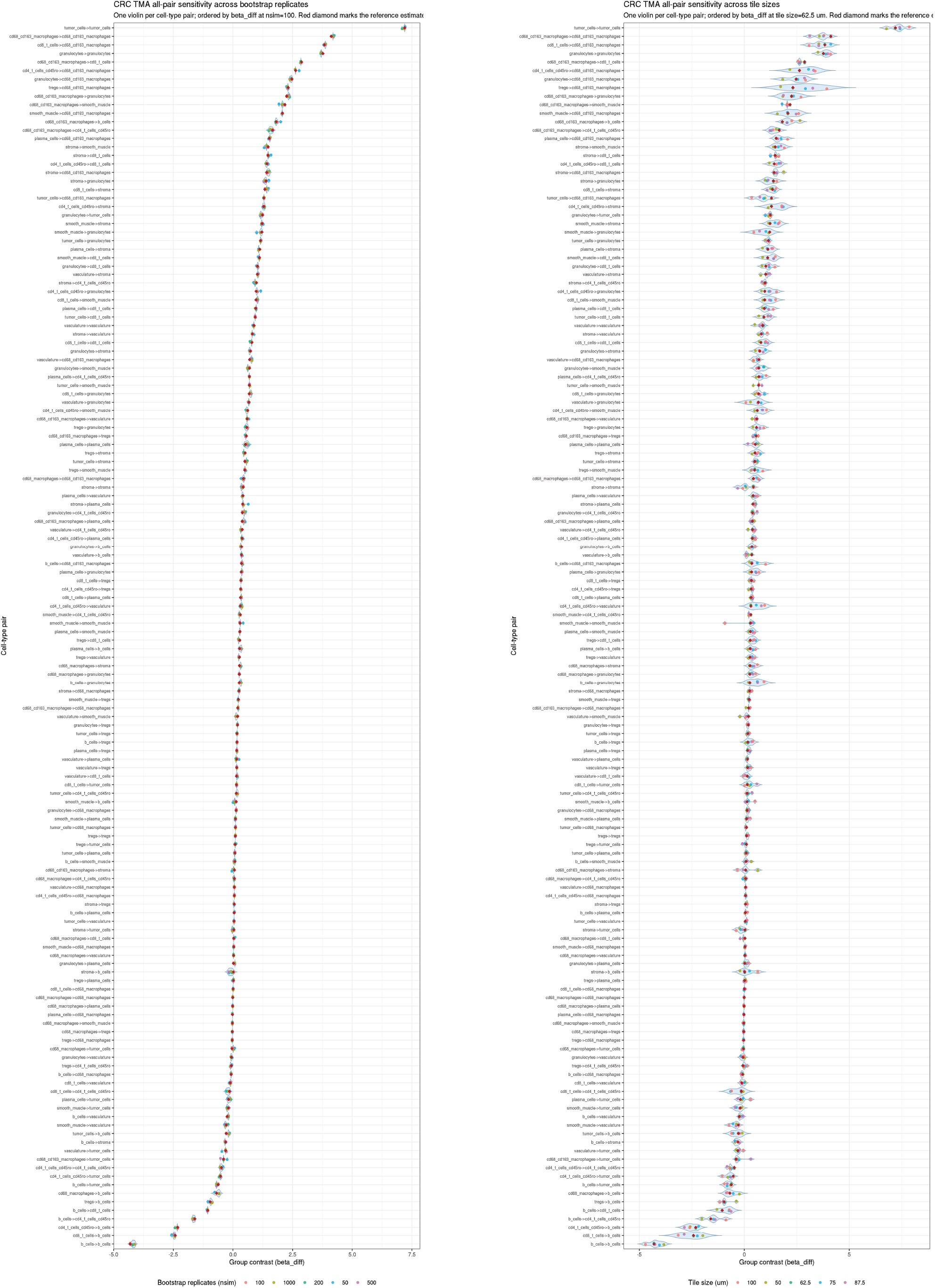
Left: Sensitivity analysis for the number of bootstrap iterations. Right: Sensitivity analysis for the choice of tile size in the spatial bootstrap.

#### S8.10 Sensitivity analysis

### S9 HNSCC supplementary results

#### S9.1 HNSCC cohort overview

**Figure S7:**
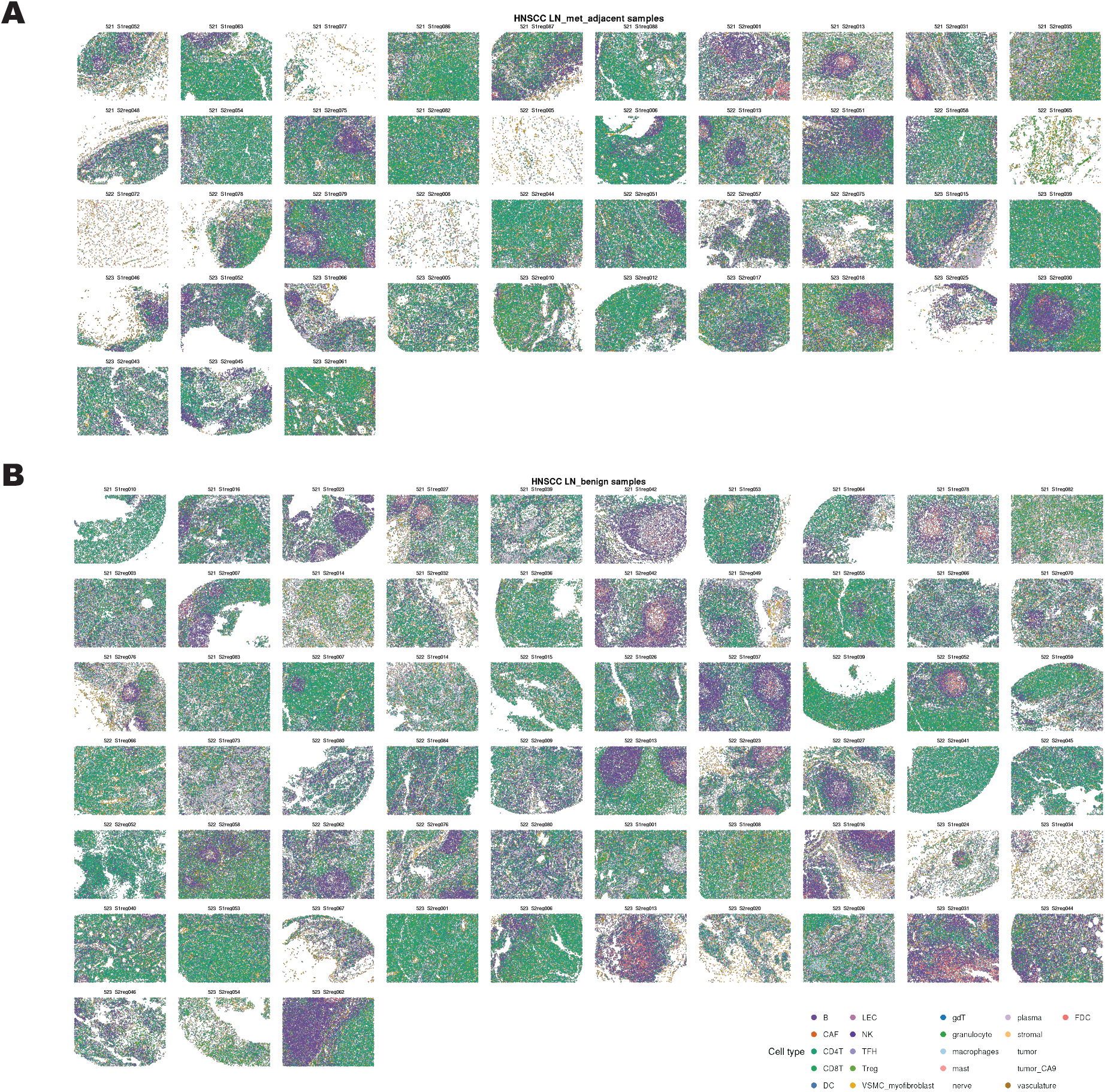
Overview of HNSCC lymph node TMA samples used in the PANORAMIC analysis. Each panel shows spatial cell maps from the Haist et al. HNSCC cohort, with points representing single cells colored by annotated cell type. (A) Tumor-adjacent lymph node regions from patients with lymph node metastasis. (B) Benign lymph node samples from node-negative patients.

**Figure S8:**
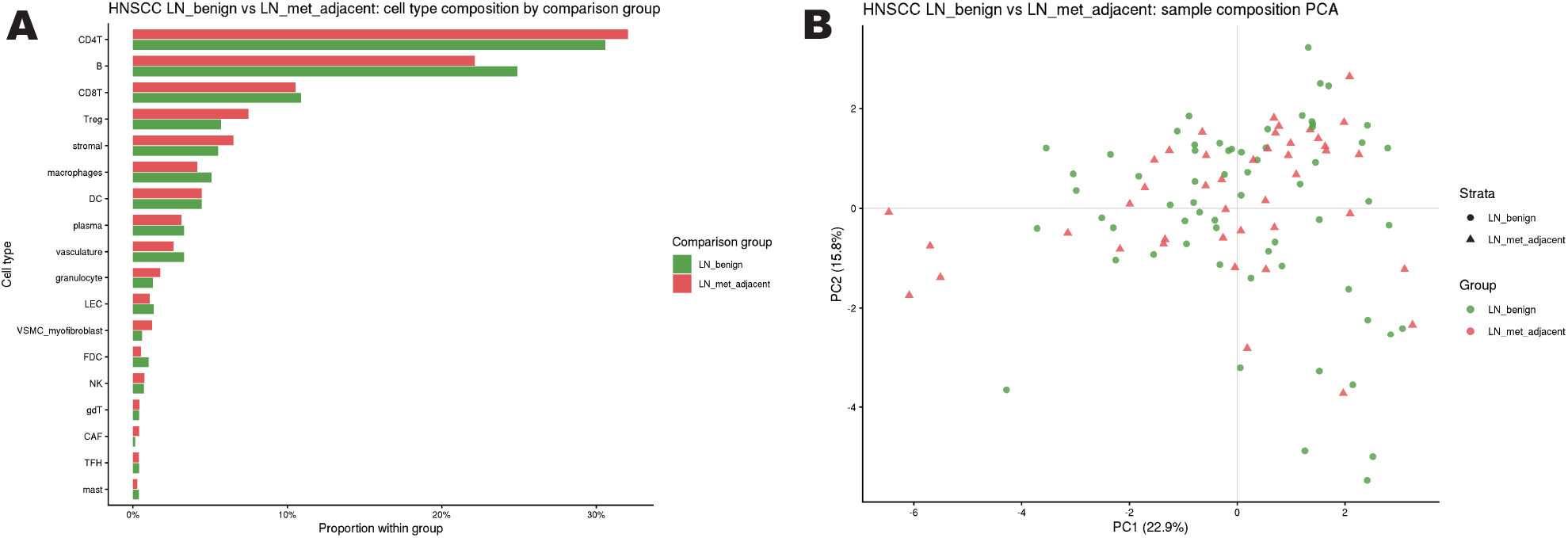
Caption

#### S9.2 Full PANORAMIC HNSCC results at 25 *µ*m

**Table S6:**
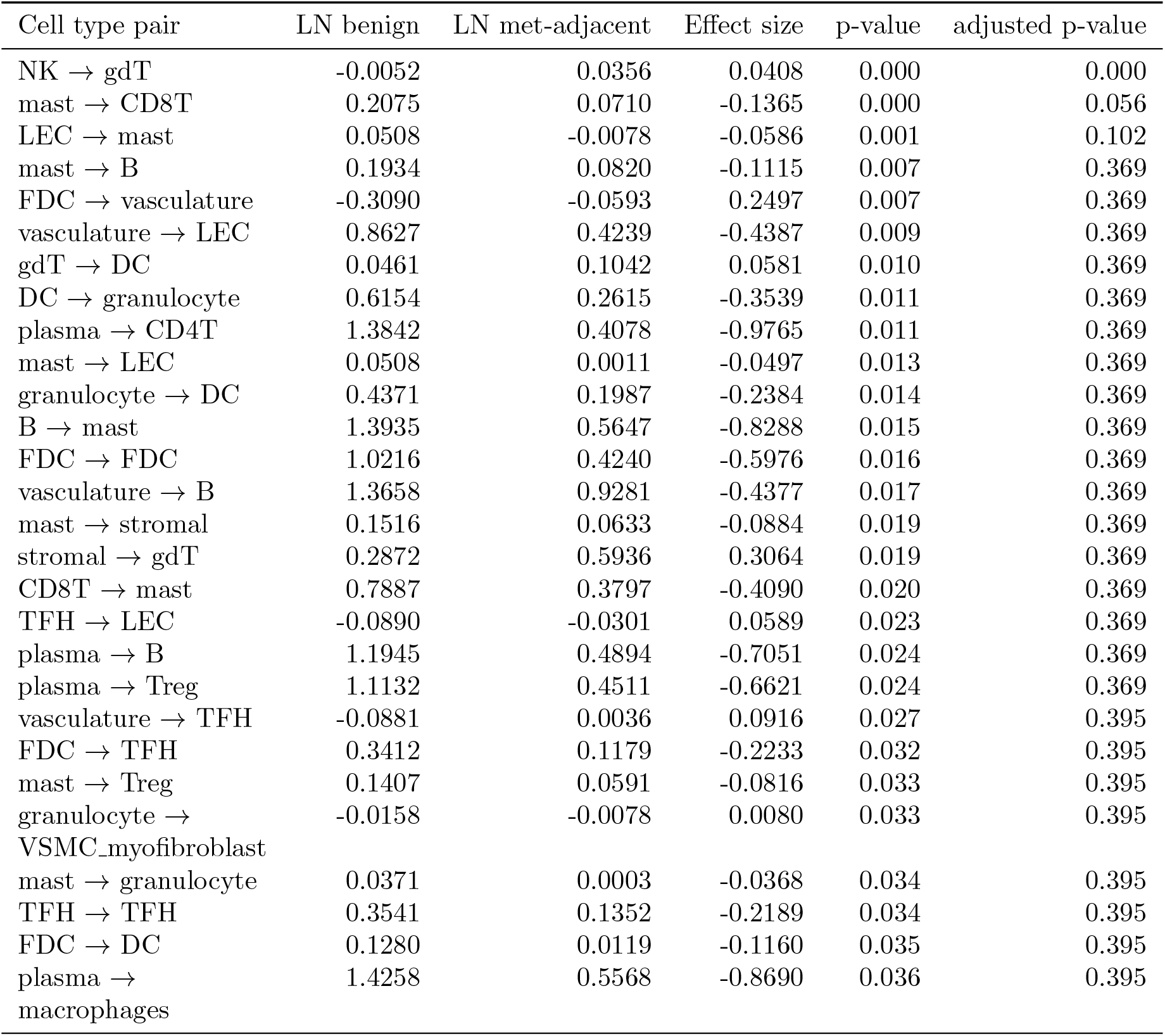
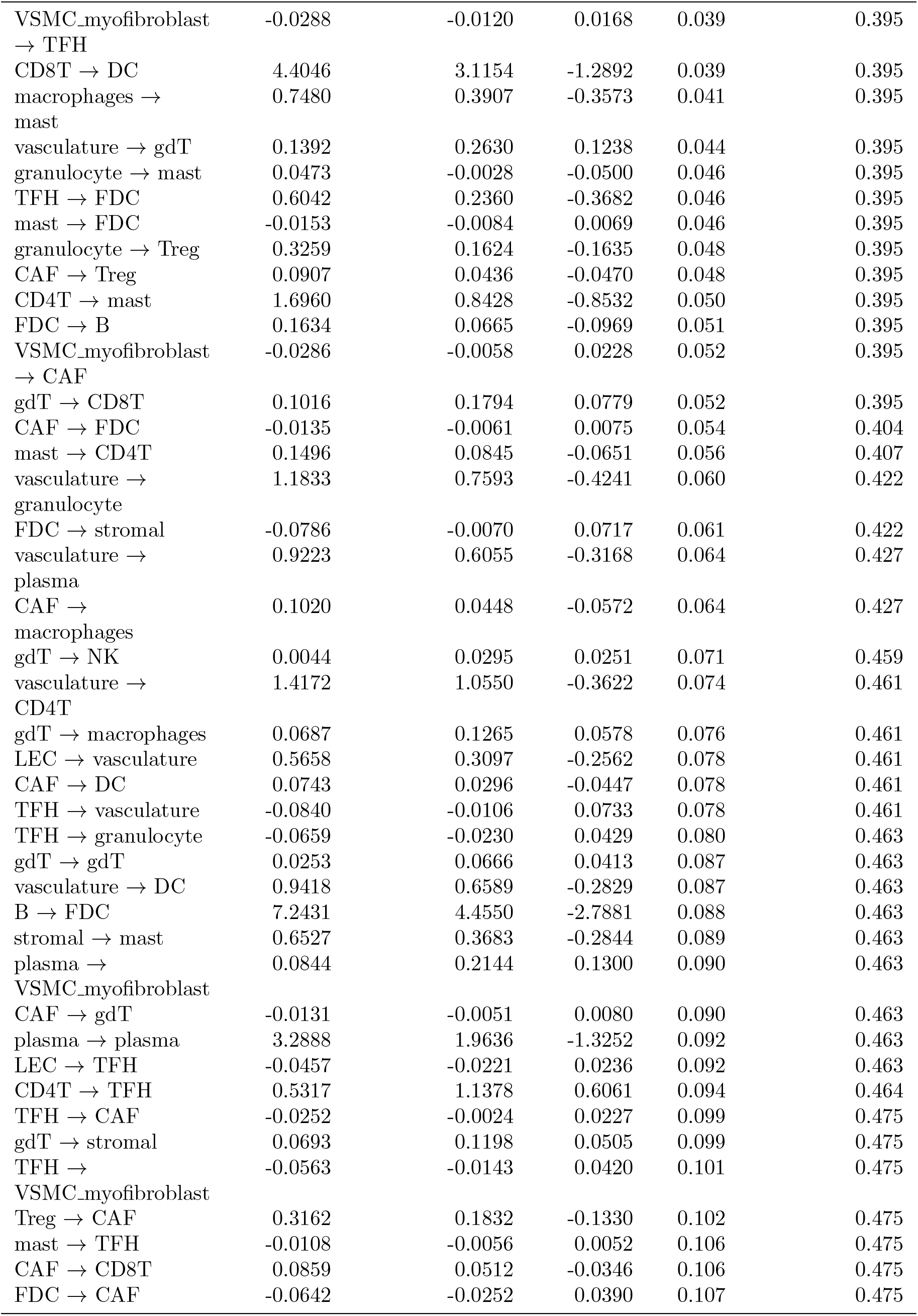
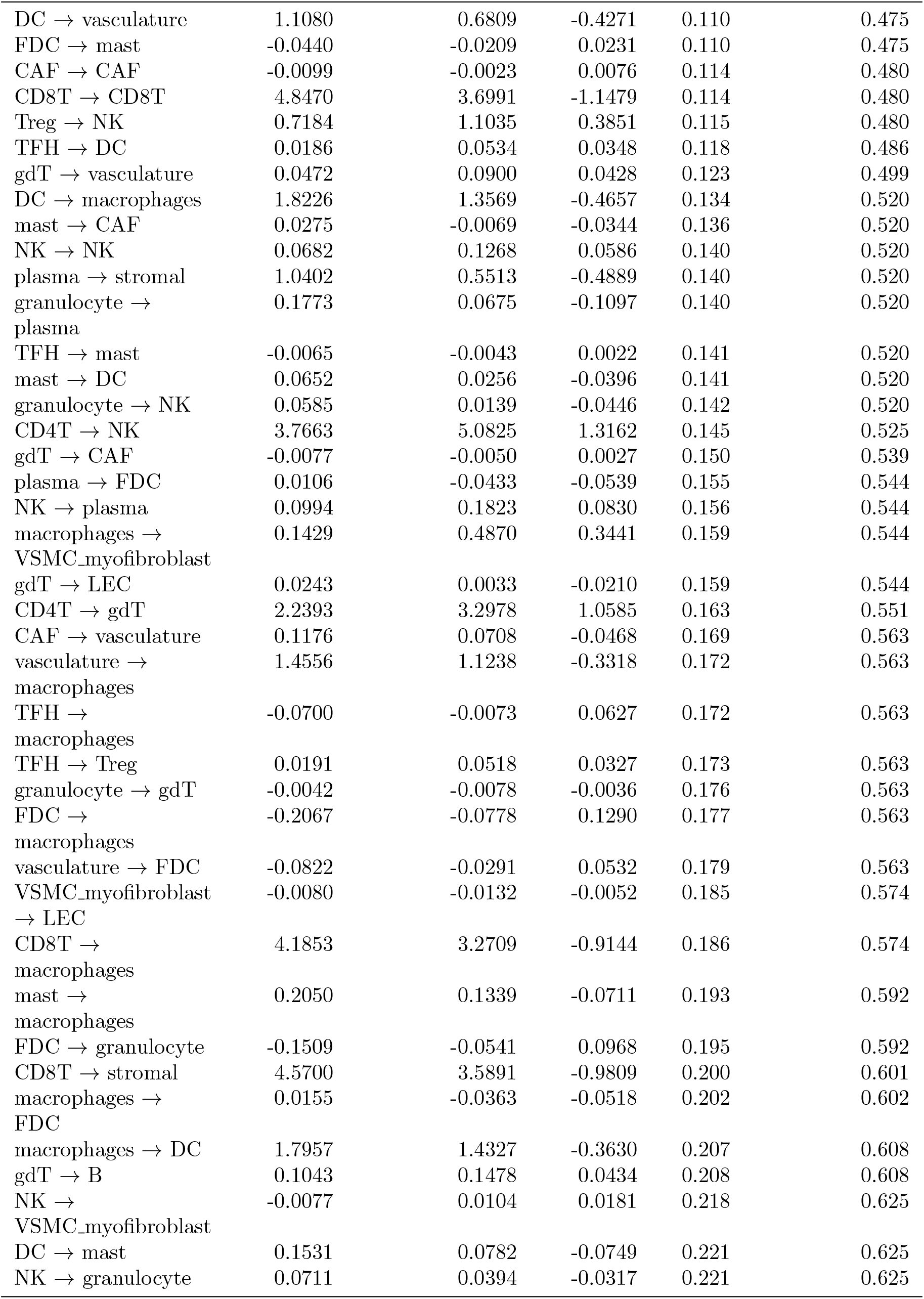
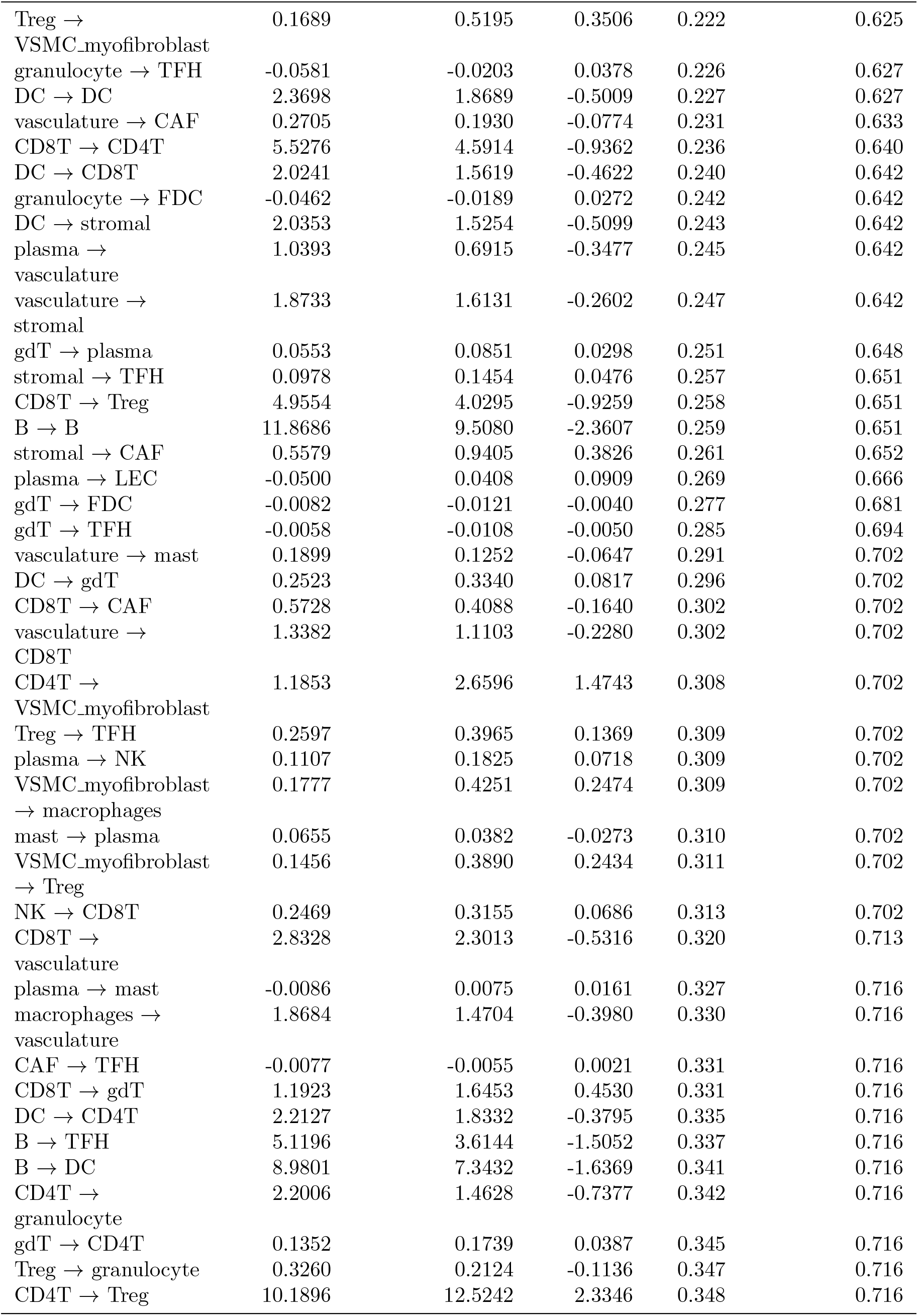
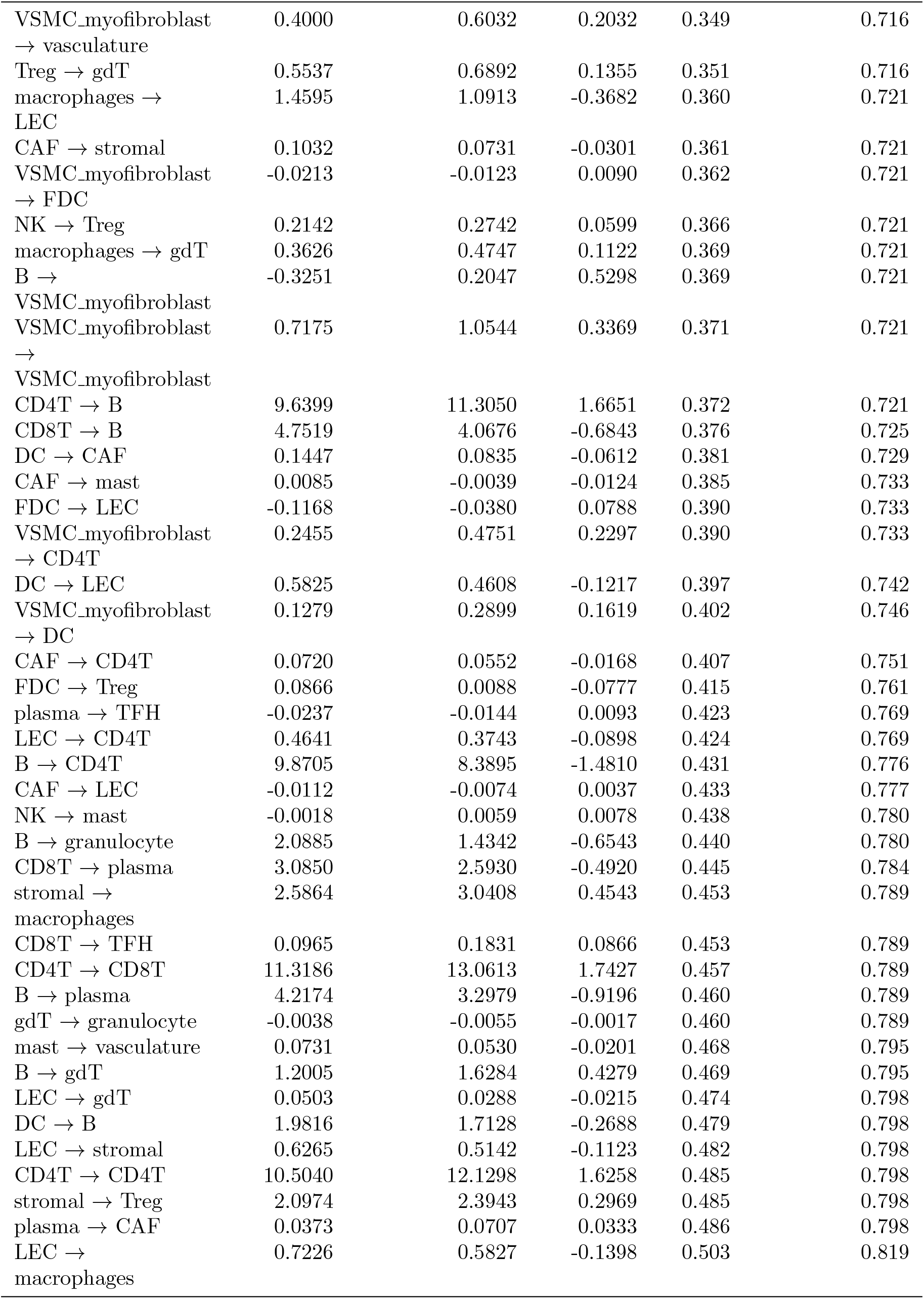
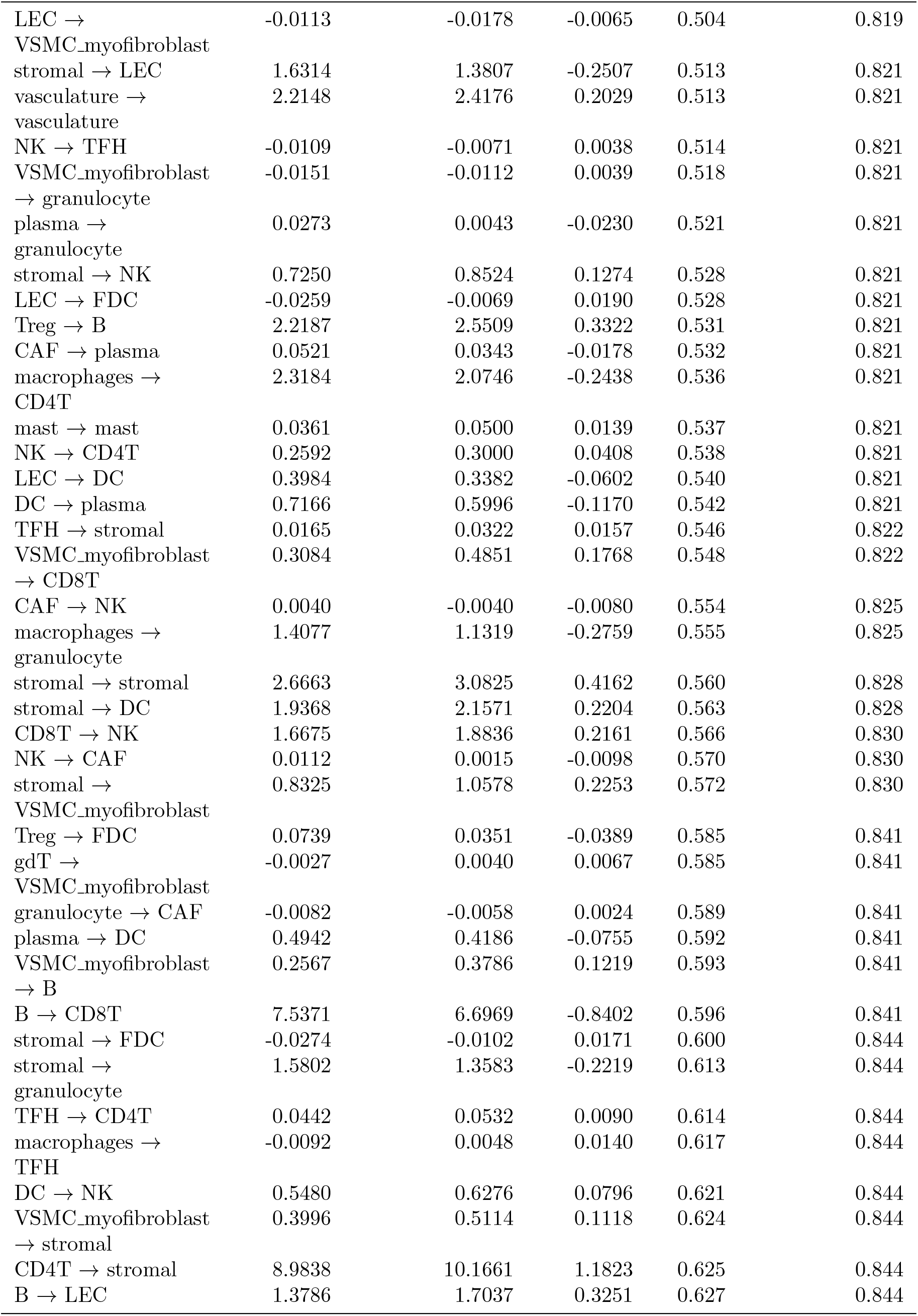
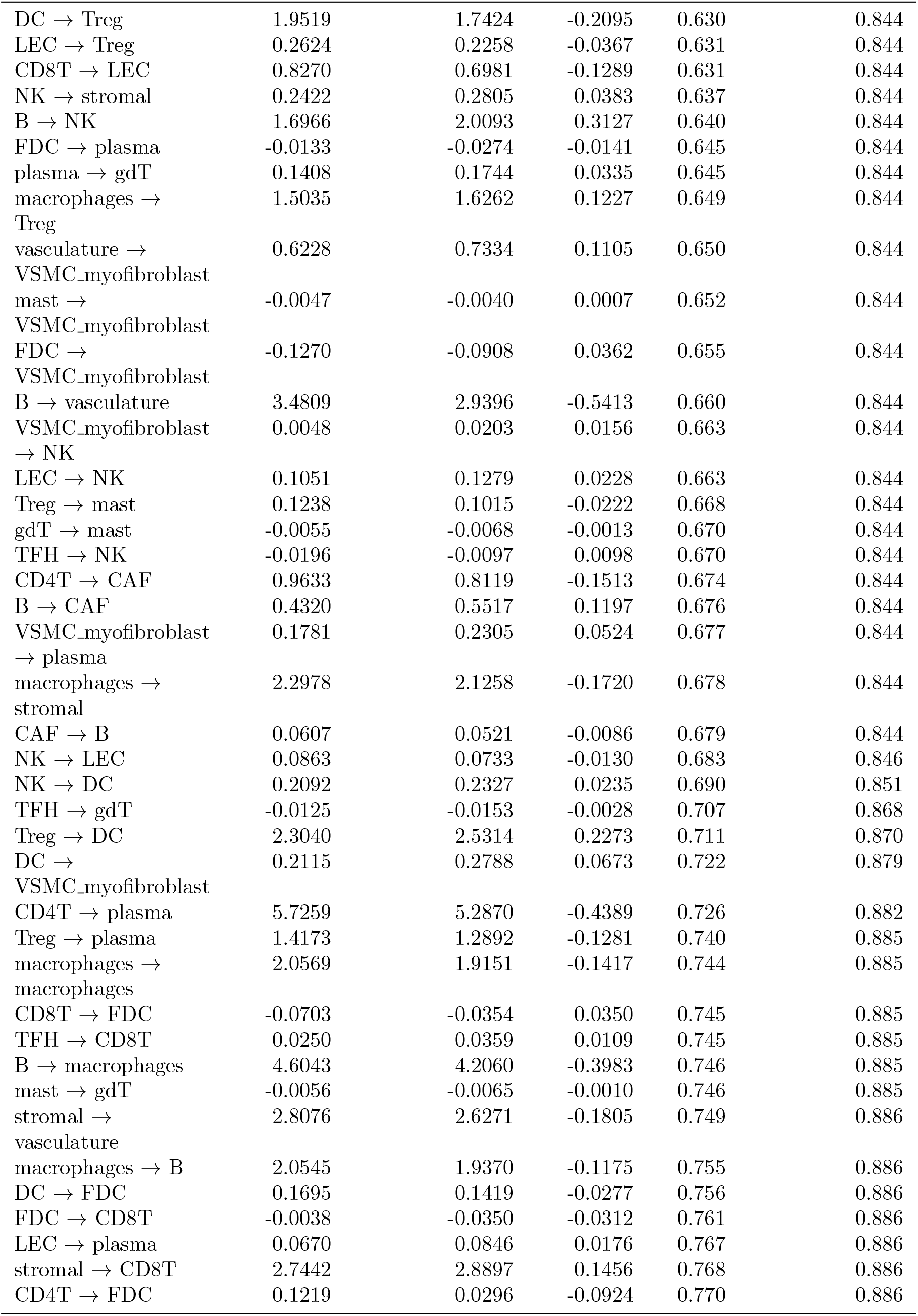
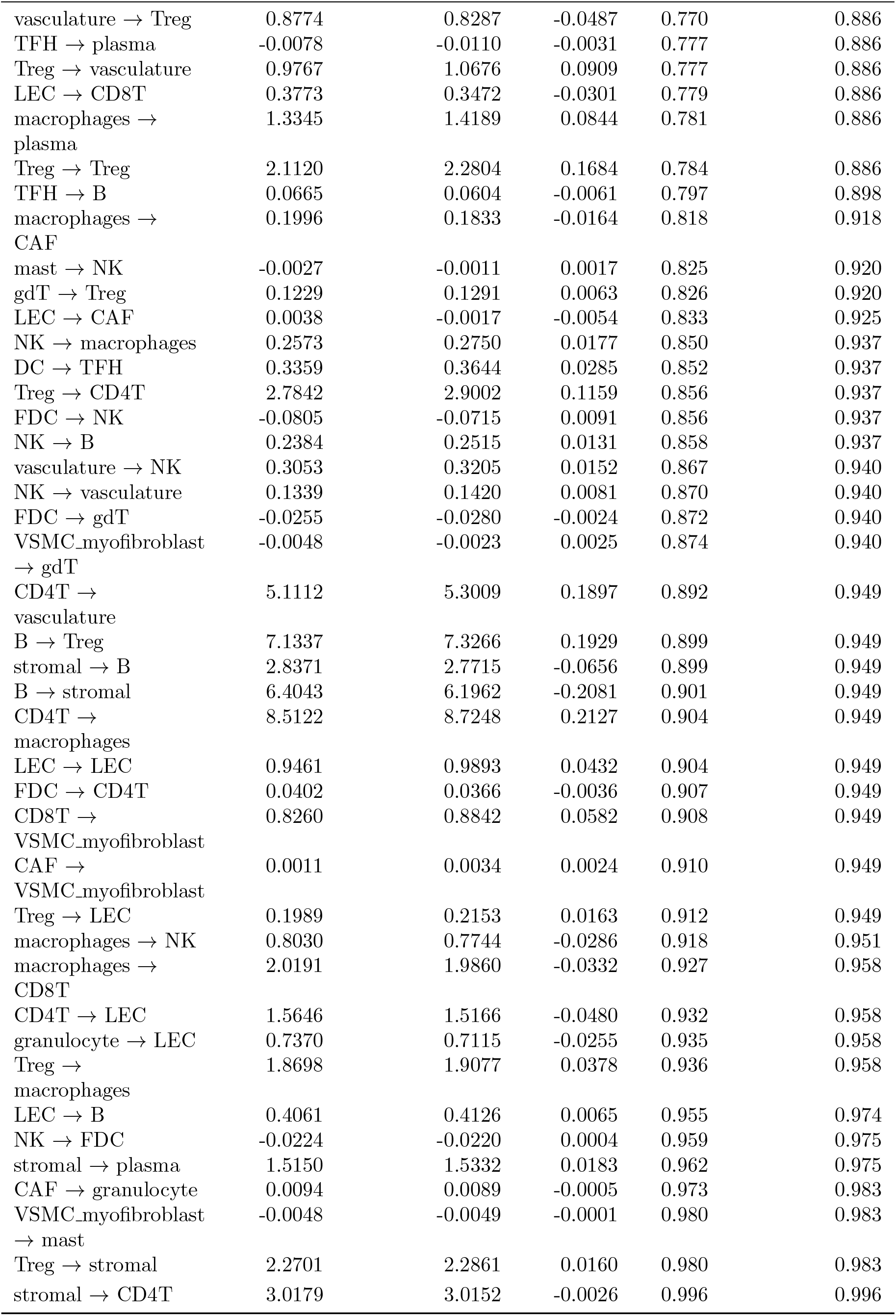
HNSCC TMA PANORAMIC results, sorted by p-value.

| Cell type pair | LN benign | LN met-adjacent | Effect size | p-value | adjusted p-value |
| --- | --- | --- | --- | --- | --- |
| NK → gdT | -0.0052 | 0.0356 | 0.0408 | 0.000 | 0.000 |
| mast → CD8T | 0.2075 | 0.0710 | -0.1365 | 0.000 | 0.056 |
| LEC → mast | 0.0508 | -0.0078 | -0.0586 | 0.001 | 0.102 |
| mast → B | 0.1934 | 0.0820 | -0.1115 | 0.007 | 0.369 |
| FDC → vasculature | -0.3090 | -0.0593 | 0.2497 | 0.007 | 0.369 |
| vasculature → LEC | 0.8627 | 0.4239 | -0.4387 | 0.009 | 0.369 |
| gdT → DC | 0.0461 | 0.1042 | 0.0581 | 0.010 | 0.369 |
| DC → granulocyte | 0.6154 | 0.2615 | -0.3539 | 0.011 | 0.369 |
| plasma → CD4T | 1.3842 | 0.4078 | -0.9765 | 0.011 | 0.369 |
| mast → LEC | 0.0508 | 0.0011 | -0.0497 | 0.013 | 0.369 |
| granulocyte → DC | 0.4371 | 0.1987 | -0.2384 | 0.014 | 0.369 |
| B → mast | 1.3935 | 0.5647 | -0.8288 | 0.015 | 0.369 |
| FDC → FDC | 1.0216 | 0.4240 | -0.5976 | 0.016 | 0.369 |
| vasculature → B | 1.3658 | 0.9281 | -0.4377 | 0.017 | 0.369 |
| mast → stromal | 0.1516 | 0.0633 | -0.0884 | 0.019 | 0.369 |
| stromal → gdT | 0.2872 | 0.5936 | 0.3064 | 0.019 | 0.369 |
| CD8T → mast | 0.7887 | 0.3797 | -0.4090 | 0.020 | 0.369 |
| TFH → LEC | -0.0890 | -0.0301 | 0.0589 | 0.023 | 0.369 |
| plasma → B | 1.1945 | 0.4894 | -0.7051 | 0.024 | 0.369 |
| plasma → Treg | 1.1132 | 0.4511 | -0.6621 | 0.024 | 0.369 |
| vasculature → TFH | -0.0881 | 0.0036 | 0.0916 | 0.027 | 0.395 |
| FDC → TFH | 0.3412 | 0.1179 | -0.2233 | 0.032 | 0.395 |
| mast → Treg | 0.1407 | 0.0591 | -0.0816 | 0.033 | 0.395 |
| granulocyte → VSMC_myofibroblast | -0.0158 | -0.0078 | 0.0080 | 0.033 | 0.395 |
| mast → granulocyte | 0.0371 | 0.0003 | -0.0368 | 0.034 | 0.395 |
| TFH → TFH | 0.3541 | 0.1352 | -0.2189 | 0.034 | 0.395 |
| FDC → DC | 0.1280 | 0.0119 | -0.1160 | 0.035 | 0.395 |
| plasma → macrophages | 1.4258 | 0.5568 | -0.8690 | 0.036 | 0.395 |
| VSMC_myofibroblast | -0.0288 | -0.0120 | 0.0168 | 0.039 | 0.395 |
| → TFH |  |  |  |  |  |
| CD8T → DC | 4.4046 | 3.1154 | -1.2892 | 0.039 | 0.395 |
| macrophages → | 0.7480 | 0.3907 | -0.3573 | 0.041 | 0.395 |
| mast |  |  |  |  |  |
| vasculature → gdT | 0.1392 | 0.2630 | 0.1238 | 0.044 | 0.395 |
| granulocyte → mast | 0.0473 | -0.0028 | -0.0500 | 0.046 | 0.395 |
| TFH → FDC | 0.6042 | 0.2360 | -0.3682 | 0.046 | 0.395 |
| mast → FDC | -0.0153 | -0.0084 | 0.0069 | 0.046 | 0.395 |
| granulocyte → Treg | 0.3259 | 0.1624 | -0.1635 | 0.048 | 0.395 |
| CAF → Treg | 0.0907 | 0.0436 | -0.0470 | 0.048 | 0.395 |
| CD4T → mast | 1.6960 | 0.8428 | -0.8532 | 0.050 | 0.395 |
| FDC → B | 0.1634 | 0.0665 | -0.0969 | 0.051 | 0.395 |
| VSMC_myofibroblast | -0.0286 | -0.0058 | 0.0228 | 0.052 | 0.395 |
| → CAF |  |  |  |  |  |
| gdT → CD8T | 0.1016 | 0.1794 | 0.0779 | 0.052 | 0.395 |
| CAF → FDC | -0.0135 | -0.0061 | 0.0075 | 0.054 | 0.404 |
| mast → CD4T | 0.1496 | 0.0845 | -0.0651 | 0.056 | 0.407 |
| vasculature → | 1.1833 | 0.7593 | -0.4241 | 0.060 | 0.422 |
| granulocyte |  |  |  |  |  |
| FDC → stromal | -0.0786 | -0.0070 | 0.0717 | 0.061 | 0.422 |
| vasculature → | 0.9223 | 0.6055 | -0.3168 | 0.064 | 0.427 |
| plasma |  |  |  |  |  |
| CAF → | 0.1020 | 0.0448 | -0.0572 | 0.064 | 0.427 |
| macrophages |  |  |  |  |  |
| gdT → NK | 0.0044 | 0.0295 | 0.0251 | 0.071 | 0.459 |
| vasculature → | 1.4172 | 1.0550 | -0.3622 | 0.074 | 0.461 |
| CD4T |  |  |  |  |  |
| gdT → macrophages | 0.0687 | 0.1265 | 0.0578 | 0.076 | 0.461 |
| LEC → vasculature | 0.5658 | 0.3097 | -0.2562 | 0.078 | 0.461 |
| CAF → DC | 0.0743 | 0.0296 | -0.0447 | 0.078 | 0.461 |
| TFH → vasculature | -0.0840 | -0.0106 | 0.0733 | 0.078 | 0.461 |
| TFH → granulocyte | -0.0659 | -0.0230 | 0.0429 | 0.080 | 0.463 |
| gdT → gdT | 0.0253 | 0.0666 | 0.0413 | 0.087 | 0.463 |
| vasculature → DC | 0.9418 | 0.6589 | -0.2829 | 0.087 | 0.463 |
| B → FDC | 7.2431 | 4.4550 | -2.7881 | 0.088 | 0.463 |
| stromal → mast | 0.6527 | 0.3683 | -0.2844 | 0.089 | 0.463 |
| plasma → | 0.0844 | 0.2144 | 0.1300 | 0.090 | 0.463 |
| VSMC_myofibroblast |  |  |  |  |  |
| CAF → gdT | -0.0131 | -0.0051 | 0.0080 | 0.090 | 0.463 |
| plasma → plasma | 3.2888 | 1.9636 | -1.3252 | 0.092 | 0.463 |
| LEC → TFH | -0.0457 | -0.0221 | 0.0236 | 0.092 | 0.463 |
| CD4T → TFH | 0.5317 | 1.1378 | 0.6061 | 0.094 | 0.464 |
| TFH → CAF | -0.0252 | -0.0024 | 0.0227 | 0.099 | 0.475 |
| gdT → stromal | 0.0693 | 0.1198 | 0.0505 | 0.099 | 0.475 |
| TFH → | -0.0563 | -0.0143 | 0.0420 | 0.101 | 0.475 |
| VSMC_myofibroblast |  |  |  |  |  |
| Treg → CAF | 0.3162 | 0.1832 | -0.1330 | 0.102 | 0.475 |
| mast → TFH | -0.0108 | -0.0056 | 0.0052 | 0.106 | 0.475 |
| CAF → CD8T | 0.0859 | 0.0512 | -0.0346 | 0.106 | 0.475 |
| FDC → CAF | -0.0642 | -0.0252 | 0.0390 | 0.107 | 0.475 |
| DC → vasculature | 1.1080 | 0.6809 | -0.4271 | 0.110 | 0.475 |
| FDC → mast | -0.0440 | -0.0209 | 0.0231 | 0.110 | 0.475 |
| CAF → CAF | -0.0099 | -0.0023 | 0.0076 | 0.114 | 0.480 |
| CD8T → CD8T | 4.8470 | 3.6991 | -1.1479 | 0.114 | 0.480 |
| Treg → NK | 0.7184 | 1.1035 | 0.3851 | 0.115 | 0.480 |
| TFH → DC | 0.0186 | 0.0534 | 0.0348 | 0.118 | 0.486 |
| gdT → vasculature | 0.0472 | 0.0900 | 0.0428 | 0.123 | 0.499 |
| DC → macrophages | 1.8226 | 1.3569 | -0.4657 | 0.134 | 0.520 |
| mast → CAF | 0.0275 | -0.0069 | -0.0344 | 0.136 | 0.520 |
| NK → NK | 0.0682 | 0.1268 | 0.0586 | 0.140 | 0.520 |
| plasma → stromal | 1.0402 | 0.5513 | -0.4889 | 0.140 | 0.520 |
| granulocyte → plasma | 0.1773 | 0.0675 | -0.1097 | 0.140 | 0.520 |
| TFH → mast | -0.0065 | -0.0043 | 0.0022 | 0.141 | 0.520 |
| mast → DC | 0.0652 | 0.0256 | -0.0396 | 0.141 | 0.520 |
| granulocyte → NK | 0.0585 | 0.0139 | -0.0446 | 0.142 | 0.520 |
| CD4T → NK | 3.7663 | 5.0825 | 1.3162 | 0.145 | 0.525 |
| gdT → CAF | -0.0077 | -0.0050 | 0.0027 | 0.150 | 0.539 |
| plasma → FDC | 0.0106 | -0.0433 | -0.0539 | 0.155 | 0.544 |
| NK → plasma | 0.0994 | 0.1823 | 0.0830 | 0.156 | 0.544 |
| macrophages → VSMC_myofibroblast | 0.1429 | 0.4870 | 0.3441 | 0.159 | 0.544 |
| gdT → LEC | 0.0243 | 0.0033 | -0.0210 | 0.159 | 0.544 |
| CD4T → gdT | 2.2393 | 3.2978 | 1.0585 | 0.163 | 0.551 |
| CAF → vasculature | 0.1176 | 0.0708 | -0.0468 | 0.169 | 0.563 |
| vasculature → macrophages | 1.4556 | 1.1238 | -0.3318 | 0.172 | 0.563 |
| TFH → macrophages | -0.0700 | -0.0073 | 0.0627 | 0.172 | 0.563 |
| TFH → Treg | 0.0191 | 0.0518 | 0.0327 | 0.173 | 0.563 |
| granulocyte → gdT | -0.0042 | -0.0078 | -0.0036 | 0.176 | 0.563 |
| FDC → macrophages | -0.2067 | -0.0778 | 0.1290 | 0.177 | 0.563 |
| vasculature → FDC | -0.0822 | -0.0291 | 0.0532 | 0.179 | 0.563 |
| VSMC_myofibroblast → LEC | -0.0080 | -0.0132 | -0.0052 | 0.185 | 0.574 |
| CD8T → macrophages | 4.1853 | 3.2709 | -0.9144 | 0.186 | 0.574 |
| mast → macrophages | 0.2050 | 0.1339 | -0.0711 | 0.193 | 0.592 |
| FDC → granulocyte | -0.1509 | -0.0541 | 0.0968 | 0.195 | 0.592 |
| CD8T → stromal | 4.5700 | 3.5891 | -0.9809 | 0.200 | 0.601 |
| macrophages → FDC | 0.0155 | -0.0363 | -0.0518 | 0.202 | 0.602 |
| macrophages → DC | 1.7957 | 1.4327 | -0.3630 | 0.207 | 0.608 |
| gdT → B | 0.1043 | 0.1478 | 0.0434 | 0.208 | 0.608 |
| NK → VSMC_myofibroblast | -0.0077 | 0.0104 | 0.0181 | 0.218 | 0.625 |
| DC → mast | 0.1531 | 0.0782 | -0.0749 | 0.221 | 0.625 |
| NK → granulocyte | 0.0711 | 0.0394 | -0.0317 | 0.221 | 0.625 |
| Treg → | 0.1689 | 0.5195 | 0.3506 | 0.222 | 0.625 |
| VSMC_myofibroblast |  |  |  |  |  |
| granulocyte → TFH | -0.0581 | -0.0203 | 0.0378 | 0.226 | 0.627 |
| DC → DC | 2.3698 | 1.8689 | -0.5009 | 0.227 | 0.627 |
| vasculature → CAF | 0.2705 | 0.1930 | -0.0774 | 0.231 | 0.633 |
| CD8T → CD4T | 5.5276 | 4.5914 | -0.9362 | 0.236 | 0.640 |
| DC → CD8T | 2.0241 | 1.5619 | -0.4622 | 0.240 | 0.642 |
| granulocyte → FDC | -0.0462 | -0.0189 | 0.0272 | 0.242 | 0.642 |
| DC → stromal | 2.0353 | 1.5254 | -0.5099 | 0.243 | 0.642 |
| plasma → | 1.0393 | 0.6915 | -0.3477 | 0.245 | 0.642 |
| vasculature |  |  |  |  |  |
| vasculature → | 1.8733 | 1.6131 | -0.2602 | 0.247 | 0.642 |
| stromal |  |  |  |  |  |
| gdT → plasma | 0.0553 | 0.0851 | 0.0298 | 0.251 | 0.648 |
| stromal → TFH | 0.0978 | 0.1454 | 0.0476 | 0.257 | 0.651 |
| CD8T → Treg | 4.9554 | 4.0295 | -0.9259 | 0.258 | 0.651 |
| B → B | 11.8686 | 9.5080 | -2.3607 | 0.259 | 0.651 |
| stromal → CAF | 0.5579 | 0.9405 | 0.3826 | 0.261 | 0.652 |
| plasma → LEC | -0.0500 | 0.0408 | 0.0909 | 0.269 | 0.666 |
| gdT → FDC | -0.0082 | -0.0121 | -0.0040 | 0.277 | 0.681 |
| gdT → TFH | -0.0058 | -0.0108 | -0.0050 | 0.285 | 0.694 |
| vasculature → mast | 0.1899 | 0.1252 | -0.0647 | 0.291 | 0.702 |
| DC → gdT | 0.2523 | 0.3340 | 0.0817 | 0.296 | 0.702 |
| CD8T → CAF | 0.5728 | 0.4088 | -0.1640 | 0.302 | 0.702 |
| vasculature → | 1.3382 | 1.1103 | -0.2280 | 0.302 | 0.702 |
| CD8T |  |  |  |  |  |
| CD4T → | 1.1853 | 2.6596 | 1.4743 | 0.308 | 0.702 |
| VSMC_myofibroblast |  |  |  |  |  |
| Treg → TFH | 0.2597 | 0.3965 | 0.1369 | 0.309 | 0.702 |
| plasma → NK | 0.1107 | 0.1825 | 0.0718 | 0.309 | 0.702 |
| VSMC_myofibroblast | 0.1777 | 0.4251 | 0.2474 | 0.309 | 0.702 |
| → macrophages |  |  |  |  |  |
| mast → plasma | 0.0655 | 0.0382 | -0.0273 | 0.310 | 0.702 |
| VSMC_myofibroblast | 0.1456 | 0.3890 | 0.2434 | 0.311 | 0.702 |
| → Treg |  |  |  |  |  |
| NK → CD8T | 0.2469 | 0.3155 | 0.0686 | 0.313 | 0.702 |
| CD8T → | 2.8328 | 2.3013 | -0.5316 | 0.320 | 0.713 |
| vasculature |  |  |  |  |  |
| plasma → mast | -0.0086 | 0.0075 | 0.0161 | 0.327 | 0.716 |
| macrophages → | 1.8684 | 1.4704 | -0.3980 | 0.330 | 0.716 |
| vasculature |  |  |  |  |  |
| CAF → TFH | -0.0077 | -0.0055 | 0.0021 | 0.331 | 0.716 |
| CD8T → gdT | 1.1923 | 1.6453 | 0.4530 | 0.331 | 0.716 |
| DC → CD4T | 2.2127 | 1.8332 | -0.3795 | 0.335 | 0.716 |
| B → TFH | 5.1196 | 3.6144 | -1.5052 | 0.337 | 0.716 |
| B → DC | 8.9801 | 7.3432 | -1.6369 | 0.341 | 0.716 |
| CD4T → | 2.2006 | 1.4628 | -0.7377 | 0.342 | 0.716 |
| granulocyte |  |  |  |  |  |
| gdT → CD4T | 0.1352 | 0.1739 | 0.0387 | 0.345 | 0.716 |
| Treg → granulocyte | 0.3260 | 0.2124 | -0.1136 | 0.347 | 0.716 |
| CD4T → Treg | 10.1896 | 12.5242 | 2.3346 | 0.348 | 0.716 |
| VSMC_myofibroblast | 0.4000 | 0.6032 | 0.2032 | 0.349 | 0.716 |
| → vasculature |  |  |  |  |  |
| Treg → gdT | 0.5537 | 0.6892 | 0.1355 | 0.351 | 0.716 |
| macrophages → | 1.4595 | 1.0913 | -0.3682 | 0.360 | 0.721 |
| LEC |  |  |  |  |  |
| CAF → stromal | 0.1032 | 0.0731 | -0.0301 | 0.361 | 0.721 |
| VSMC_myofibroblast | -0.0213 | -0.0123 | 0.0090 | 0.362 | 0.721 |
| → FDC |  |  |  |  |  |
| NK → Treg | 0.2142 | 0.2742 | 0.0599 | 0.366 | 0.721 |
| macrophages → gdT | 0.3626 | 0.4747 | 0.1122 | 0.369 | 0.721 |
| B → | -0.3251 | 0.2047 | 0.5298 | 0.369 | 0.721 |
| VSMC_myofibroblast |  |  |  |  |  |
| VSMC_myofibroblast | 0.7175 | 1.0544 | 0.3369 | 0.371 | 0.721 |
| → |  |  |  |  |  |
| VSMC_myofibroblast |  |  |  |  |  |
| CD4T → B | 9.6399 | 11.3050 | 1.6651 | 0.372 | 0.721 |
| CD8T → B | 4.7519 | 4.0676 | -0.6843 | 0.376 | 0.725 |
| DC → CAF | 0.1447 | 0.0835 | -0.0612 | 0.381 | 0.729 |
| CAF → mast | 0.0085 | -0.0039 | -0.0124 | 0.385 | 0.733 |
| FDC → LEC | -0.1168 | -0.0380 | 0.0788 | 0.390 | 0.733 |
| VSMC_myofibroblast | 0.2455 | 0.4751 | 0.2297 | 0.390 | 0.733 |
| → CD4T |  |  |  |  |  |
| DC → LEC | 0.5825 | 0.4608 | -0.1217 | 0.397 | 0.742 |
| VSMC_myofibroblast | 0.1279 | 0.2899 | 0.1619 | 0.402 | 0.746 |
| → DC |  |  |  |  |  |
| CAF → CD4T | 0.0720 | 0.0552 | -0.0168 | 0.407 | 0.751 |
| FDC → Treg | 0.0866 | 0.0088 | -0.0777 | 0.415 | 0.761 |
| plasma → TFH | -0.0237 | -0.0144 | 0.0093 | 0.423 | 0.769 |
| LEC → CD4T | 0.4641 | 0.3743 | -0.0898 | 0.424 | 0.769 |
| B → CD4T | 9.8705 | 8.3895 | -1.4810 | 0.431 | 0.776 |
| CAF → LEC | -0.0112 | -0.0074 | 0.0037 | 0.433 | 0.777 |
| NK → mast | -0.0018 | 0.0059 | 0.0078 | 0.438 | 0.780 |
| B → granulocyte | 2.0885 | 1.4342 | -0.6543 | 0.440 | 0.780 |
| CD8T → plasma | 3.0850 | 2.5930 | -0.4920 | 0.445 | 0.784 |
| stromal → | 2.5864 | 3.0408 | 0.4543 | 0.453 | 0.789 |
| macrophages |  |  |  |  |  |
| CD8T → TFH | 0.0965 | 0.1831 | 0.0866 | 0.453 | 0.789 |
| CD4T → CD8T | 11.3186 | 13.0613 | 1.7427 | 0.457 | 0.789 |
| B → plasma | 4.2174 | 3.2979 | -0.9196 | 0.460 | 0.789 |
| gdT → granulocyte | -0.0038 | -0.0055 | -0.0017 | 0.460 | 0.789 |
| mast → vasculature | 0.0731 | 0.0530 | -0.0201 | 0.468 | 0.795 |
| B → gdT | 1.2005 | 1.6284 | 0.4279 | 0.469 | 0.795 |
| LEC → gdT | 0.0503 | 0.0288 | -0.0215 | 0.474 | 0.798 |
| DC → B | 1.9816 | 1.7128 | -0.2688 | 0.479 | 0.798 |
| LEC → stromal | 0.6265 | 0.5142 | -0.1123 | 0.482 | 0.798 |
| CD4T → CD4T | 10.5040 | 12.1298 | 1.6258 | 0.485 | 0.798 |
| stromal → Treg | 2.0974 | 2.3943 | 0.2969 | 0.485 | 0.798 |
| plasma → CAF | 0.0373 | 0.0707 | 0.0333 | 0.486 | 0.798 |
| LEC → | 0.7226 | 0.5827 | -0.1398 | 0.503 | 0.819 |
| macrophages |  |  |  |  |  |
| LEC →<br>VSMC_myofibroblast | -0.0113 | -0.0178 | -0.0065 | 0.504 | 0.819 |
| stromal → LEC | 1.6314 | 1.3807 | -0.2507 | 0.513 | 0.821 |
| vasculature →<br>vasculature | 2.2148 | 2.4176 | 0.2029 | 0.513 | 0.821 |
| NK → TFH | -0.0109 | -0.0071 | 0.0038 | 0.514 | 0.821 |
| VSMC_myofibroblast<br>→ granulocyte | -0.0151 | -0.0112 | 0.0039 | 0.518 | 0.821 |
| plasma →<br>granulocyte | 0.0273 | 0.0043 | -0.0230 | 0.521 | 0.821 |
| stromal → NK | 0.7250 | 0.8524 | 0.1274 | 0.528 | 0.821 |
| LEC → FDC | -0.0259 | -0.0069 | 0.0190 | 0.528 | 0.821 |
| Treg → B | 2.2187 | 2.5509 | 0.3322 | 0.531 | 0.821 |
| CAF → plasma | 0.0521 | 0.0343 | -0.0178 | 0.532 | 0.821 |
| macrophages →<br>CD4T | 2.3184 | 2.0746 | -0.2438 | 0.536 | 0.821 |
| mast → mast | 0.0361 | 0.0500 | 0.0139 | 0.537 | 0.821 |
| NK → CD4T | 0.2592 | 0.3000 | 0.0408 | 0.538 | 0.821 |
| LEC → DC | 0.3984 | 0.3382 | -0.0602 | 0.540 | 0.821 |
| DC → plasma | 0.7166 | 0.5996 | -0.1170 | 0.542 | 0.821 |
| TFH → stromal | 0.0165 | 0.0322 | 0.0157 | 0.546 | 0.822 |
| VSMC_myofibroblast<br>→ CD8T | 0.3084 | 0.4851 | 0.1768 | 0.548 | 0.822 |
| CAF → NK | 0.0040 | -0.0040 | -0.0080 | 0.554 | 0.825 |
| macrophages →<br>granulocyte | 1.4077 | 1.1319 | -0.2759 | 0.555 | 0.825 |
| stromal → stromal | 2.6663 | 3.0825 | 0.4162 | 0.560 | 0.828 |
| stromal → DC | 1.9368 | 2.1571 | 0.2204 | 0.563 | 0.828 |
| CD8T → NK | 1.6675 | 1.8836 | 0.2161 | 0.566 | 0.830 |
| NK → CAF | 0.0112 | 0.0015 | -0.0098 | 0.570 | 0.830 |
| stromal →<br>VSMC_myofibroblast | 0.8325 | 1.0578 | 0.2253 | 0.572 | 0.830 |
| Treg → FDC | 0.0739 | 0.0351 | -0.0389 | 0.585 | 0.841 |
| gdT →<br>VSMC_myofibroblast | -0.0027 | 0.0040 | 0.0067 | 0.585 | 0.841 |
| granulocyte → CAF | -0.0082 | -0.0058 | 0.0024 | 0.589 | 0.841 |
| plasma → DC | 0.4942 | 0.4186 | -0.0755 | 0.592 | 0.841 |
| VSMC_myofibroblast<br>→ B | 0.2567 | 0.3786 | 0.1219 | 0.593 | 0.841 |
| B → CD8T | 7.5371 | 6.6969 | -0.8402 | 0.596 | 0.841 |
| stromal → FDC | -0.0274 | -0.0102 | 0.0171 | 0.600 | 0.844 |
| stromal →<br>granulocyte | 1.5802 | 1.3583 | -0.2219 | 0.613 | 0.844 |
| TFH → CD4T | 0.0442 | 0.0532 | 0.0090 | 0.614 | 0.844 |
| macrophages →<br>TFH | -0.0092 | 0.0048 | 0.0140 | 0.617 | 0.844 |
| DC → NK | 0.5480 | 0.6276 | 0.0796 | 0.621 | 0.844 |
| VSMC_myofibroblast<br>→ stromal | 0.3996 | 0.5114 | 0.1118 | 0.624 | 0.844 |
| CD4T → stromal | 8.9838 | 10.1661 | 1.1823 | 0.625 | 0.844 |
| B → LEC | 1.3786 | 1.7037 | 0.3251 | 0.627 | 0.844 |
| DC → Treg | 1.9519 | 1.7424 | -0.2095 | 0.630 | 0.844 |
| LEC → Treg | 0.2624 | 0.2258 | -0.0367 | 0.631 | 0.844 |
| CD8T → LEC | 0.8270 | 0.6981 | -0.1289 | 0.631 | 0.844 |
| NK → stromal | 0.2422 | 0.2805 | 0.0383 | 0.637 | 0.844 |
| B → NK | 1.6966 | 2.0093 | 0.3127 | 0.640 | 0.844 |
| FDC → plasma | -0.0133 | -0.0274 | -0.0141 | 0.645 | 0.844 |
| plasma → gdT | 0.1408 | 0.1744 | 0.0335 | 0.645 | 0.844 |
| macrophages → Treg | 1.5035 | 1.6262 | 0.1227 | 0.649 | 0.844 |
| vasculature → VSMC_myofibroblast | 0.6228 | 0.7334 | 0.1105 | 0.650 | 0.844 |
| mast → VSMC_myofibroblast | -0.0047 | -0.0040 | 0.0007 | 0.652 | 0.844 |
| FDC → VSMC_myofibroblast | -0.1270 | -0.0908 | 0.0362 | 0.655 | 0.844 |
| B → vasculature | 3.4809 | 2.9396 | -0.5413 | 0.660 | 0.844 |
| VSMC_myofibroblast → NK | 0.0048 | 0.0203 | 0.0156 | 0.663 | 0.844 |
| LEC → NK | 0.1051 | 0.1279 | 0.0228 | 0.663 | 0.844 |
| Treg → mast | 0.1238 | 0.1015 | -0.0222 | 0.668 | 0.844 |
| gdT → mast | -0.0055 | -0.0068 | -0.0013 | 0.670 | 0.844 |
| TFH → NK | -0.0196 | -0.0097 | 0.0098 | 0.670 | 0.844 |
| CD4T → CAF | 0.9633 | 0.8119 | -0.1513 | 0.674 | 0.844 |
| B → CAF | 0.4320 | 0.5517 | 0.1197 | 0.676 | 0.844 |
| VSMC_myofibroblast → plasma | 0.1781 | 0.2305 | 0.0524 | 0.677 | 0.844 |
| macrophages → stromal | 2.2978 | 2.1258 | -0.1720 | 0.678 | 0.844 |
| CAF → B | 0.0607 | 0.0521 | -0.0086 | 0.679 | 0.844 |
| NK → LEC | 0.0863 | 0.0733 | -0.0130 | 0.683 | 0.846 |
| NK → DC | 0.2092 | 0.2327 | 0.0235 | 0.690 | 0.851 |
| TFH → gdT | -0.0125 | -0.0153 | -0.0028 | 0.707 | 0.868 |
| Treg → DC | 2.3040 | 2.5314 | 0.2273 | 0.711 | 0.870 |
| DC → VSMC_myofibroblast | 0.2115 | 0.2788 | 0.0673 | 0.722 | 0.879 |
| CD4T → plasma | 5.7259 | 5.2870 | -0.4389 | 0.726 | 0.882 |
| Treg → plasma | 1.4173 | 1.2892 | -0.1281 | 0.740 | 0.885 |
| macrophages → macrophages | 2.0569 | 1.9151 | -0.1417 | 0.744 | 0.885 |
| CD8T → FDC | -0.0703 | -0.0354 | 0.0350 | 0.745 | 0.885 |
| TFH → CD8T | 0.0250 | 0.0359 | 0.0109 | 0.745 | 0.885 |
| B → macrophages | 4.6043 | 4.2060 | -0.3983 | 0.746 | 0.885 |
| mast → gdT | -0.0056 | -0.0065 | -0.0010 | 0.746 | 0.885 |
| stromal → vasculature | 2.8076 | 2.6271 | -0.1805 | 0.749 | 0.886 |
| macrophages → B | 2.0545 | 1.9370 | -0.1175 | 0.755 | 0.886 |
| DC → FDC | 0.1695 | 0.1419 | -0.0277 | 0.756 | 0.886 |
| FDC → CD8T | -0.0038 | -0.0350 | -0.0312 | 0.761 | 0.886 |
| LEC → plasma | 0.0670 | 0.0846 | 0.0176 | 0.767 | 0.886 |
| stromal → CD8T | 2.7442 | 2.8897 | 0.1456 | 0.768 | 0.886 |
| CD4T → FDC | 0.1219 | 0.0296 | -0.0924 | 0.770 | 0.886 |
| vasculature → Treg | 0.8774 | 0.8287 | -0.0487 | 0.770 | 0.886 |
| TFH → plasma | -0.0078 | -0.0110 | -0.0031 | 0.777 | 0.886 |
| Treg → vasculature | 0.9767 | 1.0676 | 0.0909 | 0.777 | 0.886 |
| LEC → CD8T | 0.3773 | 0.3472 | -0.0301 | 0.779 | 0.886 |
| macrophages → plasma | 1.3345 | 1.4189 | 0.0844 | 0.781 | 0.886 |
| Treg → Treg | 2.1120 | 2.2804 | 0.1684 | 0.784 | 0.886 |
| TFH → B | 0.0665 | 0.0604 | -0.0061 | 0.797 | 0.898 |
| macrophages → CAF | 0.1996 | 0.1833 | -0.0164 | 0.818 | 0.918 |
| mast → NK | -0.0027 | -0.0011 | 0.0017 | 0.825 | 0.920 |
| gdT → Treg | 0.1229 | 0.1291 | 0.0063 | 0.826 | 0.920 |
| LEC → CAF | 0.0038 | -0.0017 | -0.0054 | 0.833 | 0.925 |
| NK → macrophages | 0.2573 | 0.2750 | 0.0177 | 0.850 | 0.937 |
| DC → TFH | 0.3359 | 0.3644 | 0.0285 | 0.852 | 0.937 |
| Treg → CD4T | 2.7842 | 2.9002 | 0.1159 | 0.856 | 0.937 |
| FDC → NK | -0.0805 | -0.0715 | 0.0091 | 0.856 | 0.937 |
| NK → B | 0.2384 | 0.2515 | 0.0131 | 0.858 | 0.937 |
| vasculature → NK | 0.3053 | 0.3205 | 0.0152 | 0.867 | 0.940 |
| NK → vasculature | 0.1339 | 0.1420 | 0.0081 | 0.870 | 0.940 |
| FDC → gdT | -0.0255 | -0.0280 | -0.0024 | 0.872 | 0.940 |
| VSMC_myofibroblast → gdT | -0.0048 | -0.0023 | 0.0025 | 0.874 | 0.940 |
| CD4T → vasculature | 5.1112 | 5.3009 | 0.1897 | 0.892 | 0.949 |
| B → Treg | 7.1337 | 7.3266 | 0.1929 | 0.899 | 0.949 |
| stromal → B | 2.8371 | 2.7715 | -0.0656 | 0.899 | 0.949 |
| B → stromal | 6.4043 | 6.1962 | -0.2081 | 0.901 | 0.949 |
| CD4T → macrophages | 8.5122 | 8.7248 | 0.2127 | 0.904 | 0.949 |
| LEC → LEC | 0.9461 | 0.9893 | 0.0432 | 0.904 | 0.949 |
| FDC → CD4T | 0.0402 | 0.0366 | -0.0036 | 0.907 | 0.949 |
| CD8T → VSMC_myofibroblast | 0.8260 | 0.8842 | 0.0582 | 0.908 | 0.949 |
| CAF → VSMC_myofibroblast | 0.0011 | 0.0034 | 0.0024 | 0.910 | 0.949 |
| Treg → LEC | 0.1989 | 0.2153 | 0.0163 | 0.912 | 0.949 |
| macrophages → NK | 0.8030 | 0.7744 | -0.0286 | 0.918 | 0.951 |
| macrophages → CD8T | 2.0191 | 1.9860 | -0.0332 | 0.927 | 0.958 |
| CD4T → LEC | 1.5646 | 1.5166 | -0.0480 | 0.932 | 0.958 |
| granulocyte → LEC | 0.7370 | 0.7115 | -0.0255 | 0.935 | 0.958 |
| Treg → macrophages | 1.8698 | 1.9077 | 0.0378 | 0.936 | 0.958 |
| LEC → B | 0.4061 | 0.4126 | 0.0065 | 0.955 | 0.974 |
| NK → FDC | -0.0224 | -0.0220 | 0.0004 | 0.959 | 0.975 |
| stromal → plasma | 1.5150 | 1.5332 | 0.0183 | 0.962 | 0.975 |
| CAF → granulocyte | 0.0094 | 0.0089 | -0.0005 | 0.973 | 0.983 |
| VSMC_myofibroblast → mast | -0.0048 | -0.0049 | -0.0001 | 0.980 | 0.983 |
| Treg → stromal | 2.2701 | 2.2861 | 0.0160 | 0.980 | 0.983 |

| Cell type pair | LN benign | LN met-adjacent | Effect size | p-value | adjusted p-value |
| --- | --- | --- | --- | --- | --- |
| stromal → CD4T | 3.0179 | 3.0152 | -0.0026 | 0.996 | 0.996 |

#### S9.3 HNSCC spicyR benchmark

**Table S7:**
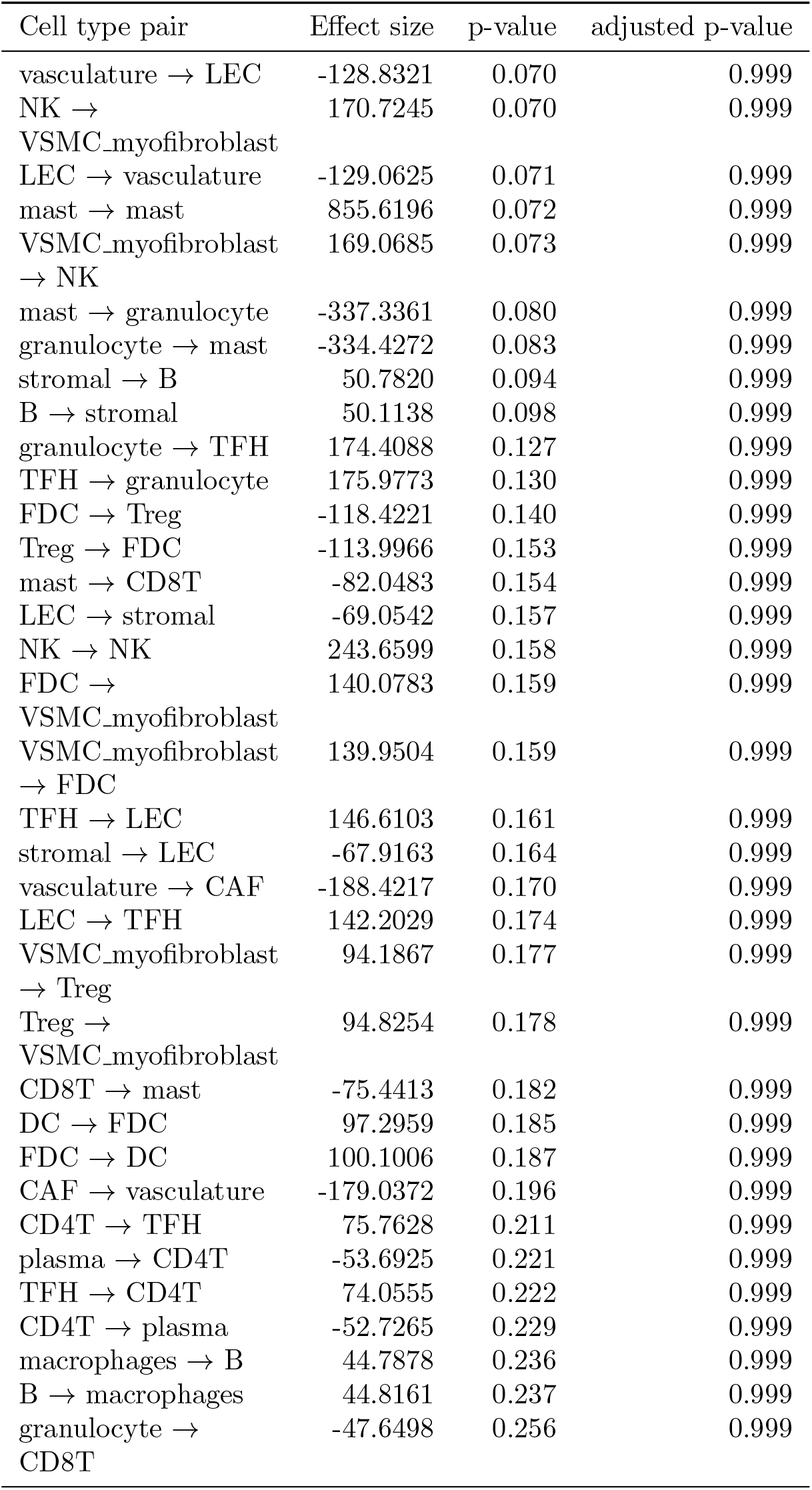
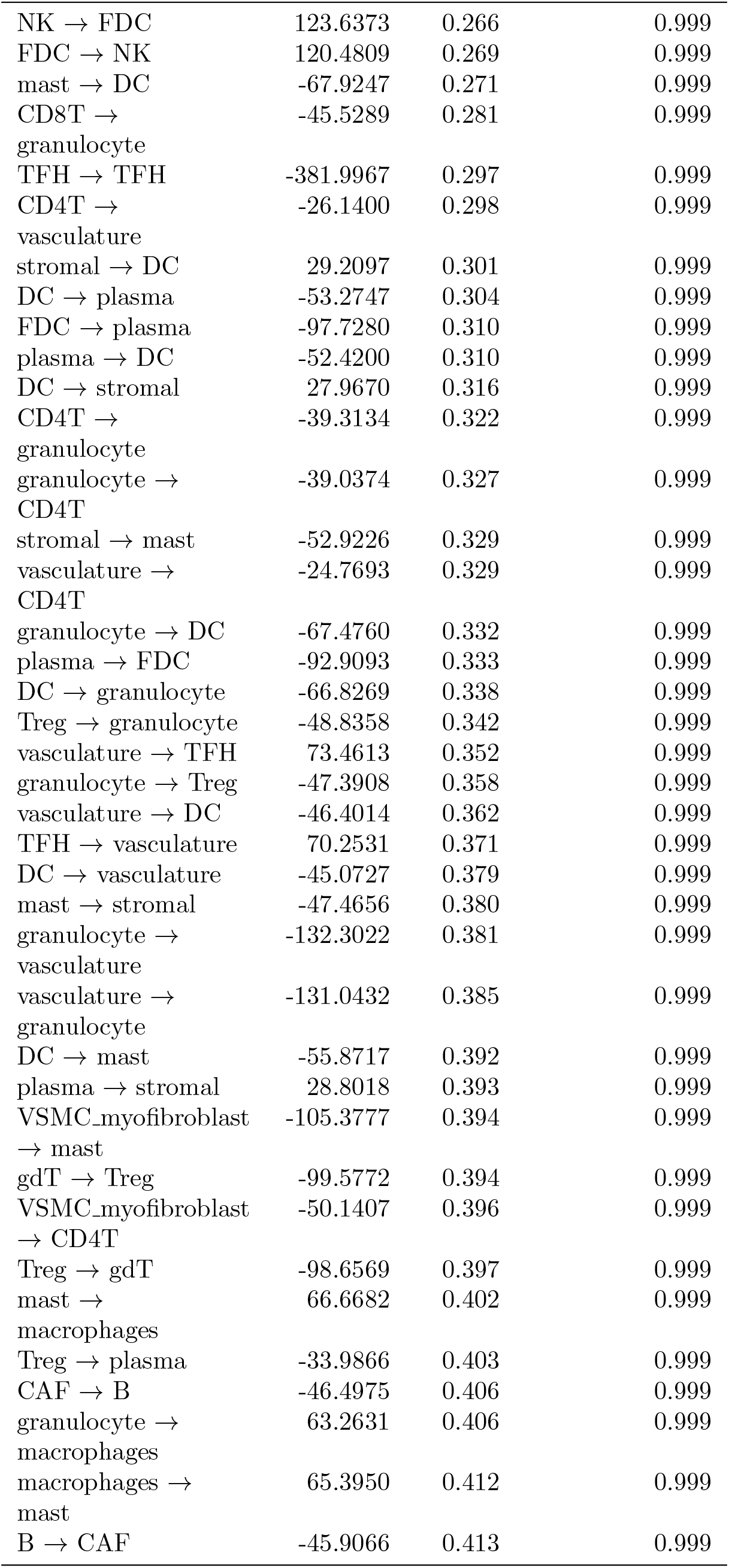
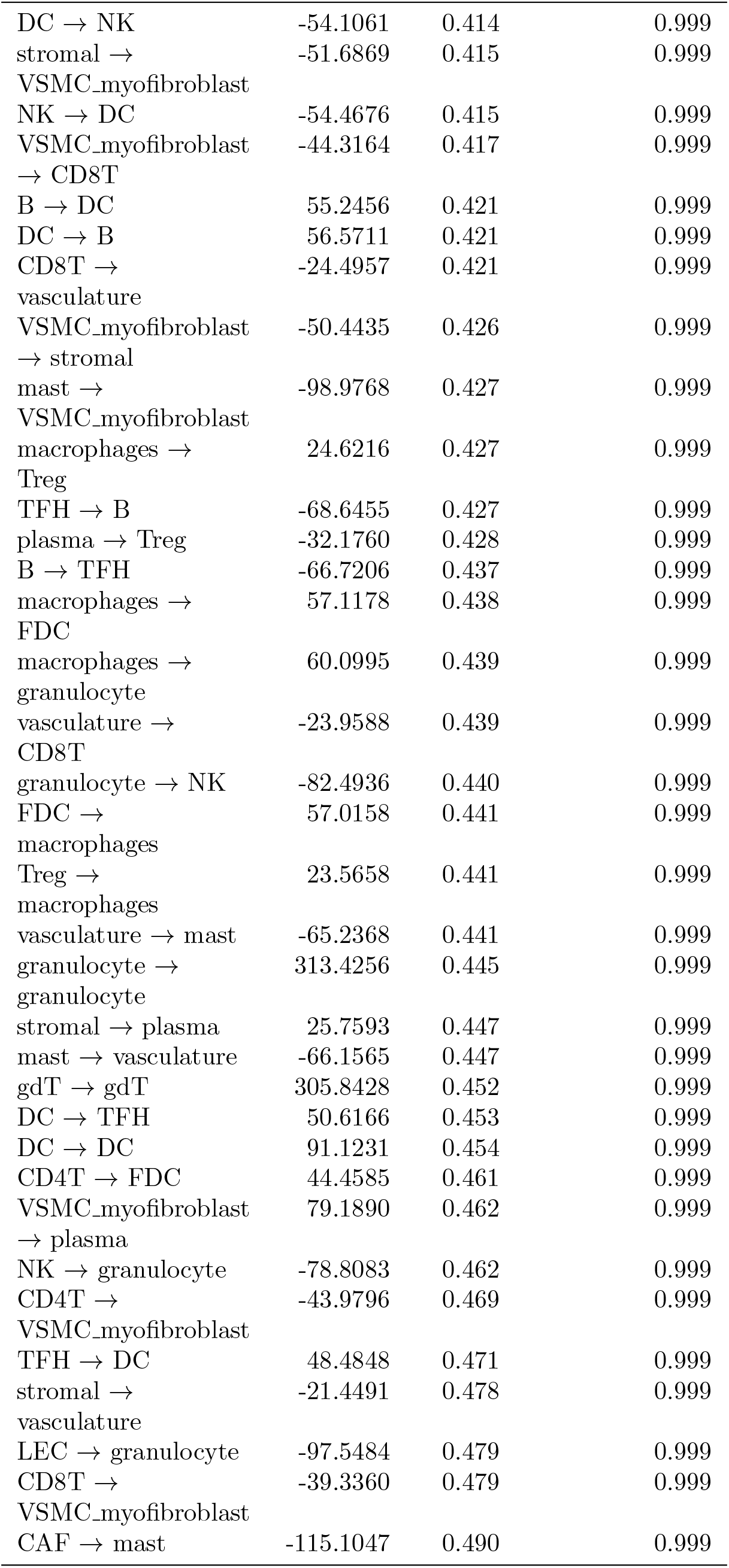
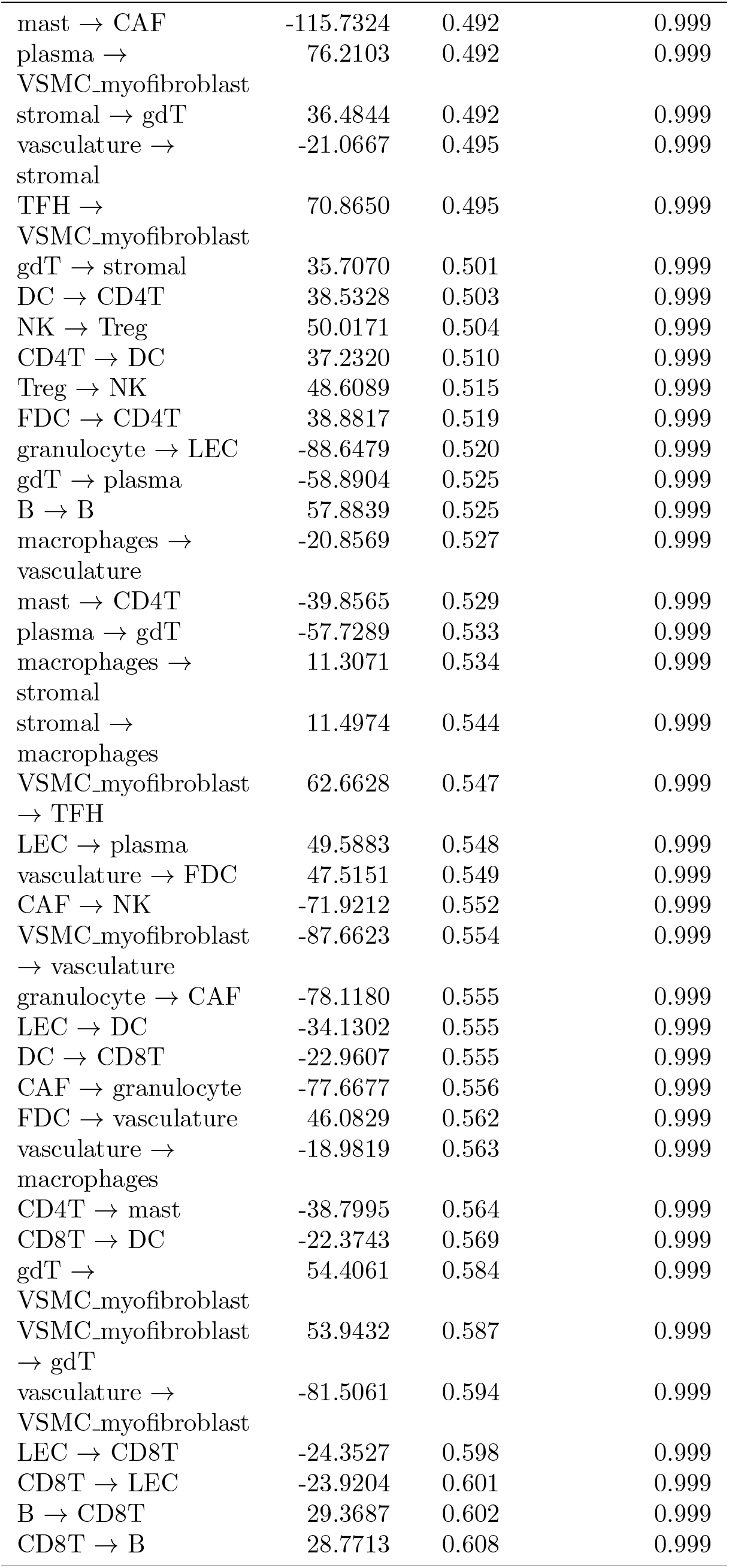
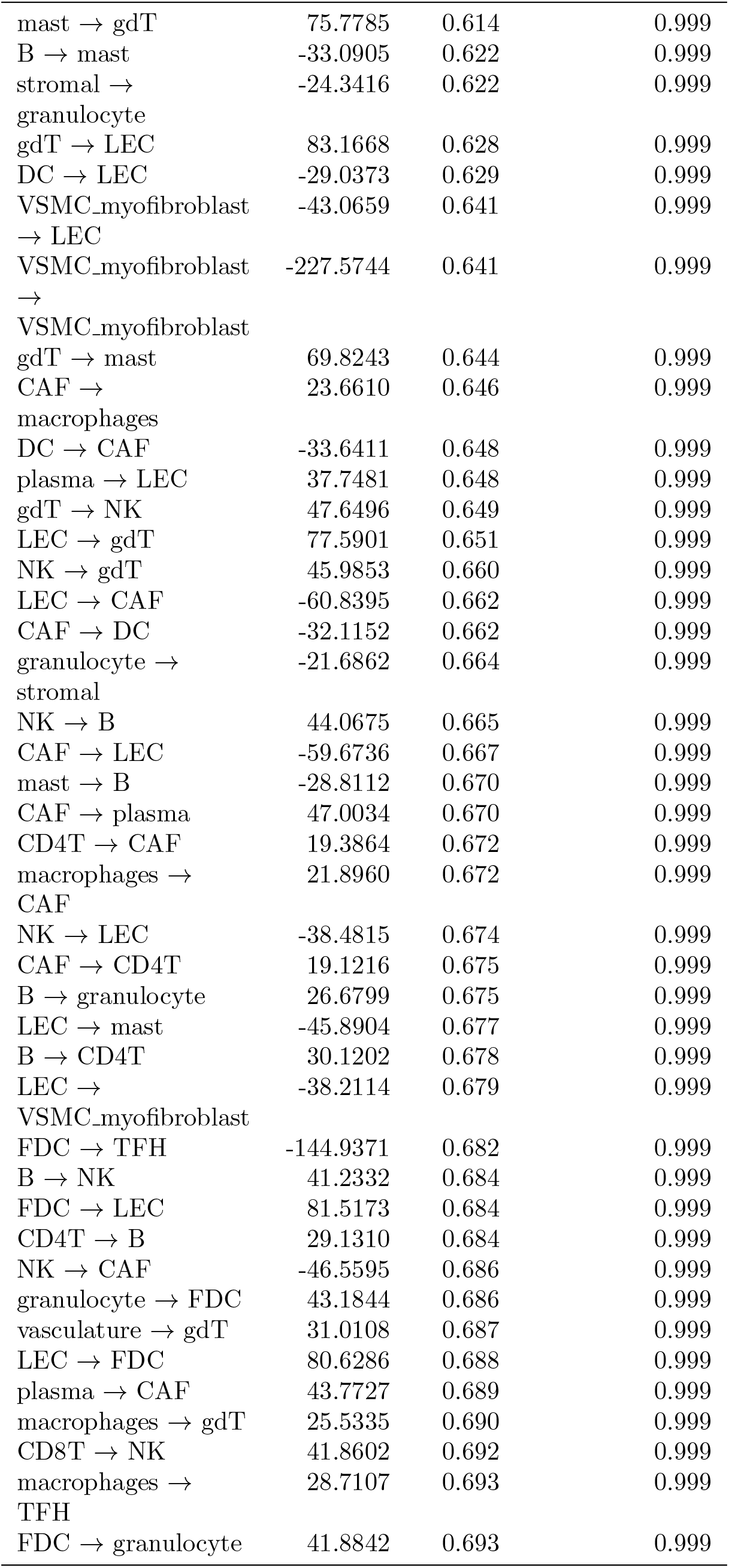
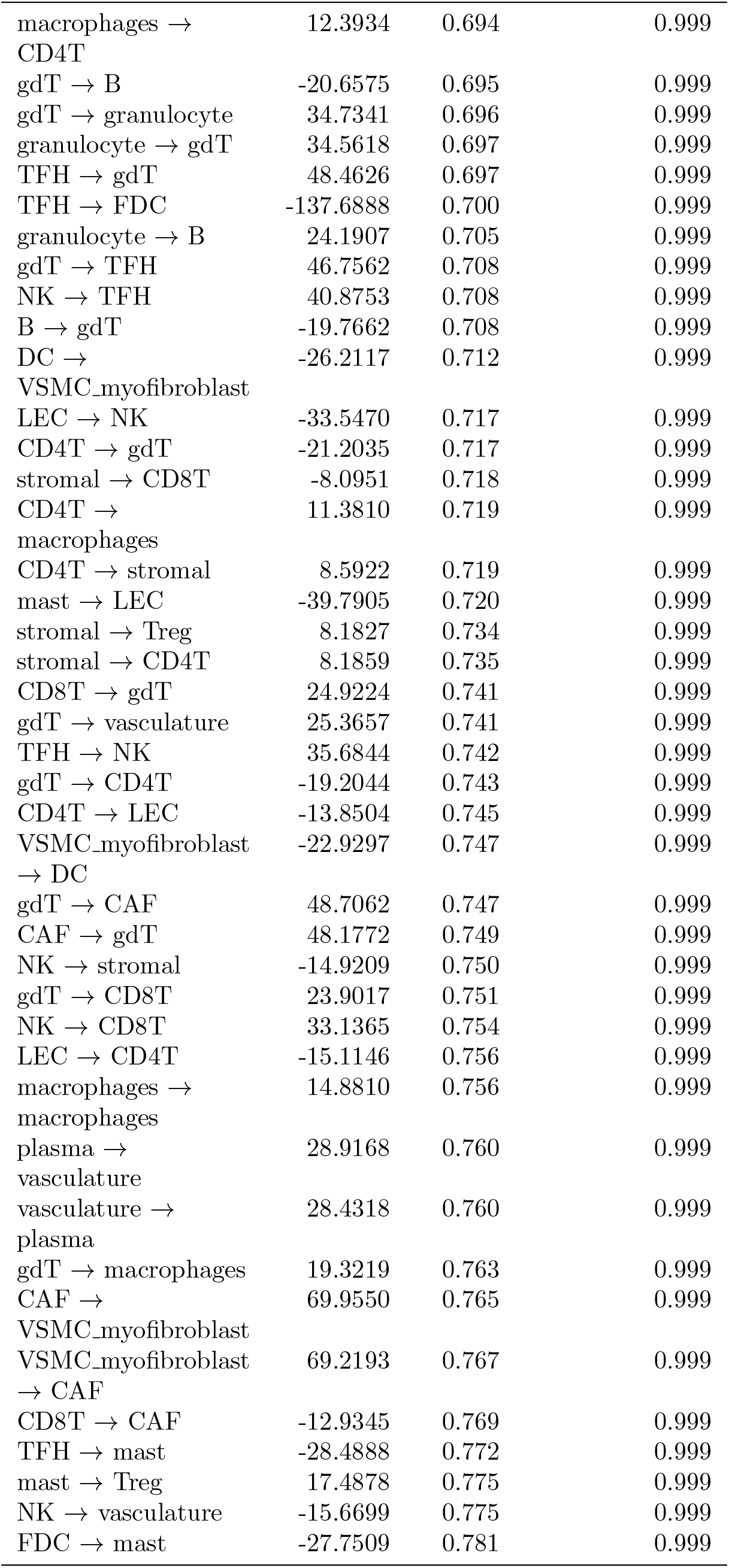
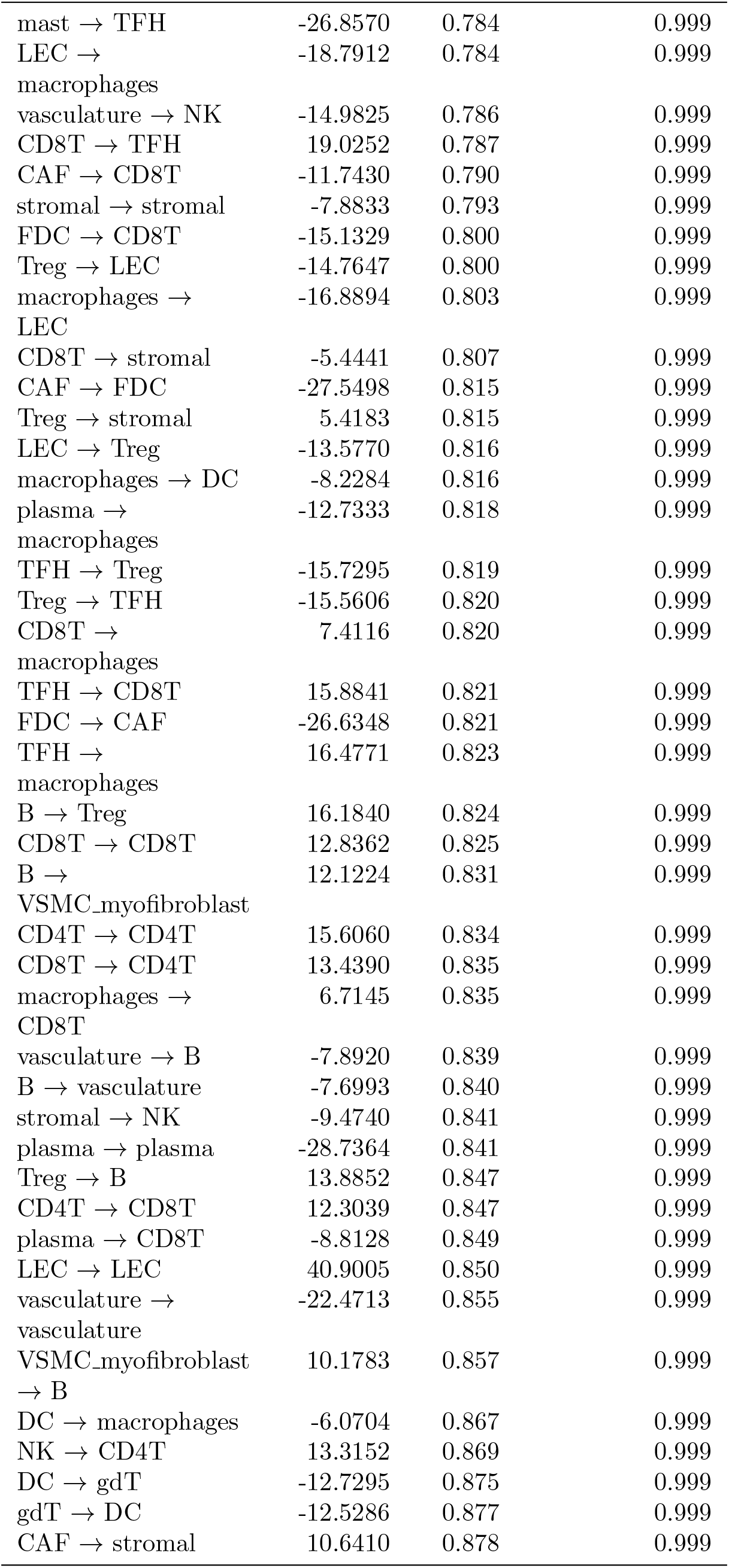
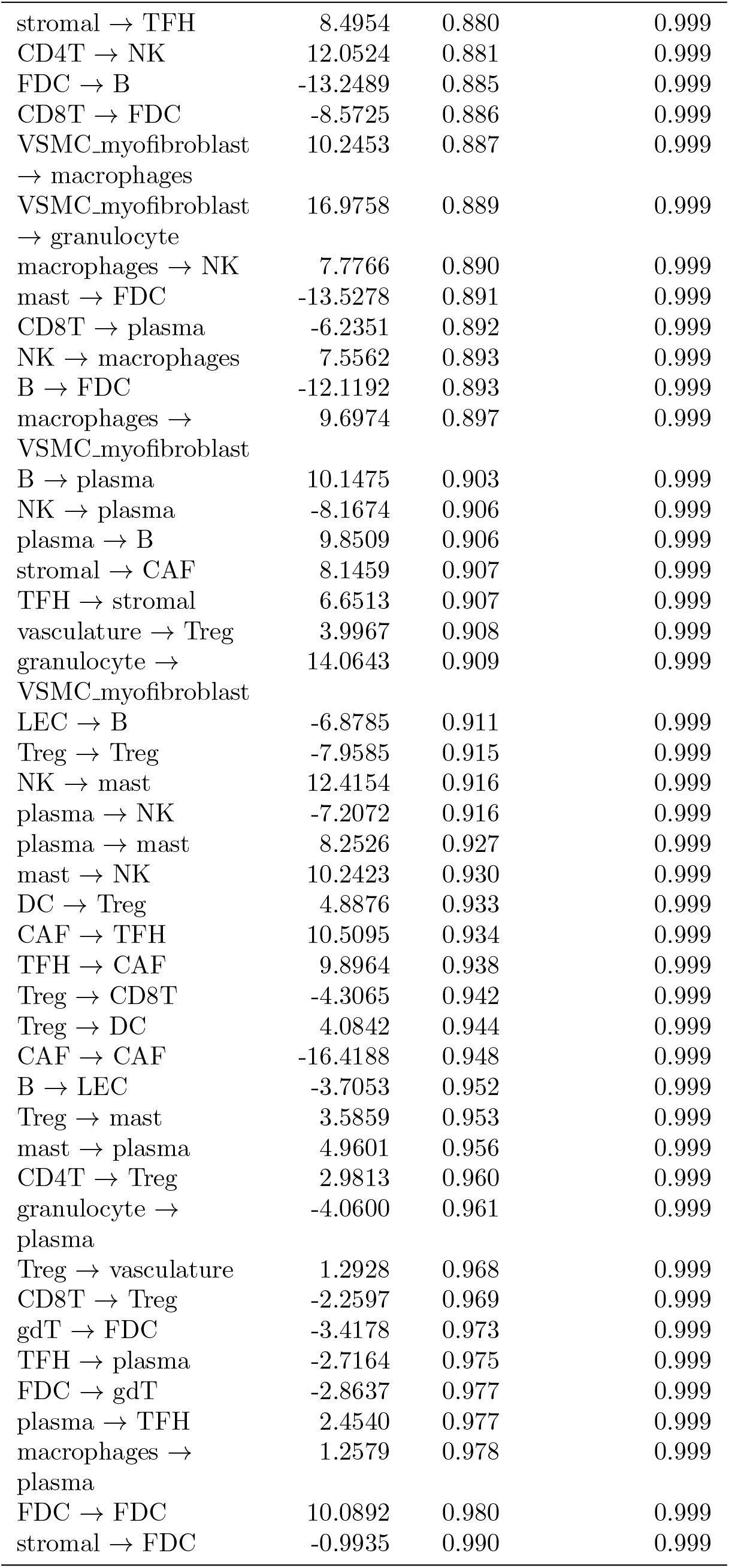
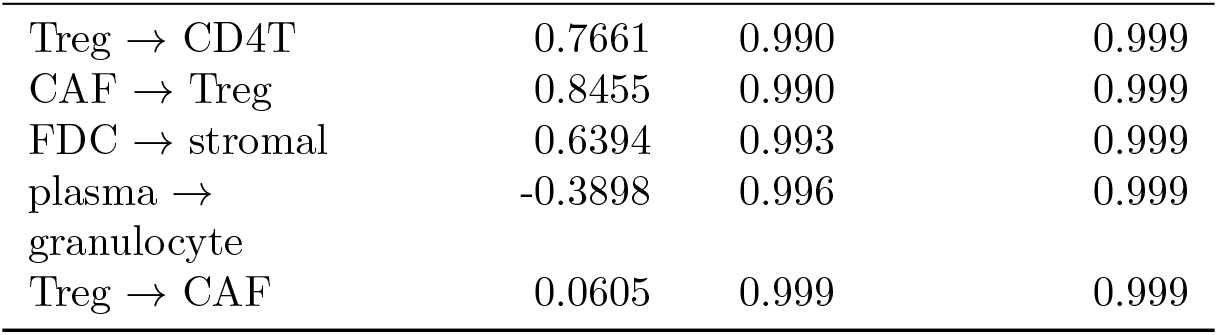
HNSCC TMA spicyR results (benchmark context).

| Cell type pair | Effect size | p-value | adjusted p-value |
| --- | --- | --- | --- |
| vasculature → LEC | -128.8321 | 0.070 | 0.999 |
| NK → | 170.7245 | 0.070 | 0.999 |
| VSMC_myofibroblast |  |  |  |
| LEC → vasculature | -129.0625 | 0.071 | 0.999 |
| mast → mast | 855.6196 | 0.072 | 0.999 |
| VSMC_myofibroblast | 169.0685 | 0.073 | 0.999 |
| → NK |  |  |  |
| mast → granulocyte | -337.3361 | 0.080 | 0.999 |
| granulocyte → mast | -334.4272 | 0.083 | 0.999 |
| stromal → B | 50.7820 | 0.094 | 0.999 |
| B → stromal | 50.1138 | 0.098 | 0.999 |
| granulocyte → TFH | 174.4088 | 0.127 | 0.999 |
| TFH → granulocyte | 175.9773 | 0.130 | 0.999 |
| FDC → Treg | -118.4221 | 0.140 | 0.999 |
| Treg → FDC | -113.9966 | 0.153 | 0.999 |
| mast → CD8T | -82.0483 | 0.154 | 0.999 |
| LEC → stromal | -69.0542 | 0.157 | 0.999 |
| NK → NK | 243.6599 | 0.158 | 0.999 |
| FDC → | 140.0783 | 0.159 | 0.999 |
| VSMC_myofibroblast |  |  |  |
| VSMC_myofibroblast | 139.9504 | 0.159 | 0.999 |
| → FDC |  |  |  |
| TFH → LEC | 146.6103 | 0.161 | 0.999 |
| stromal → LEC | -67.9163 | 0.164 | 0.999 |
| vasculature → CAF | -188.4217 | 0.170 | 0.999 |
| LEC → TFH | 142.2029 | 0.174 | 0.999 |
| VSMC_myofibroblast | 94.1867 | 0.177 | 0.999 |
| → Treg |  |  |  |
| Treg → | 94.8254 | 0.178 | 0.999 |
| VSMC_myofibroblast |  |  |  |
| CD8T → mast | -75.4413 | 0.182 | 0.999 |
| DC → FDC | 97.2959 | 0.185 | 0.999 |
| FDC → DC | 100.1006 | 0.187 | 0.999 |
| CAF → vasculature | -179.0372 | 0.196 | 0.999 |
| CD4T → TFH | 75.7628 | 0.211 | 0.999 |
| plasma → CD4T | -53.6925 | 0.221 | 0.999 |
| TFH → CD4T | 74.0555 | 0.222 | 0.999 |
| CD4T → plasma | -52.7265 | 0.229 | 0.999 |
| macrophages → B | 44.7878 | 0.236 | 0.999 |
| B → macrophages | 44.8161 | 0.237 | 0.999 |
| granulocyte → | -47.6498 | 0.256 | 0.999 |
| CD8T |  |  |  |
| NK → FDC | 123.6373 | 0.266 | 0.999 |
| FDC → NK | 120.4809 | 0.269 | 0.999 |
| mast → DC | -67.9247 | 0.271 | 0.999 |
| CD8T → granulocyte | -45.5289 | 0.281 | 0.999 |
| TFH → TFH | -381.9967 | 0.297 | 0.999 |
| CD4T → vasculature | -26.1400 | 0.298 | 0.999 |
| stromal → DC | 29.2097 | 0.301 | 0.999 |
| DC → plasma | -53.2747 | 0.304 | 0.999 |
| FDC → plasma | -97.7280 | 0.310 | 0.999 |
| plasma → DC | -52.4200 | 0.310 | 0.999 |
| DC → stromal | 27.9670 | 0.316 | 0.999 |
| CD4T → granulocyte | -39.3134 | 0.322 | 0.999 |
| granulocyte → CD4T | -39.0374 | 0.327 | 0.999 |
| stromal → mast | -52.9226 | 0.329 | 0.999 |
| vasculature → CD4T | -24.7693 | 0.329 | 0.999 |
| granulocyte → DC | -67.4760 | 0.332 | 0.999 |
| plasma → FDC | -92.9093 | 0.333 | 0.999 |
| DC → granulocyte | -66.8269 | 0.338 | 0.999 |
| Treg → granulocyte | -48.8358 | 0.342 | 0.999 |
| vasculature → TFH | 73.4613 | 0.352 | 0.999 |
| granulocyte → Treg | -47.3908 | 0.358 | 0.999 |
| vasculature → DC | -46.4014 | 0.362 | 0.999 |
| TFH → vasculature | 70.2531 | 0.371 | 0.999 |
| DC → vasculature | -45.0727 | 0.379 | 0.999 |
| mast → stromal | -47.4656 | 0.380 | 0.999 |
| granulocyte → vasculature | -132.3022 | 0.381 | 0.999 |
| vasculature → granulocyte | -131.0432 | 0.385 | 0.999 |
| DC → mast | -55.8717 | 0.392 | 0.999 |
| plasma → stromal | 28.8018 | 0.393 | 0.999 |
| VSMC_myofibroblast → mast | -105.3777 | 0.394 | 0.999 |
| gdT → Treg | -99.5772 | 0.394 | 0.999 |
| VSMC_myofibroblast → CD4T | -50.1407 | 0.396 | 0.999 |
| Treg → gdT | -98.6569 | 0.397 | 0.999 |
| mast → macrophages | 66.6682 | 0.402 | 0.999 |
| Treg → plasma | -33.9866 | 0.403 | 0.999 |
| CAF → B | -46.4975 | 0.406 | 0.999 |
| granulocyte → macrophages | 63.2631 | 0.406 | 0.999 |
| macrophages → mast | 65.3950 | 0.412 | 0.999 |
| B → CAF | -45.9066 | 0.413 | 0.999 |
| DC → NK | -54.1061 | 0.414 | 0.999 |
| stromal → | -51.6869 | 0.415 | 0.999 |
| VSMC_myofibroblast |  |  |  |
| NK → DC | -54.4676 | 0.415 | 0.999 |
| VSMC_myofibroblast | -44.3164 | 0.417 | 0.999 |
| → CD8T |  |  |  |
| B → DC | 55.2456 | 0.421 | 0.999 |
| DC → B | 56.5711 | 0.421 | 0.999 |
| CD8T → | -24.4957 | 0.421 | 0.999 |
| vasculature |  |  |  |
| VSMC_myofibroblast | -50.4435 | 0.426 | 0.999 |
| → stromal |  |  |  |
| mast → | -98.9768 | 0.427 | 0.999 |
| VSMC_myofibroblast |  |  |  |
| macrophages → | 24.6216 | 0.427 | 0.999 |
| Treg |  |  |  |
| TFH → B | -68.6455 | 0.427 | 0.999 |
| plasma → Treg | -32.1760 | 0.428 | 0.999 |
| B → TFH | -66.7206 | 0.437 | 0.999 |
| macrophages → | 57.1178 | 0.438 | 0.999 |
| FDC |  |  |  |
| macrophages → | 60.0995 | 0.439 | 0.999 |
| granulocyte |  |  |  |
| vasculature → | -23.9588 | 0.439 | 0.999 |
| CD8T |  |  |  |
| granulocyte → NK | -82.4936 | 0.440 | 0.999 |
| FDC → | 57.0158 | 0.441 | 0.999 |
| macrophages |  |  |  |
| Treg → | 23.5658 | 0.441 | 0.999 |
| macrophages |  |  |  |
| vasculature → mast | -65.2368 | 0.441 | 0.999 |
| granulocyte → | 313.4256 | 0.445 | 0.999 |
| granulocyte |  |  |  |
| stromal → plasma | 25.7593 | 0.447 | 0.999 |
| mast → vasculature | -66.1565 | 0.447 | 0.999 |
| gdT → gdT | 305.8428 | 0.452 | 0.999 |
| DC → TFH | 50.6166 | 0.453 | 0.999 |
| DC → DC | 91.1231 | 0.454 | 0.999 |
| CD4T → FDC | 44.4585 | 0.461 | 0.999 |
| VSMC_myofibroblast | 79.1890 | 0.462 | 0.999 |
| → plasma |  |  |  |
| NK → granulocyte | -78.8083 | 0.462 | 0.999 |
| CD4T → | -43.9796 | 0.469 | 0.999 |
| VSMC_myofibroblast |  |  |  |
| TFH → DC | 48.4848 | 0.471 | 0.999 |
| stromal → | -21.4491 | 0.478 | 0.999 |
| vasculature |  |  |  |
| LEC → granulocyte | -97.5484 | 0.479 | 0.999 |
| CD8T → | -39.3360 | 0.479 | 0.999 |
| VSMC_myofibroblast |  |  |  |
| CAF → mast | -115.1047 | 0.490 | 0.999 |
| mast → CAF | -115.7324 | 0.492 | 0.999 |
| plasma → | 76.2103 | 0.492 | 0.999 |
| VSMC_myofibroblast |  |  |  |
| stromal → gdT | 36.4844 | 0.492 | 0.999 |
| vasculature → | -21.0667 | 0.495 | 0.999 |
| stromal |  |  |  |
| TFH → | 70.8650 | 0.495 | 0.999 |
| VSMC_myofibroblast |  |  |  |
| gdT → stromal | 35.7070 | 0.501 | 0.999 |
| DC → CD4T | 38.5328 | 0.503 | 0.999 |
| NK → Treg | 50.0171 | 0.504 | 0.999 |
| CD4T → DC | 37.2320 | 0.510 | 0.999 |
| Treg → NK | 48.6089 | 0.515 | 0.999 |
| FDC → CD4T | 38.8817 | 0.519 | 0.999 |
| granulocyte → LEC | -88.6479 | 0.520 | 0.999 |
| gdT → plasma | -58.8904 | 0.525 | 0.999 |
| B → B | 57.8839 | 0.525 | 0.999 |
| macrophages → | -20.8569 | 0.527 | 0.999 |
| vasculature |  |  |  |
| mast → CD4T | -39.8565 | 0.529 | 0.999 |
| plasma → gdT | -57.7289 | 0.533 | 0.999 |
| macrophages → | 11.3071 | 0.534 | 0.999 |
| stromal |  |  |  |
| stromal → | 11.4974 | 0.544 | 0.999 |
| macrophages |  |  |  |
| VSMC_myofibroblast | 62.6628 | 0.547 | 0.999 |
| → TFH |  |  |  |
| LEC → plasma | 49.5883 | 0.548 | 0.999 |
| vasculature → FDC | 47.5151 | 0.549 | 0.999 |
| CAF → NK | -71.9212 | 0.552 | 0.999 |
| VSMC_myofibroblast | -87.6623 | 0.554 | 0.999 |
| → vasculature |  |  |  |
| granulocyte → CAF | -78.1180 | 0.555 | 0.999 |
| LEC → DC | -34.1302 | 0.555 | 0.999 |
| DC → CD8T | -22.9607 | 0.555 | 0.999 |
| CAF → granulocyte | -77.6677 | 0.556 | 0.999 |
| FDC → vasculature | 46.0829 | 0.562 | 0.999 |
| vasculature → | -18.9819 | 0.563 | 0.999 |
| macrophages |  |  |  |
| CD4T → mast | -38.7995 | 0.564 | 0.999 |
| CD8T → DC | -22.3743 | 0.569 | 0.999 |
| gdT → | 54.4061 | 0.584 | 0.999 |
| VSMC_myofibroblast |  |  |  |
| VSMC_myofibroblast | 53.9432 | 0.587 | 0.999 |
| → gdT |  |  |  |
| vasculature → | -81.5061 | 0.594 | 0.999 |
| VSMC_myofibroblast |  |  |  |
| LEC → CD8T | -24.3527 | 0.598 | 0.999 |
| CD8T → LEC | -23.9204 | 0.601 | 0.999 |
| B → CD8T | 29.3687 | 0.602 | 0.999 |
| CD8T → B | 28.7713 | 0.608 | 0.999 |
| mast → gdT | 75.7785 | 0.614 | 0.999 |
| B → mast | -33.0905 | 0.622 | 0.999 |
| stromal → granulocyte | -24.3416 | 0.622 | 0.999 |
| gdT → LEC | 83.1668 | 0.628 | 0.999 |
| DC → LEC | -29.0373 | 0.629 | 0.999 |
| VSMC_myofibroblast → LEC | -43.0659 | 0.641 | 0.999 |
| VSMC_myofibroblast → VSMC_myofibroblast | -227.5744 | 0.641 | 0.999 |
| gdT → mast | 69.8243 | 0.644 | 0.999 |
| CAF → macrophages | 23.6610 | 0.646 | 0.999 |
| DC → CAF | -33.6411 | 0.648 | 0.999 |
| plasma → LEC | 37.7481 | 0.648 | 0.999 |
| gdT → NK | 47.6496 | 0.649 | 0.999 |
| LEC → gdT | 77.5901 | 0.651 | 0.999 |
| NK → gdT | 45.9853 | 0.660 | 0.999 |
| LEC → CAF | -60.8395 | 0.662 | 0.999 |
| CAF → DC | -32.1152 | 0.662 | 0.999 |
| granulocyte → stromal | -21.6862 | 0.664 | 0.999 |
| NK → B | 44.0675 | 0.665 | 0.999 |
| CAF → LEC | -59.6736 | 0.667 | 0.999 |
| mast → B | -28.8112 | 0.670 | 0.999 |
| CAF → plasma | 47.0034 | 0.670 | 0.999 |
| CD4T → CAF | 19.3864 | 0.672 | 0.999 |
| macrophages → CAF | 21.8960 | 0.672 | 0.999 |
| NK → LEC | -38.4815 | 0.674 | 0.999 |
| CAF → CD4T | 19.1216 | 0.675 | 0.999 |
| B → granulocyte | 26.6799 | 0.675 | 0.999 |
| LEC → mast | -45.8904 | 0.677 | 0.999 |
| B → CD4T | 30.1202 | 0.678 | 0.999 |
| LEC → VSMC_myofibroblast | -38.2114 | 0.679 | 0.999 |
| FDC → TFH | -144.9371 | 0.682 | 0.999 |
| B → NK | 41.2332 | 0.684 | 0.999 |
| FDC → LEC | 81.5173 | 0.684 | 0.999 |
| CD4T → B | 29.1310 | 0.684 | 0.999 |
| NK → CAF | -46.5595 | 0.686 | 0.999 |
| granulocyte → FDC | 43.1844 | 0.686 | 0.999 |
| vasculature → gdT | 31.0108 | 0.687 | 0.999 |
| LEC → FDC | 80.6286 | 0.688 | 0.999 |
| plasma → CAF | 43.7727 | 0.689 | 0.999 |
| macrophages → gdT | 25.5335 | 0.690 | 0.999 |
| CD8T → NK | 41.8602 | 0.692 | 0.999 |
| macrophages → TFH | 28.7107 | 0.693 | 0.999 |
| FDC → granulocyte | 41.8842 | 0.693 | 0.999 |
| macrophages → CD4T | 12.3934 | 0.694 | 0.999 |
| gdT → B | -20.6575 | 0.695 | 0.999 |
| gdT → granulocyte | 34.7341 | 0.696 | 0.999 |
| granulocyte → gdT | 34.5618 | 0.697 | 0.999 |
| TFH → gdT | 48.4626 | 0.697 | 0.999 |
| TFH → FDC | -137.6888 | 0.700 | 0.999 |
| granulocyte → B | 24.1907 | 0.705 | 0.999 |
| gdT → TFH | 46.7562 | 0.708 | 0.999 |
| NK → TFH | 40.8753 | 0.708 | 0.999 |
| B → gdT | -19.7662 | 0.708 | 0.999 |
| DC → VSMC_myofibroblast | -26.2117 | 0.712 | 0.999 |
| LEC → NK | -33.5470 | 0.717 | 0.999 |
| CD4T → gdT | -21.2035 | 0.717 | 0.999 |
| stromal → CD8T | -8.0951 | 0.718 | 0.999 |
| CD4T → macrophages | 11.3810 | 0.719 | 0.999 |
| CD4T → stromal | 8.5922 | 0.719 | 0.999 |
| mast → LEC | -39.7905 | 0.720 | 0.999 |
| stromal → Treg | 8.1827 | 0.734 | 0.999 |
| stromal → CD4T | 8.1859 | 0.735 | 0.999 |
| CD8T → gdT | 24.9224 | 0.741 | 0.999 |
| gdT → vasculature | 25.3657 | 0.741 | 0.999 |
| TFH → NK | 35.6844 | 0.742 | 0.999 |
| gdT → CD4T | -19.2044 | 0.743 | 0.999 |
| CD4T → LEC | -13.8504 | 0.745 | 0.999 |
| VSMC_myofibroblast → DC | -22.9297 | 0.747 | 0.999 |
| gdT → CAF | 48.7062 | 0.747 | 0.999 |
| CAF → gdT | 48.1772 | 0.749 | 0.999 |
| NK → stromal | -14.9209 | 0.750 | 0.999 |
| gdT → CD8T | 23.9017 | 0.751 | 0.999 |
| NK → CD8T | 33.1365 | 0.754 | 0.999 |
| LEC → CD4T | -15.1146 | 0.756 | 0.999 |
| macrophages → macrophages | 14.8810 | 0.756 | 0.999 |
| plasma → vasculature | 28.9168 | 0.760 | 0.999 |
| vasculature → plasma | 28.4318 | 0.760 | 0.999 |
| gdT → macrophages | 19.3219 | 0.763 | 0.999 |
| CAF → VSMC_myofibroblast | 69.9550 | 0.765 | 0.999 |
| VSMC_myofibroblast → CAF | 69.2193 | 0.767 | 0.999 |
| CD8T → CAF | -12.9345 | 0.769 | 0.999 |
| TFH → mast | -28.4888 | 0.772 | 0.999 |
| mast → Treg | 17.4878 | 0.775 | 0.999 |
| NK → vasculature | -15.6699 | 0.775 | 0.999 |
| FDC → mast | -27.7509 | 0.781 | 0.999 |
| mast → TFH | -26.8570 | 0.784 | 0.999 |
| LEC → | -18.7912 | 0.784 | 0.999 |
| macrophages |  |  |  |
| vasculature → NK | -14.9825 | 0.786 | 0.999 |
| CD8T → TFH | 19.0252 | 0.787 | 0.999 |
| CAF → CD8T | -11.7430 | 0.790 | 0.999 |
| stromal → stromal | -7.8833 | 0.793 | 0.999 |
| FDC → CD8T | -15.1329 | 0.800 | 0.999 |
| Treg → LEC | -14.7647 | 0.800 | 0.999 |
| macrophages → | -16.8894 | 0.803 | 0.999 |
| LEC |  |  |  |
| CD8T → stromal | -5.4441 | 0.807 | 0.999 |
| CAF → FDC | -27.5498 | 0.815 | 0.999 |
| Treg → stromal | 5.4183 | 0.815 | 0.999 |
| LEC → Treg | -13.5770 | 0.816 | 0.999 |
| macrophages → DC | -8.2284 | 0.816 | 0.999 |
| plasma → | -12.7333 | 0.818 | 0.999 |
| macrophages |  |  |  |
| TFH → Treg | -15.7295 | 0.819 | 0.999 |
| Treg → TFH | -15.5606 | 0.820 | 0.999 |
| CD8T → | 7.4116 | 0.820 | 0.999 |
| macrophages |  |  |  |
| TFH → CD8T | 15.8841 | 0.821 | 0.999 |
| FDC → CAF | -26.6348 | 0.821 | 0.999 |
| TFH → | 16.4771 | 0.823 | 0.999 |
| macrophages |  |  |  |
| B → Treg | 16.1840 | 0.824 | 0.999 |
| CD8T → CD8T | 12.8362 | 0.825 | 0.999 |
| B → | 12.1224 | 0.831 | 0.999 |
| VSMC_myofibroblast |  |  |  |
| CD4T → CD4T | 15.6060 | 0.834 | 0.999 |
| CD8T → CD4T | 13.4390 | 0.835 | 0.999 |
| macrophages → | 6.7145 | 0.835 | 0.999 |
| CD8T |  |  |  |
| vasculature → B | -7.8920 | 0.839 | 0.999 |
| B → vasculature | -7.6993 | 0.840 | 0.999 |
| stromal → NK | -9.4740 | 0.841 | 0.999 |
| plasma → plasma | -28.7364 | 0.841 | 0.999 |
| Treg → B | 13.8852 | 0.847 | 0.999 |
| CD4T → CD8T | 12.3039 | 0.847 | 0.999 |
| plasma → CD8T | -8.8128 | 0.849 | 0.999 |
| LEC → LEC | 40.9005 | 0.850 | 0.999 |
| vasculature → | -22.4713 | 0.855 | 0.999 |
| vasculature |  |  |  |
| VSMC_myofibroblast | 10.1783 | 0.857 | 0.999 |
| → B |  |  |  |
| DC → macrophages | -6.0704 | 0.867 | 0.999 |
| NK → CD4T | 13.3152 | 0.869 | 0.999 |
| DC → gdT | -12.7295 | 0.875 | 0.999 |
| gdT → DC | -12.5286 | 0.877 | 0.999 |
| CAF → stromal | 10.6410 | 0.878 | 0.999 |
| stromal → TFH | 8.4954 | 0.880 | 0.999 |
| CD4T → NK | 12.0524 | 0.881 | 0.999 |
| FDC → B | -13.2489 | 0.885 | 0.999 |
| CD8T → FDC | -8.5725 | 0.886 | 0.999 |
| VSMC_myofibroblast | 10.2453 | 0.887 | 0.999 |
| → macrophages |  |  |  |
| VSMC_myofibroblast | 16.9758 | 0.889 | 0.999 |
| → granulocyte |  |  |  |
| macrophages → NK | 7.7766 | 0.890 | 0.999 |
| mast → FDC | -13.5278 | 0.891 | 0.999 |
| CD8T → plasma | -6.2351 | 0.892 | 0.999 |
| NK → macrophages | 7.5562 | 0.893 | 0.999 |
| B → FDC | -12.1192 | 0.893 | 0.999 |
| macrophages → | 9.6974 | 0.897 | 0.999 |
| VSMC_myofibroblast |  |  |  |
| B → plasma | 10.1475 | 0.903 | 0.999 |
| NK → plasma | -8.1674 | 0.906 | 0.999 |
| plasma → B | 9.8509 | 0.906 | 0.999 |
| stromal → CAF | 8.1459 | 0.907 | 0.999 |
| TFH → stromal | 6.6513 | 0.907 | 0.999 |
| vasculature → Treg | 3.9967 | 0.908 | 0.999 |
| granulocyte → | 14.0643 | 0.909 | 0.999 |
| VSMC_myofibroblast |  |  |  |
| LEC → B | -6.8785 | 0.911 | 0.999 |
| Treg → Treg | -7.9585 | 0.915 | 0.999 |
| NK → mast | 12.4154 | 0.916 | 0.999 |
| plasma → NK | -7.2072 | 0.916 | 0.999 |
| plasma → mast | 8.2526 | 0.927 | 0.999 |
| mast → NK | 10.2423 | 0.930 | 0.999 |
| DC → Treg | 4.8876 | 0.933 | 0.999 |
| CAF → TFH | 10.5095 | 0.934 | 0.999 |
| TFH → CAF | 9.8964 | 0.938 | 0.999 |
| Treg → CD8T | -4.3065 | 0.942 | 0.999 |
| Treg → DC | 4.0842 | 0.944 | 0.999 |
| CAF → CAF | -16.4188 | 0.948 | 0.999 |
| B → LEC | -3.7053 | 0.952 | 0.999 |
| Treg → mast | 3.5859 | 0.953 | 0.999 |
| mast → plasma | 4.9601 | 0.956 | 0.999 |
| CD4T → Treg | 2.9813 | 0.960 | 0.999 |
| granulocyte → | -4.0600 | 0.961 | 0.999 |
| plasma |  |  |  |
| Treg → vasculature | 1.2928 | 0.968 | 0.999 |
| CD8T → Treg | -2.2597 | 0.969 | 0.999 |
| gdT → FDC | -3.4178 | 0.973 | 0.999 |
| TFH → plasma | -2.7164 | 0.975 | 0.999 |
| FDC → gdT | -2.8637 | 0.977 | 0.999 |
| plasma → TFH | 2.4540 | 0.977 | 0.999 |
| macrophages → | 1.2579 | 0.978 | 0.999 |
| plasma |  |  |  |
| FDC → FDC | 10.0892 | 0.980 | 0.999 |
| stromal → FDC | -0.9935 | 0.990 | 0.999 |
| Treg → CD4T | 0.7661 | 0.990 | 0.999 |
| CAF → Treg | 0.8455 | 0.990 | 0.999 |
| FDC → stromal | 0.6394 | 0.993 | 0.999 |
| plasma → granulocyte | -0.3898 | 0.996 | 0.999 |
| Treg → CAF | 0.0605 | 0.999 | 0.999 |

#### S9.4 HNSCC *p*-value distribution comparison

**Figure S9:**
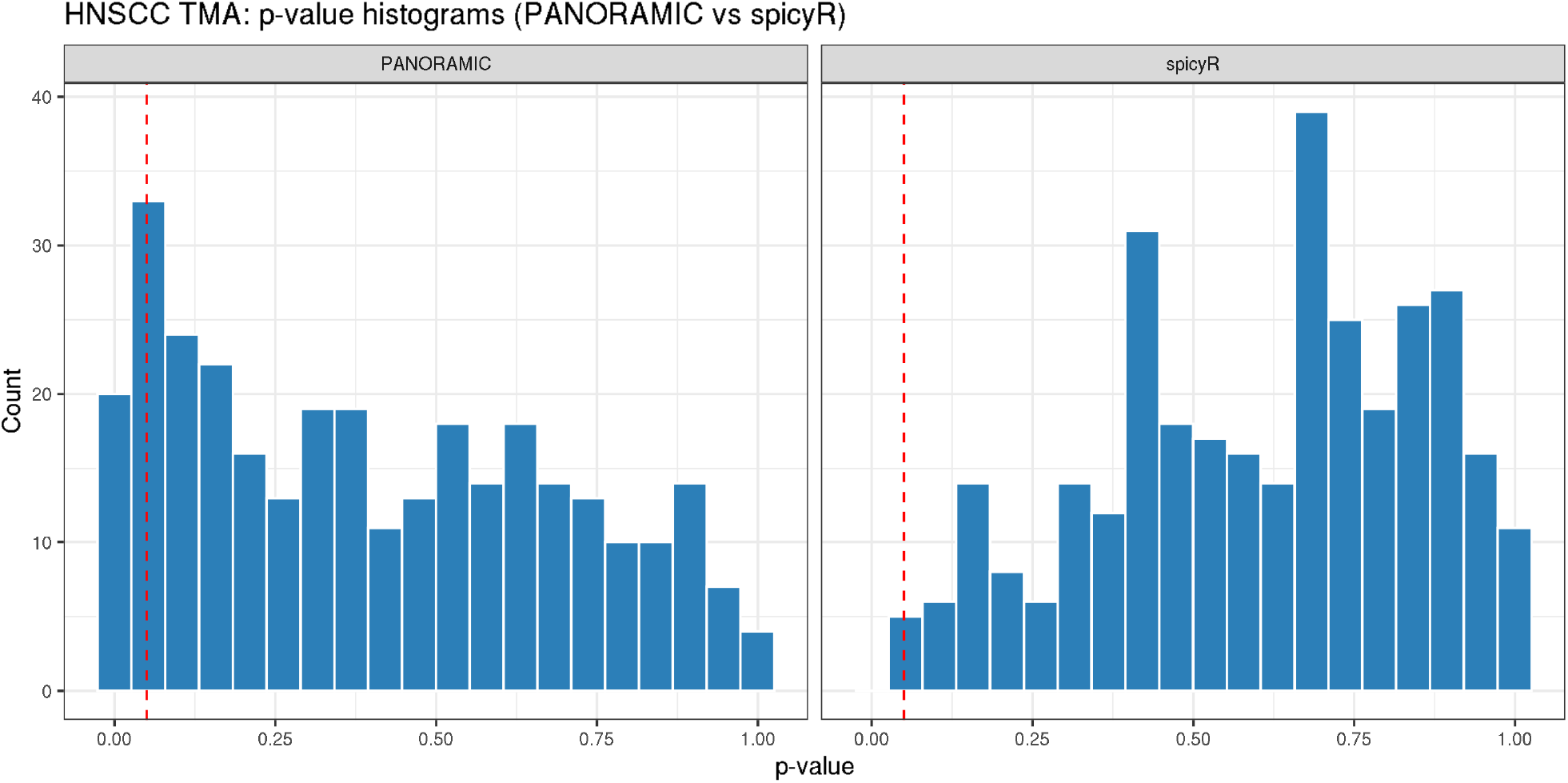
P-value distributions for the HNSCC TMA differential colocalization analysis comparing PANORAMIC and spicyR. Histograms show nominal p-values across tested directional cell-type pairs for the benign lymph node versus tumor-adjacent lymph node comparison. The red dashed vertical line marks *p* = 0.05. PANORAMIC shows enrichment of small p-values with a broad background across the remaining range, whereas spicyR produces fewer small p-values and a concentration of larger p-values.

